# Hierarchical Transformation of Navigational Variables into Coordinated Turning

**DOI:** 10.1101/2024.06.27.601106

**Authors:** Kai Feng, Corinna Gebehart, Romita Trehan, Virginia Palieri, Yasha Zaman, Karin A. van Hassel, Mariam Khan, Ryo Minegishi, Annika Müller, Matthew N. Van De Poll, Bruno van Swinderen, Katherine I. Nagel, M. Eugenia Chiappe, Hannah Haberkern, Barry J. Dickson

## Abstract

The transformation of navigational variables into coordinated motor actions is a fundamental requirement for autonomous locomotion. The circuit and control mechanisms that underlie this transformation are not well understood. Here, we combine connectomics, neuronal activity perturbations, functional imaging and electrophysiology to identify a central steering circuit in *Drosophila*. This steering circuit operates in both goal-directed navigation and exploratory search and controls turning during both walking and flight, by implementing a hierarchical, stepwise transformation from navigation-related signals to motor output. At the top of the hierarchy are the reciprocally connected DNa03 and LAL013 neurons. DNa03 integrates lateralized sensory and internal information to generate an effector-agnostic steering signal during goal-directed navigation. LAL013 promotes exploratory turning in response to bilateral input from ExR7 neurons of the central complex, which encode the uncertainty of the fly’s internal heading estimate. Both DNa03 and LAL013 provide input to a set of hierarchically organized descending neurons, at the top of which is DNa11. DNa11 targets leg motor circuits directly as well as indirectly through subordinate descending neurons, and its activation coordinately changes the stepping directions of all six legs to generate rapid saccadic turns. In contrast to DNa03, DNa11 shows unilateral, delayed and low pass filtered activity during a turn. The signal transformation between DNa03 and DNa11 thus marks the transition from motor planning to the execution of a walking turn. Together, our results reveal a hierarchical circuit architecture that transforms navigational variables into abstract steering representations and ultimately into motor commands, providing a general framework for flexible control of behaviour across contexts and motor systems.

## Introduction

To navigate through its environment, an autonomous agent requires a control system capable of transforming spatial representations into coordinated steering movements. Such control systems face several computational challenges. At any moment, they may be confronted with many different incentives to turn, varying in their degrees of certainty and often conflicting with one another. These incentives may arise from either external sensory cues or internal goal or motivational signals. The control system must therefore be able to integrate multiple inputs, weigh the confidence and significance of each, and produce a single steering output: a decision whether to turn or not, and if so, in which direction and by what amplitude. Executing a turn requires translating this generic steering instruction into precise and coordinated movements of the available effectors, whether they be legs, wings, fins or wheels.

Engineered steering control systems have approached these problems through centralized^1^or distributed architectures^2^. Biological systems, by contrast, appear to be more distributed in nature. In zebrafish larvae, for example, the rapid turns made during prey pursuit are controlled by brain circuits distinct from those used for exploratory turns^3^. Evidence for a distributed steering control system has also emerged in *Drosophila,* for which it has been shown that natural turns are frequent and variable^4^, that turning-related neural activity is widespread in the brain^5-7^ and that turning can be elicited by multiple distinct classes of descending neurons^8-11^ – the neurons that project from the brain to motor circuits in the ventral nerve cord (VNC). Here, we present evidence for a central steering mechanism in *Drosophila*, supporting a more unified view of turning control.

In *Drosophila* and other insects, external cues, such as visual targets or odour plumes, trigger turns by asymmetric activating sensory neurons^8,12-16^. This lateralized input leads to asymmetric activity in the lateral accessory lobe (LAL), a pre-motor region that receives high-order input from all sensory modalities and generates descending steering commands^17-20^. Recent research has also begun to reveal how turns are generated from internally stored variables. During goal-directed spatial navigation, neurons in the central complex track the fly’s current^21^ and goal headings^22^ and compute the difference between the two^22,23^. The resulting difference vector is sent via the PFL2 and PFL3 neurons to the LAL^22,23^. How the PFL2 and PFL3 outputs are translated into a corrective turn of the appropriate magnitude and direction is unknown. The influence of PFL3 neurons on turning behaviour is probabilistic^22,24^, indicating that steering decisions in the LAL are likely made by integrating the central complex output with external sensory signals and possibly also stochastic factors (Fig. 1a). In moths and ants, exploratory turns are also initiated in the LAL^19,25^. Male silk moths use pheromones to locate a mate^26^ and ants use visual cues to locate their nest^27^. When these sensory cues are lost, silk moths and ants make frequent turns and counterturns in search of the lost cues^26,28^. Flies similarly initiate alternating turns upon the offset of an attractive odour^29-31^. Such exploratory turns may represent an adaptive strategy for reducing uncertainty in sensory or internal navigational representations. External sensory information, course correction signals, and exploratory drives thus all converge upon the LAL. If and how the LAL integrates these various incentives, contends with uncertainty, and makes steering decisions is unknown.

**Fig. 1:**
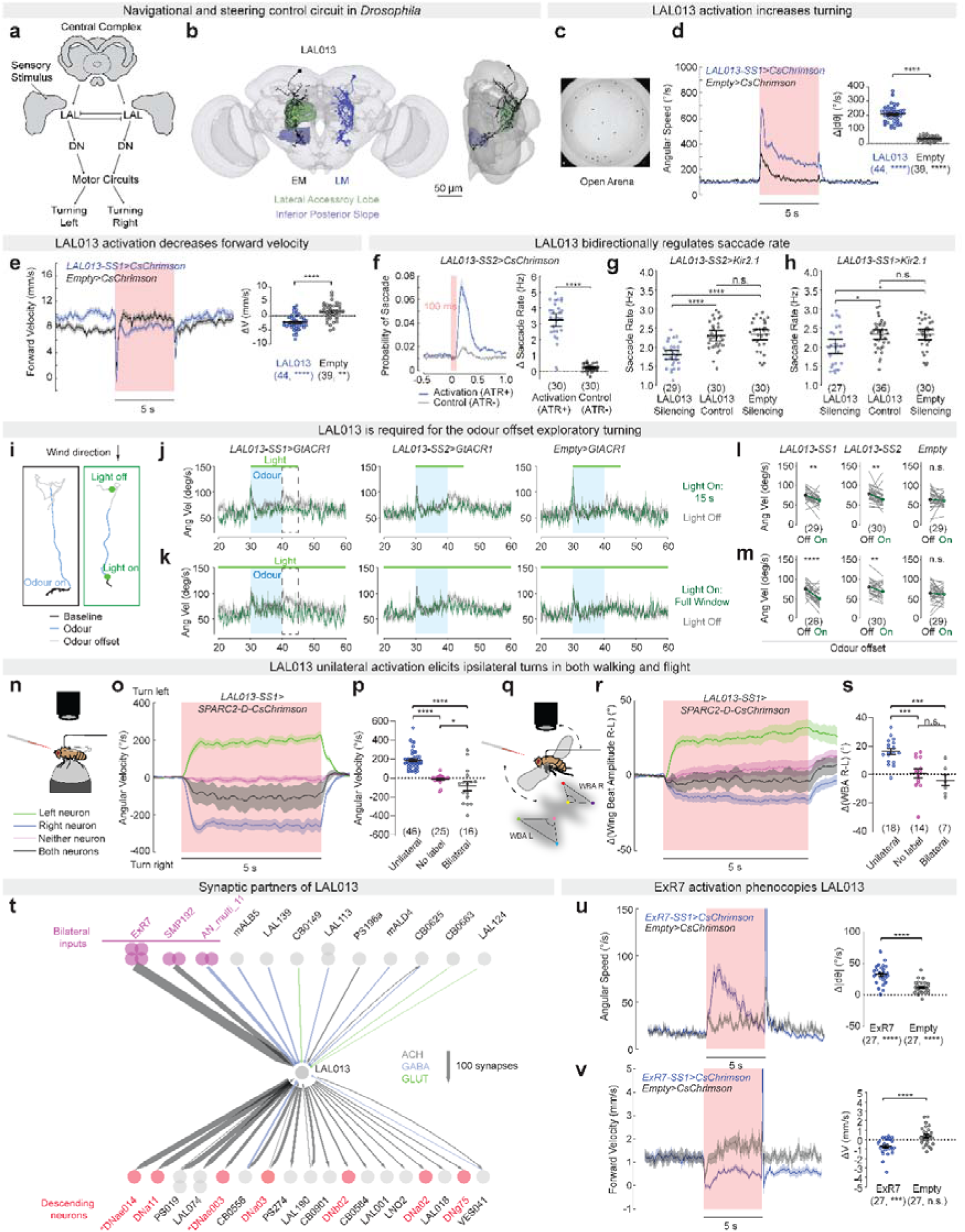
LAL013 controls exploratory turning. **a**, Schematic of the *Drosophila* steering circuit. LAL, lateral accessory lobe; DN, descending neurons. **b**, Frontal (left) and side (right) views of a fly brain with LAL013 neurons labelled. Black: the LAL013 neuron from the electron microscopy (EM: the hemibrain) volume; blue: a segmented neuron matching LAL013 from a light microscopy (LM) image of *VT002063-GAL4*. **c**, Example frame of the “Fly Bowl” assay. **d**, Left, average angular speed across 9 trials of before, during and after 5-s red-light stimuli (pink rectangle). Right, quantification of the data in (d) as single fly’s change in angular speed. **e**, Average forward velocity quantified as in (d). **f**, Time course of the probability of saccade before and following a 100-ms red optogenetic stimulus for *LAL013-SS2>CsChrimson* flies fed with all-trans retinal (Activation) or regular food (Control). **g, h**, Saccade rates of free walking flies. Experiments in (d-e) were performed in Fly Bowl, while (f-g) were performed in a different open arena. See Methods for details. **i**, Fly’s walking trajectories from a miniature wind-tunnel paradigm in sessions with or without optogenetic stimulation. Both panels are from a *LAL013-SS1>GtACR1* fly. **j**, Angular velocity of LAL013 silencing and control flies during experimental sessions (15-s light, green) and control sessions (light off, grey) before, during and after odour delivery (blue shade). **k**, Same as (j) but with light on throughout the experimental sessions. Dashed box indicates the odour offset period. **l, m**, Quantification of each fly’s average angular velocity during the odour-offset period of experimental and control sessions. **n**, Schematic of the spherical treadmill walking assay with optogenetic stimuli. **o**, Average angular velocity across 10 trials per fly. **p**, Quantification of the data in (o) with left neuron and right neuron groups merged as a single “Unilateral” group. Each data point represents the average angular velocity of a single fly across 10 trials during the first 2 seconds after red-light onset. **q**, Schematic of the flight assay with optogenetic stimuli. The difference in wing beat amplitude (WBA) between the right and left wings was used as a proxy of angular velocity. **r**, Average change of the difference in WBA across 10 trials per fly, plotted as in (o). **s**, Quantification of the data in (r), plotted as in (p). **t**, Top upstream and downstream synaptic partners of LAL013 grouped by types and ranked by synapse counts. Line widths indicate synapse numbers, relative to reference widths for 100 synapses. Asterisks, see Methods for nomenclature. **u, v**, Average angular speed (u) and forward velocity (v) of *ExR7>CsChrimson* and control flies, shown as in (d-e). Data with error bars or shades are mean ±Ls.e.m. across flies. Numbers in brackets are n of flies. ****, *p*<0.0001; ***, *p*<0.001; **, *p*<0.01; *, *p*<0.05, n.s., *p*>0.05; unpaired t-test (d-e, u-v), Mann–Whitney U test with Bonferroni corrections (f-h), paired t-test (l-m) and Holm-Sidak’s test (p, s) for comparisons between groups; one sample t-test for comparisons to zero (d-e, u-v, asterisks in brackets).

Amongst the LAL outputs, the DNa02, DNae014, DNb01 and DNg13 descending neurons have all recently been shown to function in turning^8-10^. As turning-related activity is wide-spread among descending axons^5^, these neurons likely represent a small fraction of descending pathways that participate in steering control. Moreover, activation of the walking neurons DNa02 and DNg13 elicits rather subtle turns^9^ which do not resemble the large-amplitude saccadic turns often seen in freely walking flies^32-34^. The overall organization of the descending pathways for turning thus remains unclear. In particular, do all the descending neurons involved in turning execute the low-level motor control typified by DNa02 and DNg13^9^, or might some exert a higher level of control in coordinating the many independent changes in motor function that underlie a graceful turn^35^?

In this study, we identify the reciprocally connected DNa03 and LAL013 neurons as a central steering control hub in the *Drosophila* LAL. These neurons receive both directional and non-directional navigational signals from the central complex and communicate with both leg and wing motor circuits through a suite of hierarchically connected descending neurons. We investigate how this hierarchical steering-control circuit progressively transforms navigation variables into coordinated turning actions across behavioural contexts and motor systems.

## Results

### LAL013 controls exploratory turning

In an optogenetic screen for locomotion phenotypes, we identified a GAL4 driver line, *VT002063* (ref^36^), that elicited robust turning upon activation with CsChrimson^37^. *VT002063>CsChrimson* flies turned persistently during a stimulus period of several seconds. The *VT002063* driver is broadly expressed in the nervous system^36^. Through a series of stochastic activation experiments we were able to assign the turning response to the activation of a pair of unilateral neurons, one in each brain hemisphere (Fig. 1b and data not shown). We obtained two stable split-GAL4 driver lines (SS lines), which label these neurons sparsely (ref^38^ and Fig. S1), and subsequently used the SS lines rather than *VT002063* for the anatomical and functional characterization of these neurons.

The neurons labelled by the SS lines primarily innervate the LAL and the inferior posterior slope (IPS). Both brain regions show turning-related neural activity^6,39^ and are richly innervated by descending neurons that project to leg and wing motor circuits in the VNC^40^. The closest match to these neurons in the connectomes^41-43^ are the LAL013 neurons, the name by which we subsequently refer to them (Fig. 1b).

In freely-walking flies in an open arena (Fig. 1c), bilateral LAL013 activation elicited robust turning with a decrease in forward velocity (Fig. 1d-e), a pattern resembling body saccades. In natural settings, flies often utilize a fixation-saccade locomotion strategy, consisting of forward runs used to stabilize visual gaze, interrupted by rapid body saccades used for quick reorientation^34^. By applying established criteria based on unsupervised clustering of natural walking kinematics to detect saccades^34^, we found that a 100-ms optogenetic stimulation of LAL013 elicited turns that are indistinguishable from natural saccades (Fig. 1f), whereas silencing LAL013 reduced the saccade rate in free walking flies (Fig. 1g-h).

The turning direction during LAL013 bilateral activation appeared to be random, with a single fly switching from one direction to the other between trials and occasionally even during a 5-s trial (Fig. S2, Supplementary Video 1). This activation phenotype resembles the searching behaviours of silk moths and other insects^19^, including *Drosophila*^29,30^. Upon losing track of an attractive odour plume, *Drosophila* exhibit search behaviour characterized by reduced ground speed, increased angular velocity, and longer history dependence in angular velocity^30^. The locomotor pattern evoked by bilateral LAL013 activation showed all these features of search behaviour (Fig. S2). We therefore asked whether LAL013 is required for such exploratory turns. Indeed, we found that acute silencing of LAL013 significantly reduced the increase in turning upon odour offset (Fig. 1i-m). These results are consistent with the hypothesis that LAL013 is recruited to increase turning in settings where flies need to maximize their environment sampling.

To further characterize the turning induced by LAL013 neurons, we used the SPARC system^44^ to stochastically activate single neurons in flies walking on an air-suspended ball (Fig. 1n). Each fly was subject to multiple activation trials, after which its brain was dissected and imaged to determine whether the left, right, both, or neither LAL013 neuron was labelled. Flies with unilateral LAL013 labelling consistently turned towards the ipsilateral side (Fig. 1o-p). By contrast, those with bilateral LAL013 labelling turned stochastically in either direction (Fig. 1o-p). Flies with no LAL013 neurons labelled showed very little turning (Fig. 1o-p). A similar pattern was observed for turns executed during flight (Fig. 1q-s).

The connectomes^41-43^ suggest that LAL013 receives its strongest synaptic input from ExR7, an output channel of the central complex^45^, and SMP192, which arborizes in a brain region involved in high order sensory processing (the superior medial protocerebrum^46,47^) (Fig. 1t). Neither of these neuronal cell types has been functionally characterized, but this connectivity suggests that LAL013 neurons integrate both internal and external information to generate a turning signal. Interestingly, however, all the ExR7 and SMP192 cells connect to both the left and right LAL013 neurons, implying that these inputs alone are unlikely to account for any unilateral LAL013 activity that might occur during naturally evoked turns. Instead, they may modulate LAL013 activity bilaterally to promote exploratory behaviour. Consistent with this idea, we found bilateral ExR7 activation elicits a similar, albeit milder, phenotype as LAL013 activation, characterized by increased turning and decreased forward locomotion (Fig. 1u-v).

The ability of bilateral optogenetic activation of LAL013 to induce robust unilateral turning suggests that a “winner-take-all” mechanism operates between the two LAL013 neurons.

### LAL013 interfaces with a hierarchical network of descending neurons

LAL013 is presynaptic to several classes of descending neuron (Fig. 1t). These descending outputs include DNae014, which has been shown to control saccades made during flight^10^, and DNa02, which contributes to turns made during walking^8,9^. Three other functionally uncharacterized descending neurons also receive strong direct input from LAL013: DNa11, DNae003, and DNa03 (Fig. 1t and Fig. 2a).

**Fig. 2:**
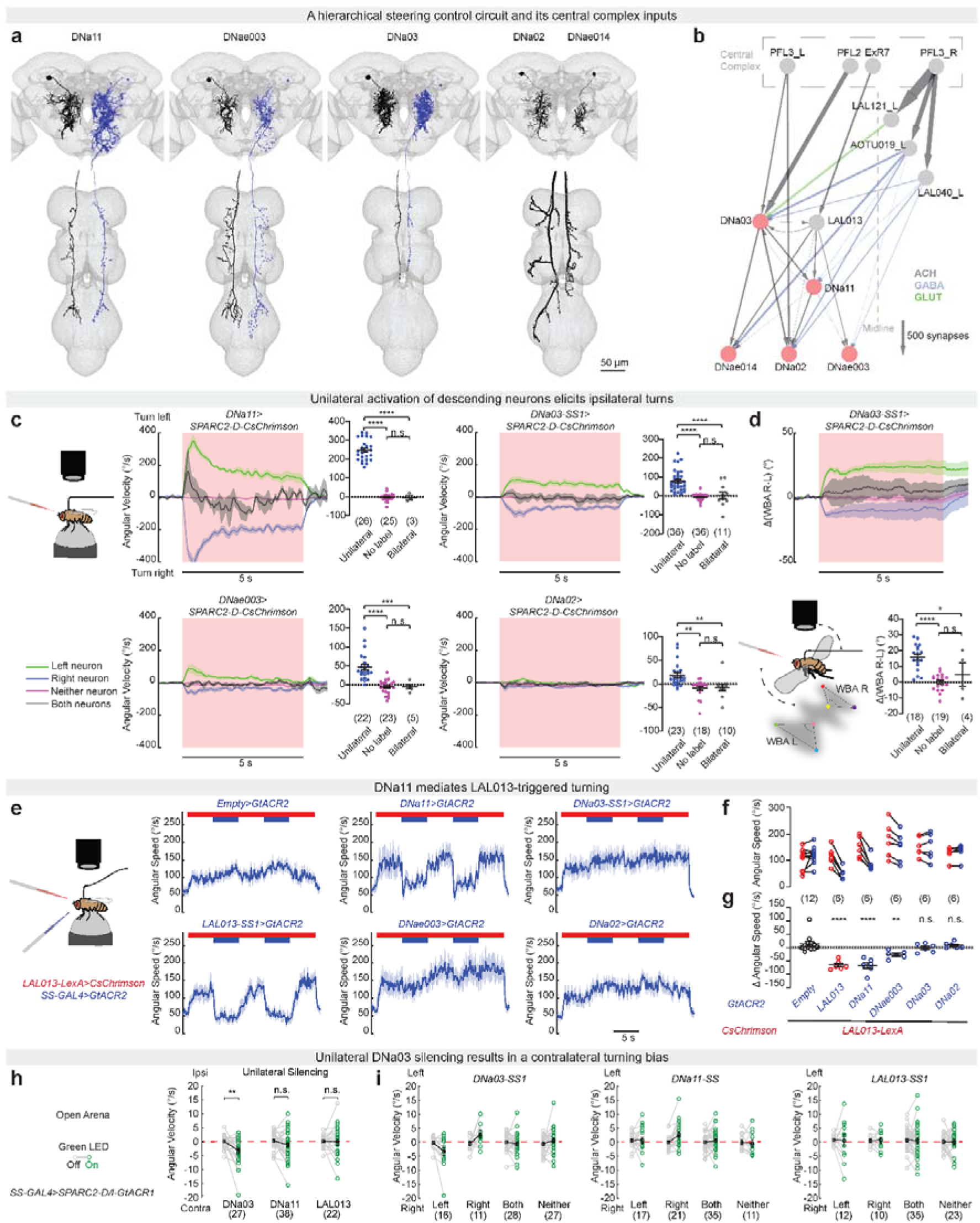
LAL013 interfaces with a hierarchical network of descending neurons. **a**, Fly brain and VNC images with descending neurons labelled. Black: neurons from EM (the hemibrain and MANC); blue: segmented neurons from LM. **b**, Synaptic connections between LAL013, selected descending neurons and central complex output neurons. Line widths indicate synapse numbers. **c**, Stochastic activation of descending neurons in flies walking on a spherical treadmill. **d**, Stochastic activation of DNa03 in tethered flight. **e**, Left, schematic of neuronal epistasis experiments. Right, angular speed per fly, averaged across 10 trials per fly. Red and blue bars indicate red- and blue-light stimuli, respectively. **f**, Average speed during periods of red light only (red circles) and periods of simultaneous red and blue light (blue circles) for each fly. **g**, The angular speed change between the two periods in (f) for each fly. **h**, Average angular velocities of flies unilaterally expressing GtACR1 in DNa03, DNa11 or LAL013 and walking spontaneously in Fly Bowl during green-light-off (grey) and -on (green) periods. Black squares indicate mean across flies. **i**, Four cohorts of flies with the left, right, both or neither neuron labelled for each cell type are shown separately. Data with error bars or shades are mean ±Ls.e.m. across flies. Numbers in brackets are n of flies. ****, *p*<0.0001; ***, *p*<0.001; **, *p*<0.01; *, *p*<0.05, n.s., *p*>0.05; Holm-Sidak’s test (c, d) for comparisons between groups; Dunnett’s test (g) for comparisons between each experimental group and the control; paired t-test for comparisons between light on and off (h).

LAL013 and its output descending neurons are organized in a hierarchical network. LAL013 and DNa03 comprise the top layer, forming connections with each other as well as all four of the other descending neurons (Fig. 2b). Although DNa03 projects to the VNC, the majority of its output synapses are in the brain. DNa11 represents an intermediate layer, receiving input from LAL013 and DNa03. DNa11 innervates the leg neuropils in the VNC but also has output synapses with other descending neurons in the brain. DNa11 provides strong output to DNa02 and weaker output to DNae003 and DNae014. DNa02, DNae003, and DNae014 form the bottom layer of the network, with relatively little output to other neurons in the brain (Fig. 2b). The output synapses of DNa02 and DNae003 are primarily located in the leg neuropils; the outputs of DNae014 are in the wing neuropils.

We generated genetic driver lines specific for each of the previously uncharacterized descending neurons (Fig. S1) and obtained existing reagents for DNa02 (ref^40^). Using these lines, we first performed stochastic activation experiments analogous to those above for LAL013 (Fig. 1n-s). We found that all four descending neurons elicited ipsilateral turns during walking, albeit with widely varying potency (Fig. 2c and Fig. S3). DNa11 reliably evoked turns of high angular velocity (>200°/s), resembling those elicited by LAL013 activation (Fig. 2c and Fig. S3). DNa03 and DNae003 triggered turns of progressively smaller amplitude and with lower penetrance (Fig. 2c and Fig. S3). Turns induced by DNa02 activation were weakest, typically below 50°/s and highly probabilistic, consistent with previous reports^8,9^ (Fig. 2c and Fig. S3). Like LAL013, DNa03 is also presynaptic to DNae014, which targets wing motor circuits and elicits turns during flight^10^. We found that, as for LAL013 (Fig. 1q-s) and DNae014^10^, unilateral DNa03 activation also elicited turns during flight (Fig. 2d and Supplementary Video 2).

We next asked to what extent each of these descending neurons contributes to LAL013-triggered turns during walking. For this, we generated a *LAL013-LexA* driver line and combined it with *LexAop-CsChrimson* to produce flies that turned persistently during continuous red-light stimulation (Fig. 2e). This allowed us to use the GAL4 system to express the optogenetic anion channel GtACR2^48^ in selected descending neurons so that they could be acutely silenced with intermittent pulses of blue light, revealing whether their activity is required for LAL013-triggered turning. We found that silencing DNa11 blocked LAL013-triggered turning just as effectively as the positive control of silencing LAL013 itself (Fig. 2e-g). Silencing DNae003 had a modest but significant effect, whereas the DNa02 and DNa03 drivers, like the negative control GAL4 driver, were ineffective in blocking LAL013-triggered turns. We conclude that LAL013 drives ipsilateral turning primarily through DNa11 and independently of DNa03.

To assess the contribution of these neurons to spontaneous turns in a more natural setting, we used GtACR1^48^ to silence LAL013, DNa03 and DNa11 stochastically in flies walking in an open arena. We found unilateral silencing of DNa03 biased angular velocity towards the contralateral side (Fig. 2h-i). By contrast, unilateral silencing of LAL013 or DNa11 failed to bias angular velocity (Fig. 2h-i), as previously observed also for DNa02 (ref^9^). These results, together with the connectivity data, suggest that DNa03 has a central function in steering control, especially in setting the directional bias, and that it can induce turns through any of several effector pathways.

### The central complex provides both directional and non-directional inputs to the steering circuit

Connectome analysis revealed that the steering circuit receives substantial input from the central complex, most notably from the columnar PFL2/PFL3 cells and from the ExR7 cells (Fig. 2b). PFL3 neurons are of particular interest because they convey directional error signals during goal-directed navigation^22,23^. DNa03 and DNa02 receive some direct synaptic input from PFL3. However, they are even more strongly connected to PFL3 indirectly through several LAL and anterior optic tubercle (AOTU) neuron types (Fig. 2b, also see Figure 63 in ref^45^), and computational analysis suggests that goal-directed navigation relies primarily on these indirect pathways from PFL3 to DNa03 and DNa02^49^. The most prominent indirect pathway is via the bilateral pair of LAL121 neurons. LAL121 is the strongest downstream synaptic partner of PFL3, receiving 13% of PFL3 output^42,43^ (about 10-fold more than any descending neuron). Conversely PFL3 accounts for 45% of the total synaptic inputs received by LAL121^42,43^ (Fig. 3a). Each LAL121 neuron integrates the output of PFL3 neurons to one brain hemisphere and projects to the contralateral hemisphere, where it provides strong glutamatergic (likely inhibitory^50^) input to DNa03 (14% of LAL121 output synapses^42,43^). The ExR7 cells are another prominent output channel of the highly recurrent central complex^45^, the function of which is less well understood. Outside the central complex, the principal output target of the four isomorphic and reciprocally connected ExR7 cells is LAL013 (Fig. 2b and Fig. 3a).

**Fig. 3:**
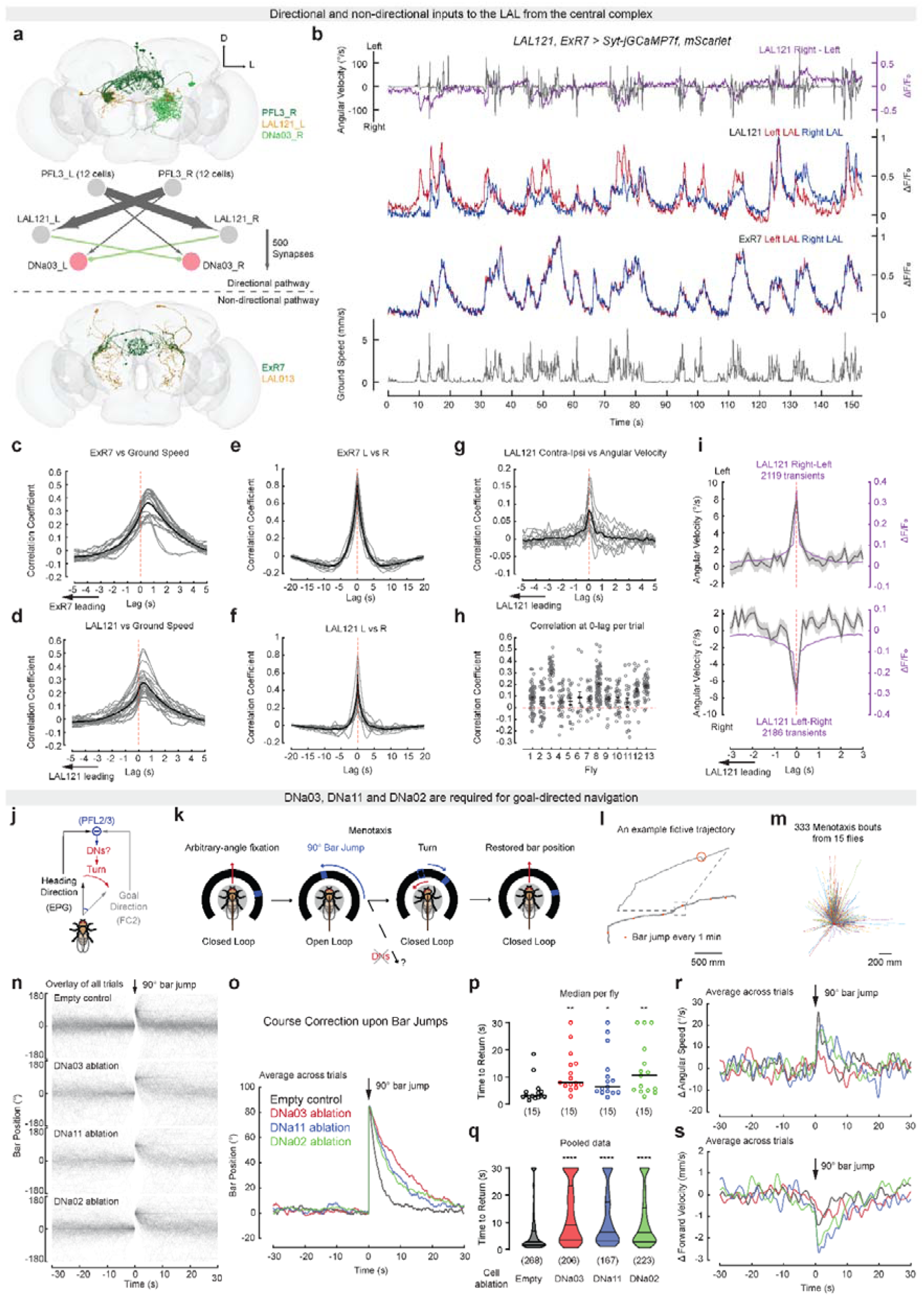
DNa03, DNa11 and DNa02 are required for goal-directed navigation. **a**, Overlay of PFL3, LAL121 and DNa03 in an EM image (top) and their connectivity pattern (middle). Overlay of ExR7 and LAL013 (bottom). Images and connection weights are from FlyWire. **b**, Example traces from LAL121 and ExR7 dual imaging experiments. **c**, Cross-correlation between ExR7 and ground speed. Each grey line represents one fly and the thick black line represents the mean across flies (n = 13 flies). **d**, Same as (c) for LAL121. **e**, Cross-correlation between ExR7 left and right neuron activity. **f**, Same as (e) for LAL121. **g**, Cross-correlation between contralateral-minus-ipsilateral activity of LAL121 and angular velocity. **h**, Quantification of the 0-lag correlation coefficient for each 22-s trial from 13 flies, lines and error bars indicate mean ±Ls.e.m. across trials for each fly. **i**, Bilateral differences in LAL121 activity and contraversive angular velocity aligned and averaged by the peak of LAL121 transients, for both directions. Data are mean ±Ls.e.m. across transients. **j**, Schematic of the circuit mechanisms for goal-directed navigation in *Drosophila*^22,23^. **k**, Schematic of the menotaxis assay. **l**, An example fictive trajectory of a control fly, with red dots indicating the locations where the bar jumps at 1-minute intervals occurred. **m**, Fictive trajectories of 333 menotaxis bouts from 15 control flies. Red dot indicates fly’s starting point. The bar was fixed at north (top on graph) to the fly’s starting point to compute the fictive trajectories. **n**, The trajectories of bar position in all menotaxis trials aligned around the 90° bar jumps and overlaid. The bar position in the frame immediately before the jump (time 0 s) in each trial is designated as 0°. **o**, Bar position data in (n) averaged (circular mean) for each genotype. **p**, Time until the bar returns to within 30° from the pre-jump position (no return within 30 s counted as 30 s). Each data point is the median from all trials for each fly. Numbers in brackets are n of flies. Lines indicate the medians across flies. **q**, Violin plot of the bar return times pooled for all menotaxis trials across flies for each genotype. Numbers in brackets are n of trials. The thick lines indicate medians, and the thin lines indicate quartiles. **r**, Angular speed change and **s**, forward velocity change around 90° bar jumps, averaged across all menotaxis trials and flies for each genotype. ****, *p*<0.0001; **, *p*<0.01; *, *p*<0.05; Dunn’s test (p, q) for comparisons between DN-ablated flies and the control.

To investigate how these two major central complex output channels influence the steering circuits, we imaged the activity of LAL121 and ExR7 neurons in flies walking spontaneously on a spherical treadmill. We combined SS lines for LAL121 and ExR7 (Fig. S1) to drive synaptotagmin-jGCaMP7f expression in both cell types, allowing us to simultaneously image calcium activity at output sites of both cell types in the same flies. We found that both cell types are strongly modulated by locomotor activity (Fig. 3b-d). The cross-correlations between both LAL121 and ExR7 activity and ground speed have wide peaks, suggesting that they not only influence but also receive feedback from ongoing locomotor behaviours.

We observed substantial bilateral differences in LAL121 but not ExR7 (Fig. 3b,e,f), suggesting that the PFL3 pathway provides directional information to the LAL steering circuits but ExR7 does not. This conclusion is consistent with the pronounced laterality of the PFL3-LAL121 pathway, in contrast to the bilateral symmetry of ExR7 connections. Although the left and right LAL121 also had highly correlated activity, their difference shows contraversive correlation with angular velocity (Fig. 3b,g-i), consistent with LAL121 providing inhibitory inputs to DNa03 and other steering neurons. However, the correlation between bilateral difference of LAL121 and angular velocity had a wide range of distribution among flies and time windows (Fig. 3g-i), consistent with the idea that the goal-heading error signal provided by PFL3 acts as an incentive to turn and is only probabilistically transformed into action. Under self-paced walking in darkness, the goal or heading signals may often be weak or poorly defined, such that turns are often triggered by competing drives.

### DNa03, DNa11 and DNa02 are required for goal-directed navigation

During goal-directed navigation, the central complex tracks the fly’s current and goal headings. By calculating the difference between the two, it triggers any turns needed to correct a deviation from the intended path. PFL2 and PFL3 initiate these course corrections (Fig. 3j). The connectome and our behavioural data suggest that these corrective turns are executed by activating DNa03 and, during walking, its subordinate descending neurons DNa11 and DNa02. We performed menotaxis assays^22,23,51^ to test this hypothesis.

During menotaxis, flies maintain a constant heading at some arbitrary angle with respect to a visual cue. This behaviour resembles the long-distance translocation of flies in the wild using cues such as the sun as a visual landmark^52^. In the laboratory paradigm with tethered flies, the position of the visual cue is controlled in closed loop so that the visual stimulus rotates in the opposite direction whenever the fly turns on the ball, thus providing the visual illusion of a change in heading. If the closed loop is briefly broken and the visual cue artificially rotated by 90°, then the fly typically performs a corrective turn, realigning its heading to the visual cue as soon as the loop is closed (Fig. 3k). Activities of both the PFL2 and PFL3 neurons contribute to these corrective turns^22,23^.

We used such a set up to test whether DNa03, DNa11, and DNa02 also function in goal-directed navigation. All control and experimental flies reliably performed menotaxis in our assay (Fig. 3l-m and Fig. S4), whereby, we defined menotaxis as a bout of continuous straight trajectory of at least 200 mm with no more than 25-mm deviation^22^ (Fig. 3m and Fig. S4g). To assess a fly’s ability to correct its heading after the bar jumps, we aligned the bar positions around each of the jumps that the fly experienced during a menotaxis bout. Control flies typically returned the bar to its original position within a few seconds (Fig. 3n-o). In DNa03, DNa11, or DNa02-ablated flies (Fig. S4a-f), however, the bar returned more slowly and less reliably (Fig. 3n-o). To quantify these differences, we calculated the median time for each fly to return the bar to within 30° of its original position. These median return times were significantly slower for each of the DN-ablated genotypes than for control flies (Fig. 3p-q), with several of the DN-ablated flies but none of the control flies failing to return the bar position within 20s of a jump. Consistent with previous reports^22,23,51^, all genotypes showed increased angular speed and decreased forward velocity during the corrective turns (Fig. 3r-s), indicating that the navigation system is still sensitive to course deviation in the DN-ablated flies. Together, these results show that DNa03, DNa11 and DNa02 are all required for rapid course corrections during goal-directed navigation.

### ExR7 promotes exploratory turns via LAL013 when the compass is less informative

Like the directional PFL2/3 pathway, the non-directional ExR7 pathway is also heavily influenced by the head direction system. In *Drosophila*, the head direction is represented as an activity bump in the population of EPG neurons, each of which occupies a wedge of the ring-like ellipsoid body (EB)^21^. ExR7 receives its strongest synaptic inputs from EPG neurons (Fig. 4a, 32% of ExR7’s total synaptic inputs^42,43^), with each ExR7 cell arborizing around the entire EB ring and contacting the full complement of EPG neurons (Fig. 4b). To gain insight into how the ExR7 cells might transform the heading signal encoded in EPG neurons into a useful input to the LAL steering circuit, we performed simultaneous imaging of ExR7 and EPG neurons (Fig. 4b).

**Fig. 4:**
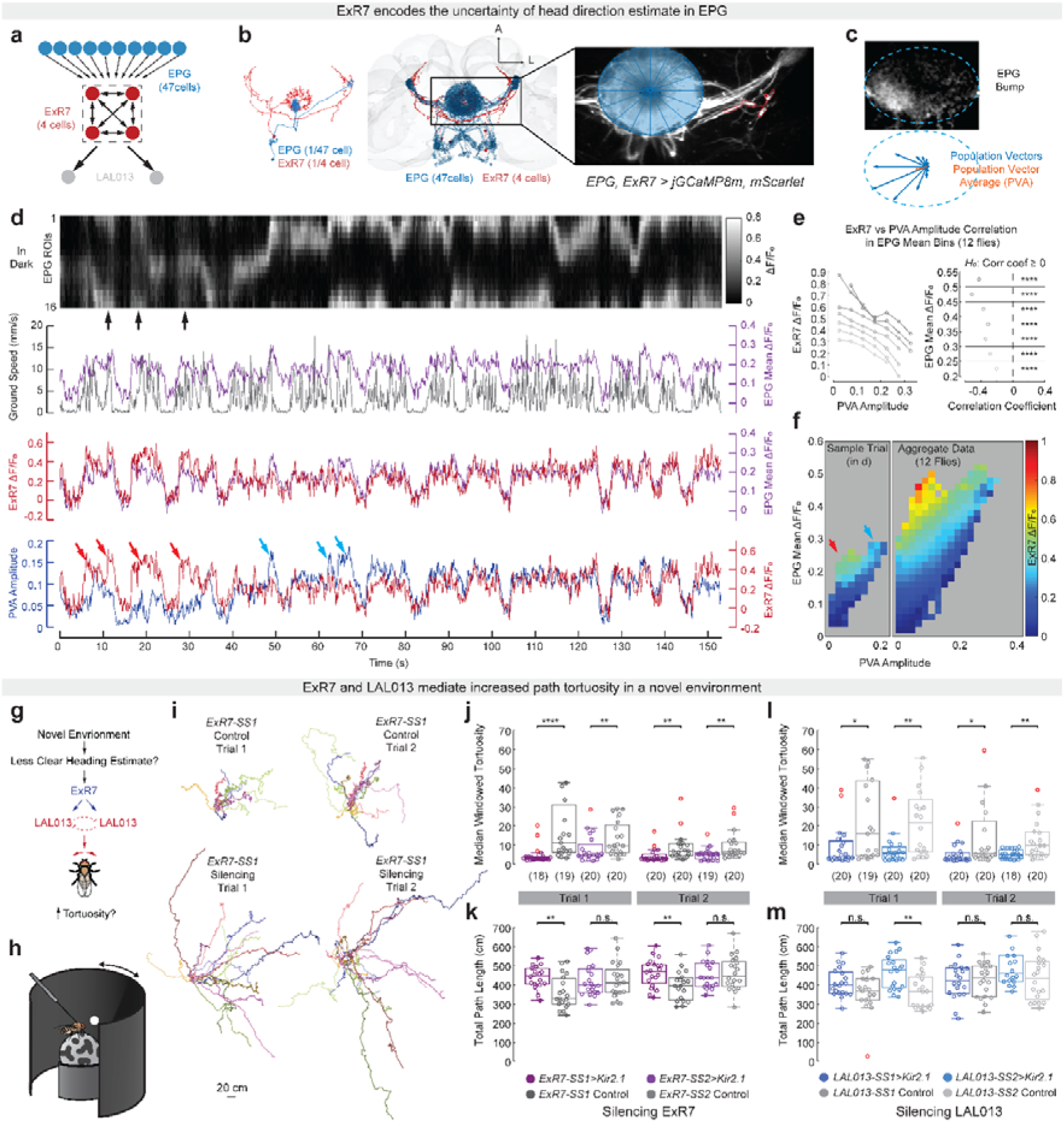
ExR7 is modulated by uncertainty of heading estimates to control path tortuosity via LAL013. **a**, Connectivity between EPG, ExR7 and LAL013. **b**, EM image (FlyWire) with single cell (left) or all cells (middle) of EPG and ExR7 overlaid (dorsal views) and a matching frame of *in vivo* calcium imaging of both cell types (right). **c**, Top: an image of EPG bump overlaid with blue dashed ellipse indicating the arborization of EPG in ellipsoid body. Bottom: population vectors (blue) and population vector average (PVA, red) calculated from the EPG activity **d**, Example traces of EPG/ExR7 dual imaging experiments. First row, 16 EPG ROIs, with activity encoded in grey values and black arrows indicating moments that lack a coherent bump. Second to fourth rows, ground speed/EPG mean, ExR7 activity/EPG mean, or PVA amplitude/ExR7 activity overlaid. Arrows indicate moments when the PVA amplitude was low and ExR7 activity was high (red) or vice versa (blue). Note that EPG mean was comparable between these time points. **e**, Correlation between ExR7 and PVA amplitude binned by corresponding EPG mean. Left, each curve represents data from a bin of EPG mean. For each bin, average ExR7 activity is plotted against the corresponding PVA amplitude band. Right, the correlation coefficient between ExR7 and PVA amplitude in each bin of EPG mean. **f**, Average ExR7 activity colour-coded in different bins of EPG mean (Y axis) and PVA amplitude (X axis), calculated for data from the sample trial (left, red and blue arrows correspond to those in d) or from all data across 12 flies (right). **g**, Schematic of hypothesized function of the ExR7-LAL013 pathway. **h**, Schematic of the virtual reality with a disk stimulus. **i**. Trajectories of *ExR7-SS1* control and *ExR7-SS1>Kir* flies in both naïve and experienced trials with the disk stimulus. Trajectories are colour-coded by fly identity within the respective experimental group. **j**, Boxplots depicting the median windowed tortuosity of the trajectory (window size 20s) calculated per fly and per trial, for the first and second disk stimulus trials. The box shows the interquartile range (IQR). The line inside the box marks the median. Outliers are determined by boundaries of 1.5 × IQR from the first and third quartiles. Whiskers extend to the most remote non-outliner data points. Numbers in brackets are n of flies. **k**, Same as (j), but for the total path length of the trajectory. **l**, Same as (j), but for two SS lines targeting LAL013. **m**, Same as (l), but for the total path length of the trajectory. ****, *p*<0.0001; **, *p*<0.01; *, *p*<0.05; n.s., *p*>0.05; Pearson’s *p*-values (e) for the null hypothesis of population correlation coefficient ρ ≥ 0; Wilcoxon tests (j-l) comparing the silenced and control genotypes.

We quantified EPG activity by calculating two metrics: the average activity across each of sixteen wedge-shaped ROIs tiling the ring (EPG mean) and a population vector average (PVA). The orientation of the PVA represents the head direction estimation and its amplitude represents the informativeness of that estimation^21^ (Fig. 4c). As reported previously, EPG neurons often but not always show a localized activity bump (Fig. 4d top row, black arrows indicate moments lacking a localized bump), and both EPG mean and the PVA amplitude are highly dynamic, with large moment-to-moment fluctuations^21,53,54^ (Fig. 4d).

Consistent with the all-to-all connection pattern between EPG and ExR7 (Fig. 4a), we found that ExR7 activity is tightly correlated with EPG mean (Fig. 4d and Fig. S5a-c), with both cell types more active during walking bouts than quiescence (Fig. S5c). ExR7 activity is noticeably less well correlated with the PVA amplitude, even though PVA amplitude and EPG mean are themselves, as expected, highly correlated (Fig. 4d and Fig. S5a-c). Moreover, when we compared time points at which the EPG mean stays relatively constant, ExR7 activity was in fact inversely correlated to PVA amplitude (Fig. 4d, red and blue arrows; Fig. 4e). Plotting EPG mean, PVA amplitude, and ExR7 activity for the sample trials (Fig. 4f and Fig. S5d) or across all flies and all sessions (Fig. 4f and Fig. S5e-f) showed clearly that ExR7 activity varies positively with EPG mean but negatively with PVA amplitude (at constant EPG mean). We conclude that ExR7 is recruited whenever EPG activity is high but dispersed (i.e. the EPG bump is weak), and thus uninformative as a sense of head direction. In such instances, by activating LAL013, ExR7 could promote turns without specifying their directionality.

### Silencing ExR7 or LAL013 reduces turning in a novel environment

We inferred from these imaging data that the ExR7/LAL013 pathway might elicit turns whenever the fly needs to increase its environmental sampling. This occurs, for example, when the fly encounters a new visual environment. In this situation, the fly’s head direction system does not accurately track the animal’s current orientation^54-56^. We predicted, therefore, that flies would show increased turning, and hence more tortuous walking paths, in a novel visual environment, and that this increase in tortuosity should be suppressed by silencing ExR7 or LAL013 (Fig. 4g).

We tested this hypothesis by measuring the walking trajectories of tethered flies walking in a visual virtual reality. Flies were presented with a visual panorama consisting of a single white disk on a black background (Fig. 4h). We presented the same panorama twice, such that flies were initially naïve and later experienced with this environment. In both naïve and experienced trials, control flies showed menotaxis behaviour, choosing arbitrary walking directions with respect to the azimuthal position of the white disk (Fig. 4i and Fig. S6a-c). While the flies’ walking trajectories were generally directed, they were tortuous with many small loops and small changes in direction (Fig. 4i and Fig. S6a-c). This tortuosity was reduced on the second trial, consistent with our view that it represents exploration in a novel visual environment (Fig. 4i-j).

When we silenced ExR7 neurons using either of the two SS lines, flies showed menotaxis trajectories that were significantly straighter with fewer loops and changes in direction (Fig. 4i-j and Fig. S6a). This effect was less pronounced on the second trial than the first, as expected given that tortuosity is generally lower when the visual environment is familiar. For one of the two SS lines, the reduced tortuosity on the first trial was accompanied by slight increases in total path length, and translational and absolute rotational velocities (Fig. 4k and Fig. S6d-e). The weaker and less consistent nature of these effects suggest that reduced turning, rather than increased forward walking, is the primary consequence of ExR7 silencing. Very similar results were observed when we silenced instead the LAL013 neurons, again using either of two SS lines (Fig. 4l-m and Fig. S6b,c,f,g). In summary, silencing either ExR7 or LAL013 consistently reduced the tortuosity of directed walking trajectories during menotaxis in a novel visual panorama, supporting the hypothesis that the ExR7/LAL013 pathway promotes exploratory turns.

### Asymmetric DNa03 activity anticipates turns

Our results show that the steering circuit receives directional and non-directional inputs from the navigation centre and functions in both goal-directed navigation and exploration. To gain further insights into the computation performed by this circuit, we conducted bilateral calcium imaging on each neuron type in flies walking spontaneously on a spherical treadmill. It has previously been reported that, in this setup, the bilateral difference in DNa02 activity correlates well with angular velocity^8,9^. We confirmed this result and observed a similar correlation also for each of LAL013, DNa03, DNa11 and DNae003 (Fig. 5a and Fig. S7).

**Fig. 5:**
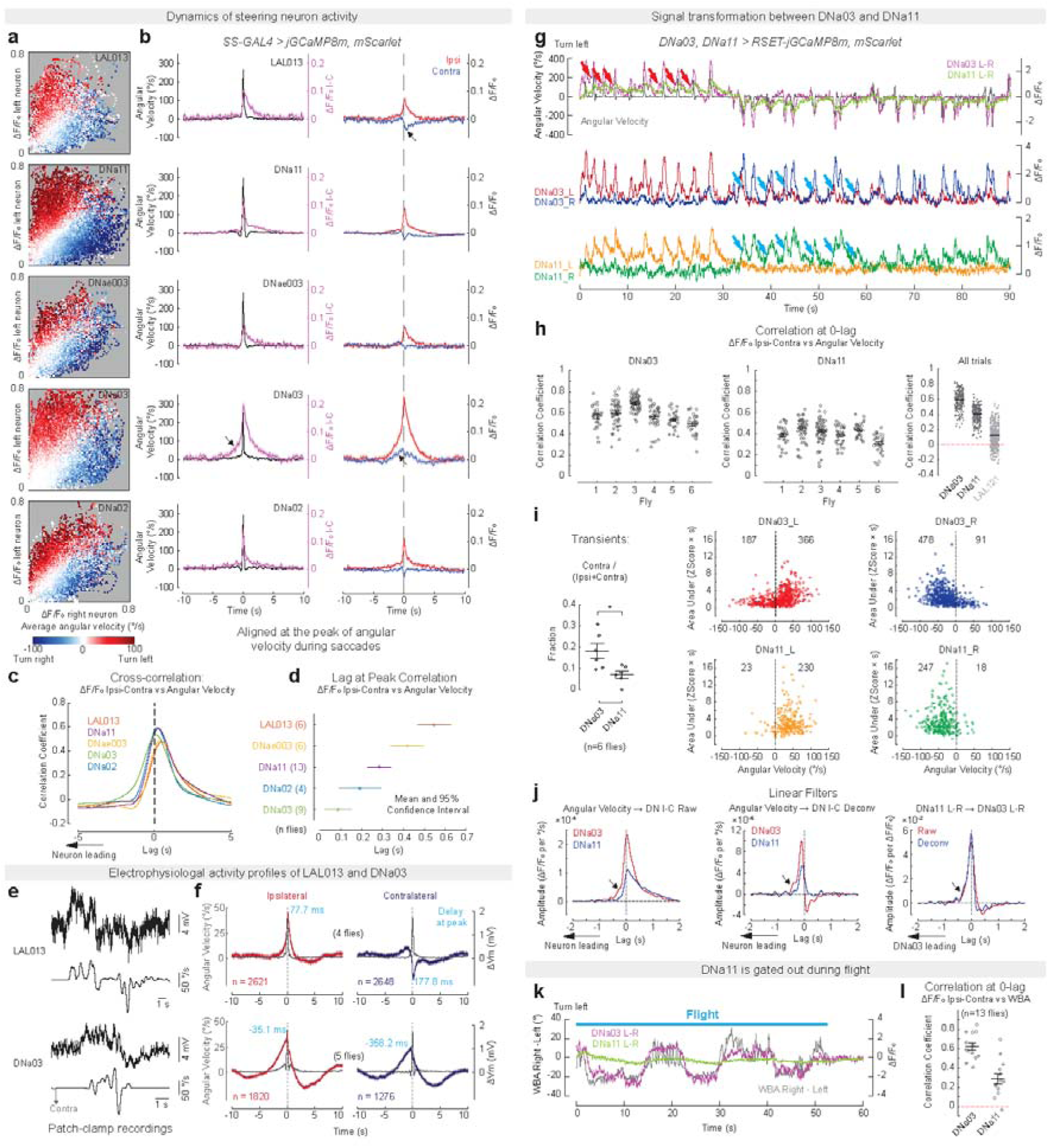
Signal transformation within the steering hierarchy. **a,** Heatmap showing fly’s average angular velocity binned by left (Y axis) and right (X axis) neuron activity for each cell type. **b**, Time series of angular velocity and activity of the ipsilateral and contralateral neurons for each cell type, aligned at the peaks of angular velocity during saccades (peak angular velocity ≥ 200 °/s). Left: angular velocity (black) and bilateral difference in neuronal activity (purple, lag at peak: LAL013/DNae003, 0.146 s; DNa11/DNa03/DNa11: 0 s). Right: activity in ipsilateral (red) and contralateral (blue) neurons (relative to the turning direction). Data are mean ±Ls.e.m. across turning events (n: LAL013, 1338; DNa11, 8819; DNae003, 1847; DNa03, 753; DNa02, 2442). Arrows: LAL013, reduced activity of the contralateral neuron during turns; DNa03, increased bilateral activity as well as the bilateral difference before turns. **c**, Cross-correlation between bilateral difference of activity and angular velocity. Data are averaged across flies for each genotype. **d**, Lags at peak and 95% confidence intervals for the cross-correlation data in (c). Numbers in brackets are n of flies. **e**, Example traces of patch clamp recordings of LAL013 and DNa03 neurons. **f**, Membrane potentials aligned with angular velocity peaks for turns with peak angular velocity greater than 20 °/s. Lags of neural activity peaks relative to angular velocity peaks are indicated in the graph. **g**, Example traces of DNa03 and DNa11 dual imaging. Bilateral differences of DNa03 and DNa11 activity overlaid with angular velocity during walking (top row). Red arrows indicate DNa03 ramping up activity prior to angular velocity changes. Activity of DNa03’s (middle row) or DNa11’s (bottom row) left and right neurons overlaid. Blue arrows indicate DNa03 showing bilateral activity prior to turns and DNa11 only showing ipsilateral activity. **h**, Quantification of correlation between bilateral difference of activity in DNa03 (left) or DNa11 (middle) and angular velocity in each 22-s trial from 6 flies. Pooled data across flies for comparison between descending neurons and LAL121 (right). **i**, Left, fraction of transients occurring in both neurons at contralateral turns. Right, transients of DNa03 and DNa11 activity extracted with the same criteria (see Methods). The area under each transient (integral of the amplitude) was plotted against angular velocity. Numbers indicate total transients ipsilateral or contralateral to the turning side. **j**, Linear filters from angular velocity to DNa03 or DNa11 bilateral difference in raw ΔF/F_0_ (left; half rise time/peak time/off time constant in s, DNa03: -0.11/0.02/0.34; DNa11: -0.08/0.04/0.73) or deconvolved ΔF/F_0_ (middle, DNa03: -0.21/-0.11/0.09; DNa11: -0.20/-0.09/0.14), or from DNa11 to DNa03 bilateral differences of activity (right, raw ΔF/F_0_: - 0.15/-0.01/0.15; deconvolved: -0.13/-0.01/0.11) during walking. The linear filters represent the impulse response of output variable to unit input variable at time 0. Arrows indicate ramping up of DNa03 prior to turns or DNa11 activity. **k**, Bilateral differences of DNa03 and DNa11 activity overlaid with wing beat amplitude (WBA) for an example trial during flight. **l**, Quantification of correlation between bilateral difference in DNa03 or DNa11 activity and WBA during flight. Lines and error bars indicate mean ±Ls.e.m. across trials for each fly (h) or across flies (i, l). *, *p* < 0.05, paired t-test between the two neurons (i).

To further explore the dynamics of these turning-related activities, we aligned the angular velocity and calcium signals around turning events, binning turns into saccades (peak angular velocity > 200°/s)^33^ and progressively slower turns at 50°/s intervals (Fig. 5b and Fig. S8). This analysis revealed that, in saccades, the bilateral difference of activity in all neuron types peaked largely in synchrony with angular velocity (Fig. 5b). For each of the descending neurons, this difference was primarily attributable to an increase in ipsilateral activity. LAL013 however formed a notable exception, in that the increase in activity of the ipsilateral neuron was accompanied by a decrease of comparable size in the activity of the contralateral neuron (Fig. 5b).

With respect to the timing of these signals, it was instead DNa03 that formed the exception. DNa03 was the only cell type tested in which the calcium signals began to rise a few seconds before the turn. Both the ipsilateral and contralateral DNa03 activities increased in anticipation of the turn, with the ipsilateral activity increasing more rapidly than the contralateral activity (Fig. 5b). Cross-correlation analysis between neural signals and angular velocity also reveals small but consistent time lags at the peak correlation among these neurons (Fig. 5c-d and Fig. S7b-e), whereby DNa03 peaks earliest and LAL013 last. DNa02, DNa11 and DNae003 showed intermediate lags, with the relative timing of their peaks roughly matching their connection strength to the two top-layer neurons (Fig. 2b, Fig. 5c-d and Fig. S7b,e).

These general patterns were also observed for slower turns (Fig. S8). For the slowest turns (<50°/s), however, only DNa03 and DNa02 showed significant activity changes. Across all angular velocities above 50°/s, the ipsilateral neuronal activity increased roughly linearly with angular velocity for all cell types except LAL013, in which it plateaued above ∼150°/s. DNa03 activity anticipated the turn in all cases, but it was only in saccades (>200°/s) that an increase could be detected in the contralateral as well as the ipsilateral neuron. These data are consistent with the view that DNa03 receives bilateral input from PFL2 only when the fly’s heading deviates widely from the goal and a large corrective turn is needed^23^.

We performed patch clamp recordings on LAL013 and DNa03 to corroborate these findings. These recordings confirmed the bilateral ramping of DNa03 activity prior to turns and the opposing changes in LAL013 activity in the two hemispheres during turns (Fig. 5e-f and Fig. S9). The higher temporal resolution of these recordings also showed that peak activity of DNa03 precedes the turn whereas peak LAL013 activity lags the turn (Fig. 5e-f).

### Turning signals are rectified and filtered between DNa03 and DNa11

The connectome suggests that DNa03 may act as a sensorimotor integration and distribution hub. Besides PFL2/PFL3, DNa03 also receives inputs from AOTU019/025, which mediates visual object tracking^16^, MBON neurons involved in associative learning^57^, and other high-order sensory neurons (Fig. S10). On the other hand, DNa03 provides direct synaptic input to 8 descending neuron types and indirect input to an additional 48 (Fig. S10, considering connections with a minimum of 50 synapses in FlyWire^42,43^). We hypothesize that DNa03 integrates many different turning drives to compute a unified angular velocity signal and coordinate a population code of steering motor command by broadcasting to many other descending neurons. Given the short latency between activity peaks of DNa03 and DNa11, as well as DNa11’s position near the top of the hierarchy, we focussed on DNa11 in an effort to understand how steering signals are translated into turns.

To probe the signal transformation between DNa03 and DNa11, we simultaneously imaged calcium signals in both neurons of tethered flies during both walking and flight. We achieved higher signal-to-noise ratios and temporal resolution with the RSET-restored jGCaMP8m^58^. Consistent with our individual neuron imaging experiments, the bilateral differences in both neuron types correlate with angular velocity tightly during spontaneous walking, with DNa03 activity occasionally ramping a few seconds before turns (Fig. 5g and Fig. S11a-b). The correlation of DNa03 and DNa11 activities to angular velocity was consistently high across flies and trials (Fig. 5h), in contrast to the low and variable correlation we had observed with LAL121 (Fig. 3h). This difference suggests that both DNa03 and DNa11 encode steering motor commands and not, like LAL121, turning drives that are only probabilistically transformed into actions.

DNa03 and DNa11 in the same hemisphere show tightly correlated activity (Fig. 5g and Fig. S11c-d), but they also have some characteristic differences (Fig. S11). DNa03 consistently leads DNa11, has faster off kinetics, and often shows synchronized transients on both sides, especially immediately before a turn (Fig. 5g and Fig. S11a,c-e). DNa11 activity resembles a low-pass filtered version of the DNa03 signal (Fig. 5g and Fig. S11c,d,h,i). Additionally, whereas DNa03 transients often occurred on both sides, DNa11 signals were predominantly confined to the side ipsilateral to the turn (Fig. 5g,i and Fig. S11f).

By computing the linear filters and cross-correlation between angular velocity and neural activity, we found that the initial ramping up, main rising and peaking time of DNa03 all consistently lead DNa11 (Fig. 5j and Fig. S12). DNa11 decays much slower than DNa03 and appear to integrate DNa03’s input over time. The slower off kinetics and integral property of DNa11 may reflect differences in dendritic integration, circuit mechanisms or cellular properties involving calcium buffering. We deconvolved the calcium traces to extract the putative spiking probability of both cell types to mitigate the contribution of calcium buffering and better uncover the timing of neural activity (Fig. S11g). The linear filters and cross-correlation based on the deconvolved neural signals suggest that the integral property of DNa11 is likely part of the neural computation it performs rather than an artefact from calcium handling (Fig. 5j and Fig. S12).

During flight, the bilateral difference in DNa03 activity correlates well with left-right difference in wing beat amplitude, which is proportional to yaw angular velocity^59^, but DNa11’s activity is largely suppressed (Fig. 5k-l and Fig. S13). These results suggest that the cross-modality angular velocity signal encoded by DNa03 is flexibly processed by downstream descending neurons to execute the turn in a context-appropriate manner.

We conclude from these dual imaging experiments that the signal transformation between DNa03 and DNa11 marks the transition from a preparatory and generic steering signal to the motor command to execute a large-amplitude walking turn.

### DNa11 controls all six legs to execute a walking saccade

There are many different patterns of leg movements that could result in a turn. For example, the descending neurons DNa02 and DNg13 implement turns by altering the ipsilateral and contralateral stride length, respectively^9^. The turns these neurons elicit are however relatively slow, with a peak angular velocity that is far below the ∼300°/s seen in a typical saccade^33^ or upon DNa11 activation (Fig. 2c). Furthermore, saccadic turns can occur without any forward movement and therefore cannot be solely due to changes in stride length.

To investigate how DNa11 alters leg kinematics to execute a saccade, we used the DeepFly3D system^60^ to extract the angles of all joints on all legs from high-speed videos of flies walking on a ball (Fig. 6a and Supplementary Video 3). The changes in leg kinematics upon unilateral DNa11 activation were highly stereotyped, allowing us to extract an “average step” for each of the six legs across 10 flies (Fig. 6b-c). This analysis revealed that unilateral DNa11 activation causes all six legs to sweep around the body. This stepping pattern likely arises from movements of the thorax-coxa and trochanter-femur joints^61^ (Fig. 6c).

**Fig. 6:**
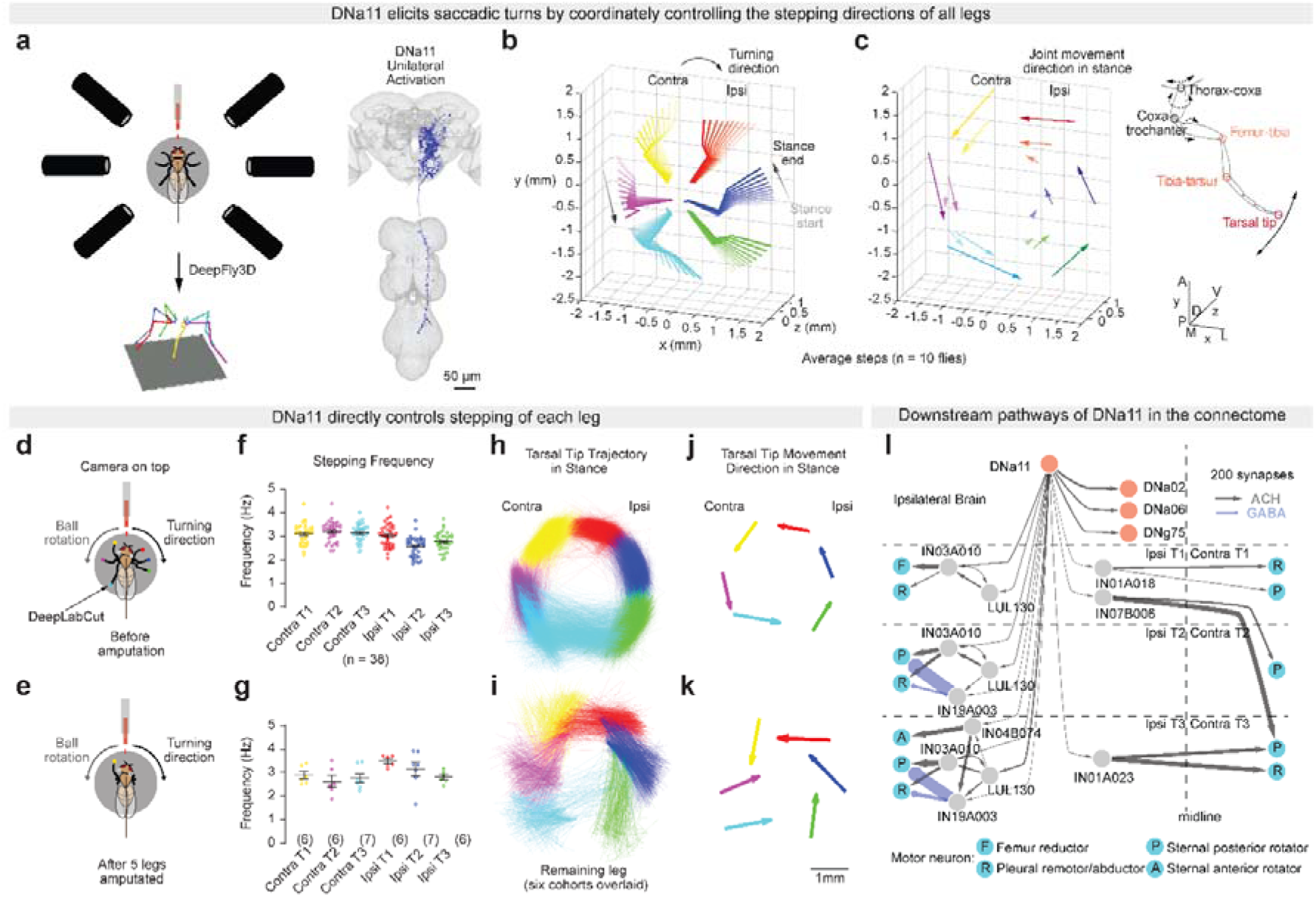
DNa11 coordinatedly controls all six legs to execute a walking saccade. **a**, Schematic of experimental setup for 3D leg kinematics upon unilateral DNa11 activation. **b**, Average steps of all six legs in stance phase during unilateral DNa11 activation. **c**, Left, stance movement vectors for femur-tibia and tibia-tarsus joints and tarsal tips (colour-coded for the ipsilateral front leg as in the schematic), for the average steps in (b). Right, schematic showing a fly leg. Solid arrows indicate the major movements during DNa11-triggered turns. Dashed arrows indicate candidate joint movements that could lead to the observed tarsal tip movements. **d**, **e**, Schematics of a fly walking on a ball before (d) and after (e) leg amputations. Tarsal tips and edges of the head are tracked from the top view using DeepLabCut. **f**, Stepping frequency of all six legs in intact flies during unilateral DNa11 activation. **g**, Stepping frequency of the remaining leg in six cohorts of amputated animals. **h, i,** Tarsal tip trajectories of all six legs in intact flies (h) or of the remaining leg in six cohorts of amputated animals (i). **j, k**, Vectorized tarsal tip movement direction of all six legs in intact flies (j) or of the remaining leg in six cohorts of amputated flies (k) during stance phase. **l**, DNa11’s top downstream descending pathways in the brain and pre-motor pathways in the VNC. Line widths indicate synapse numbers.

The sweeping movements of all six legs might arise through direct and coordinated control of all 6 legs, or by DNa11 acting primarily through a single leg or pair of legs and the others following passively due to mechanical coupling with the substrate. We performed a series of amputation experiments to distinguish between these possibilities. In these experiments, we amputated all but one leg below femur, so that only the intact leg has traction on the substrate (Fig. 6d-e, Supplementary Video 4 and Supplementary Video 5). Because the single intact fore-, mid- or hindleg could be ipsilateral or contralateral to the single DNa11 neuron stochastically targeted for activation, these experimental flies fall into six cohorts. In all six cohorts, the single intact leg stepped at the same frequency upon unilateral DNa11 activation as it did prior to amputation (Fig. 6f-g). The head also rotated to the ipsilateral side in all cases, as it does in natural saccades^34^ (Fig. S14a). We infer from these observations that the amputation does not interfere with either the motivation to turn or the ability to step. Not surprisingly, the ability to turn was noticeably affected (Fig. S14b-d), but all flies except those retaining the contralateral T2 leg were still able to turn to the ipsilateral side.

Importantly, in all 6 cohorts the footfall pattern of the single intact leg closely resembled the pattern observed for the same leg in flies with all their legs intact (Fig. 6h-k). The stepping patterns of the ipsilateral legs were generally more resilient than those of the contralateral legs (Fig. 6i,k). Nonetheless, the striking ability of each leg to step roughly in the correct direction even when all other legs are amputated strongly suggests that DNa11 coordinately controls the motor circuits of all six legs.

DNa11 has abundant synaptic outputs in both the brain and VNC. We found however that unilateral DNa11 activation is still capable of eliciting turning in decapitated flies, albeit much more slowly (Supplementary Video 6). This observation is consistent with the view that DNa11 controls turning by acting both directly on leg motor circuits and indirectly through its subordinate descending neurons. Among those subordinate descending neurons, DNa02 has been shown to control stride length^9^, and the connectome suggests that DNa06^62^ might be involved in controlling neck rotation^63^. Indeed, we saw the neck turns robustly towards the ipsilateral side with minimal body turning when we unilaterally activated DNa06 (Fig. S14e-h). Within the VNC^64-66^, DNa11 has few direct synaptic connections with motor neurons but does target many classes of premotor neuron. We mapped the strongest bi-synaptic pathways from DNa11 to leg motor neurons in all six legs (Fig. 6l). This analysis revealed that DNa11 primarily targets the body wall muscles that control coxa remotion and rotation and the femur reductor muscles that may mediate femur rotation^61^. The predicted functions of these motor neuron pathways align well with direction of leg movements during stance phase upon DNa11 activation. However, DNa11 also targets ipsilateral motor neurons that are predicted to move the legs backwards, as occurs during swing phase. These connections are via the IN03A010 neurons, which form reciprocal connections with the swing-initiating^67^ LUL130 neurons. The alternating activation of these motor pathways during swing and stance could account for the leg movements that turn the fly upon DNa11 activation.

## Discussion

The transformation of internal incentives and sensory streams into precise motor patterns remains a fundamental “black box” in both systems neuroscience and autonomous robotics. Here, we provide detailed circuit-level understanding of how this transformation is implemented within a central steering circuit that operates across behavioural contexts, supporting both goal-directed navigation and exploratory search, and controlling turning during both walking and flight. We show that this circuit is organized hierarchically, progressively transforming navigational signals from the central complex and higher-order sensory inputs into coordinated steering manoeuvres.

Our behavioural and connectomic analyses suggest that DNa03 and LAL013 play key roles in controlling turns during goal-directed and exploratory locomotion, respectively. The distinct dynamics of DNa03 and LAL013 activities, revealed by our imaging and electrophysiology experiments, suggest that DNa03 and LAL013 have complementary roles in instructing a turn. DNa03 appears to initiate the turn, while a winner-take-all mechanism involving LAL013 likely has a key role in reinforcing the turn and stabilizing its direction.

The DNa03 and LAL013 neurons are reciprocally connected and share many downstream neurons, including DNa11 and DNae014, which control walking and flight saccades, respectively. We propose that this circuit balances the need to perform a corrective turn when the fly deviates from a clear goal with the urge to perform more exploratory turns when its goal or heading are less certain. When the goal is clear and there is strong evidence that the fly is off course, the PFL2 and PFL3 inputs to DNa03 initiate a corrective turn, which may be reinforced by LAL013. When the goal or heading is less clear, or the surroundings unfamiliar, the PFL2 and PFL3 inputs to DNa03 might be outweighed by the ExR7 inputs to LAL013, directly triggering a large amplitude, self-reinforcing turn in a random direction.

### Effector-agnostic motor intent

A central challenge for the steering control system is the need to resolve two distinct computational problems. First, multiple incentives can justify a course correction and these signals may conflict. These variables must therefore be integrated and weighed to select an appropriate action. Second, even once a turning goal is specified, the motor system faces a classic degree-of-freedom problem^68^, whereby many distinct motor patterns can produce a turn of the same direction and amplitude. Thus, the system must also determine how to implement the selected action across different effectors.

A natural solution is a sensorimotor integration hub that is both convergent (integrating diverse inputs) and divergent (flexibly routing outputs to multiple motor channels). The connectomic position of DNa03 makes it a strong candidate for such a role. DNa03 receives multiple inputs that would incentivise a turn, such as deviation from an internal goal encoded by PFL2/3, drift of a visual target tracked by AOTU019/025, or scene matching based on familiar routes potentially signalled by MBONs. Moreover, our imaging results show that DNa03 activity precedes that of other descending neurons, ramps up prior to turning, and often does so bilaterally, suggesting involvement in decision-related dynamics. At the same time, DNa03 activity correlates with high fidelity to a low-dimensional kinematic variable—angular velocity—that is invariant across effectors (legs or wings). This stands in contrast to the weaker and more probabilistic correlations observed one synapse upstream in LAL121, indicating a transformation from distributed incentives into a more actionable motor intent.

Together, these observations suggest that DNa03 encodes an abstracted control variable—intermediate between decision variables and motor commands—that can be flexibly routed to distinct motor systems. This separation of “what to do” from “how to do it” parallels the concept of a hardware abstraction layer in engineered systems^69^ and recent approaches to cross-embodiment control in robotics^70^, where shared representations enable the same high-level policy to be implemented across different physical substrates. Such parallels highlight a potentially general solution, shared across biological and artificial systems, for achieving flexible and scalable control.

### Sensing uncertainty

Successful control systems must contend with uncertainty in their inputs and adapt their behaviour accordingly. It is well established that the nervous system can encode uncertainty, typically at the population level^71^, where downstream decoders are required to extract this information. Here, we provide evidence that a single neuron type can report the uncertainty of an internally generated navigational variable and use this signal in a closed loop to improve the fidelity of that representation. We find that ExR7 activity increases as the PVA amplitude of the EPG bump decreases, a proxy for the reliability of the heading estimate. Elevated ExR7 activity, in turn, drives stronger exploratory turning via LAL013, promoting increased environmental sampling and thereby facilitating recalibration of the heading representation.

The origin of the observed anticorrelation between ExR7 activity and EPG bump amplitude remains to be determined. One possible explanation is a non-linear dendritic integration mechanism. Because ExR7 tiles the entire ellipsoid body and receives input from the full population of EPG neurons, spatially localized EPG activity may saturate individual dendritic branches, leading to weaker overall activation compared to more distributed input patterns. This hypothesis is consistent with findings in pyramidal neuron dendrites, where inputs within a branch sum sublinearly at high strength, whereas inputs across branches sum more linearly^72^. Future experiments combining targeted optogenetic activation with dendritic imaging in ExR7 will be required to test this hypothesis directly.

Using estimates of information quality to guide adaptive control has strong parallels in engineered systems, such as the modulation of gain in a Kalman filter^73^, where increased uncertainty in the internal estimate leads to increased gain, promoting greater reliance on sensory input. Our results suggest that similar principles may be implemented at the level of identified neurons in biological circuits.

### Smooth execution

Control systems must balance fundamental trade-offs between sensitivity and robustness^74^. A common solution is to concentrate sensitivity in internal representations while enforcing robustness at the actuator interface, an idea consistent with optimal feedback control^75^ and dynamic systems^76^ frameworks. Our observations are consistent with such an organization: DNa11 activity exhibits low-pass filtering and rarely shows contralateral transients during turning, suggesting a temporally smoothed and robust control signal at the level of motor execution. Consistent with this, unilateral activation of DNa11 produces high-amplitude, saccadic turns, indicative of a high-threshold commitment as motor output.

The activity of DNa11 is likely shaped in part by inputs from LAL013. LAL013 neurons exhibit activity that is delayed relative to angular velocity and shows opposing changes across hemispheres during turns. In the absence of clear pathways for mutual inhibition between the two LAL013 neurons, this bilateral antagonism may arise upstream. One possibility is that LAL013 inherits a rotational velocity signal resembling an efference copy of ongoing movement. For example, GLNO neurons provide rotational velocity estimates to the central complex, exhibiting strong left–right antagonism and sigmoidal tuning to angular velocity^77^. LAL013 displays similar antagonistic structure and tuning, and its activity lags angular velocity, consistent with a derived signal. Although LAL013 does not receive direct input from GLNO neurons, it does receive input from candidate intermediates (e.g., PS196 and LAL139) that are proposed to relay rotational signals^77^.

Interestingly, bilateral activation of LAL013 produces prolonged turning in an arbitrary direction, consistent with winner-take-all dynamics and suggesting the presence of positive feedback linked to ongoing behaviour. Future work will be required to test whether such feedback contributes to the winner-take-all dynamics of LAL013 as well as the low-pass filtering and rectified directional tuning observed in DNa11.

More broadly, these results suggest that activity in the execution layer (DNa11) may reflect integration of an early and transient signal from DNa03 with a delayed and reinforcing component from LAL013. This division is reminiscent of a PI controller, in which proportional and integral components are combined to achieve both responsiveness and stability^78^. Such an architecture may provide a general solution for generating robust yet flexible motor outputs.

### Hierarchical motor control

We have described a stepwise transformation from navigational variables to abstract motor intent and ultimately to motor commands within a hierarchical circuit. At the lowest level, DNa11 still drives a rich and coordinated pattern of behaviour. Rather than simply modulating stride lengths of ongoing walking, DNa11 can simultaneously initiate stepping and coordinate stepping directions across all legs. Our connectomic analysis suggests that DNa11 recruits other descending neurons to control distinct motor subprograms while also directly targeting leg motor circuits through its own projections. Thus, even the execution layer implements a structured decomposition of motor output.

Classically, hierarchical motor control has been framed as a high-level division between motor planning, action selection, and execution, often associated with cortex, basal ganglia, and spinal circuits. Our results suggest that hierarchical organization is distributed across multiple levels of the sensorimotor pathway, with representations being progressively transformed from probabilistic and abstract variables to more concrete and discrete motor commands. This view is consistent with studies showing that hierarchical structure can be organized across scales in a self-similar manner, potentially reflecting a general design principle for biological control systems^79^.

Importantly, this hierarchy should not be viewed as purely feedforward. Feedback and sensory inputs can enter at multiple levels, and lower layers are capable of operating autonomously, as emphasized in subsumption architectures^2^. Higher layers can modulate these processes when necessary, but fast, context-specific pathways can bypass them entirely. For example, in steering, visually driven collision-avoidance signals may act directly on descending neurons without engaging the central complex^80,81^.

Similar organizational principles have emerged in robotics, where hierarchical and modular control architectures enable complex behaviour through layered policies operating at different levels of abstraction^82^. Together, these parallels suggest that distributing computation across multiple hierarchical layers, each performing partial transformations and local integration, may provide an efficient and robust solution for flexible motor control.

## Methods

### Fly genotypes

The table below shows the source of fly strains used in this study. See Supplementary Table 1 for the fly genotypes used to generate each data item.

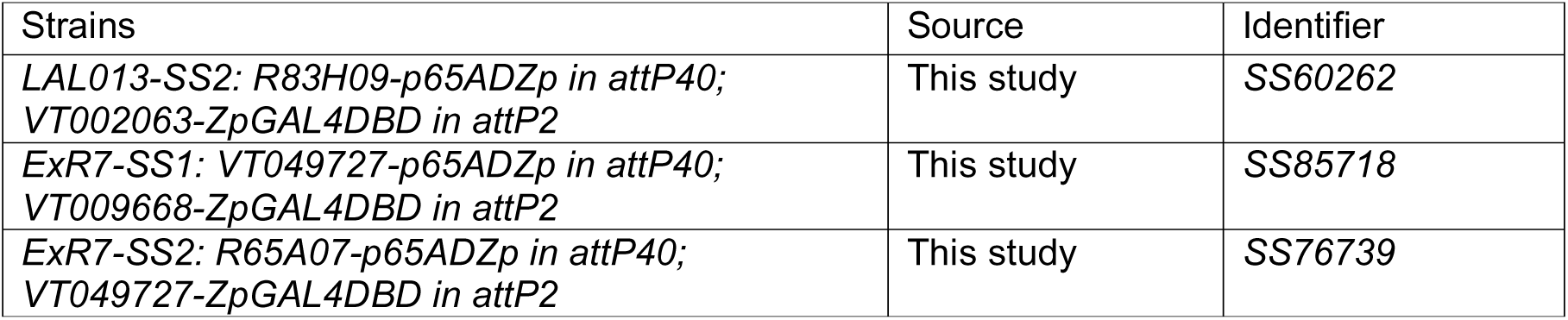
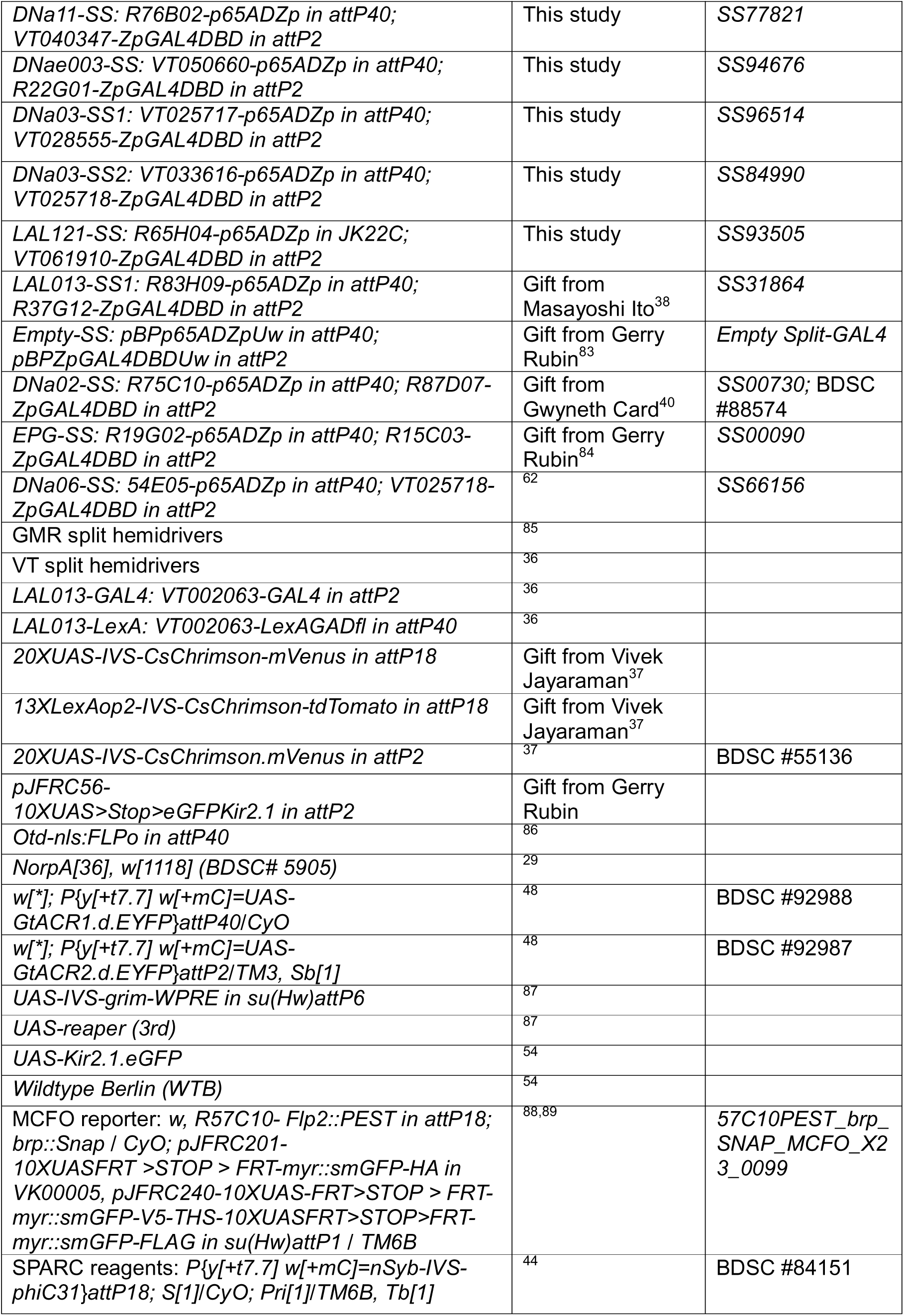
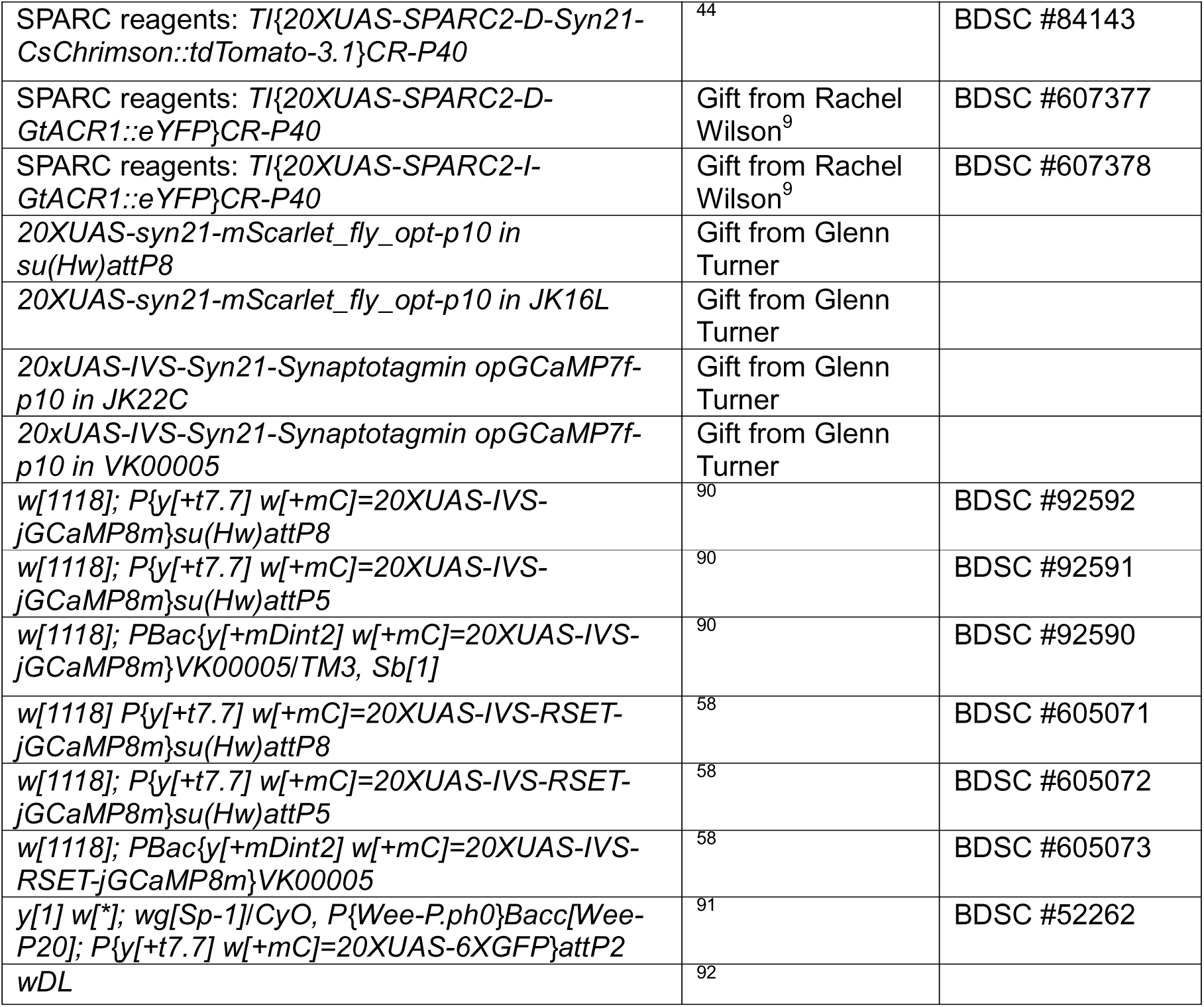

### Fly husbandry

*Drosophila melanogaster* flies were raised on standard semi-defined media at 25°C, 50-60% relative humidity and 12-12□h light-dark cycle. All flies for behavioural experiments, immunohistochemistry or functional imaging were sexed within 2 days after eclosion and aged in groups of up to 20 flies on fresh food. Flies used in optogenetic experiments were transferred to food supplemented with all-trans retinal (0.2□mM, Merck Sigma-Aldrich R2500) and kept under dark conditions (food tube wrapped with foil) for at least 48 hours before experiments. Flies were typically 3-8 days old when performing behavioural experiments or CNS dissection, except for SPARC stochastic activation and two-photon calcium imaging experiments, in which flies up to 20 days old were used to ensure strong reporter expression. In most of the experiments, only females were used. Exceptions for fly’s age and sex and special handling are described in the section of each experiment.

### Generation of split-GAL4 lines

We targeted specific neuron types (Fig. S1) following the strategy described in ref^38^. Briefly, we first selected VT^36^ and GMR^85^ enhancer tiles that have expression in the target neuron using the images^89^ and computational tools^93-95^ developed in Janelia Research Campus, HHMI. We then paired the corresponding split hemidrivers and screened for combinations that sparsely labelled the target neuron.

### Immunostaining and confocal imaging

Immunostainings were performed on the central nervous system (CNS) dissected from adult flies. For SPARC stochastic labelling experiments and verification of cell ablations (Fig. S4a-f), the dissected CNS were fixed in 4% paraformaldehyde for 25 minutes at room temperature (∼23°C). After three 15-minute washes in PBST (0.3% Triton X-100, Merck Sigma-Aldrich X100, and 0.05% Sodium Azide, Merck Sigma-Aldrich 71289, in 1X PBS), the samples were blocked in 10% goat serum (ThermoFisher 16210064) in PBST for 2 hours in room temperature or overnight at 4°C. Samples were then incubated in primary antibodies for 48 hours at 4°C. After 3 washes in PBST for 15 minutes each, samples were incubated in secondary antibodies for 48 hours at 4°C. After another 3 washes, the samples were mounted in Vectashield (Vector Laboratories, CA) or DABCO Mounting Medium (25 mg/ml of DABCO, Merck Sigma-Aldrich D2522, in 90% glycerol/10% 1xPBS, pH adjusted to 8.6) before being imaged at an Olympus SpinSR10 spinning disk confocal microscope with a 20× objective. Antibodies used were: primary rabbit Anti-RFP antibody (abcam ab62341, 1:1000) for tdTomato; primary rabbit anti-GFP (Merck/Millipore AB-3080P, 1:1000) for mVenus or eYFP; primary mouse anti-Bruchpilot (Developmental Studies Hybridoma Bank nc82, 1:20) for neuropil structures; secondary Alexa Fluor 488 goat anti-rabbit (Thermo Fisher Scientific A-11034, 1:1000); secondary Alexa Fluor 647 goat anti-mouse (Thermo Fisher Scientific A-21236, 1:1000).

All other immunostainings and confocal imaging (full pattern and MCFO of SS lines and *VT002063*) were done as described in refs^38,89^ and the protocols in https://www.janelia.org/project-team/flylight/protocols.

### Behavioural experiments

#### Open arena assays

Experiments in Fig. 1c-e, Fig. 1u-v, Fig. 2h-i and Fig. S2 were conducted in the Fly Bowl setup^96^. Briefly, video recordings were performed under backlit infrared LEDs at 30 fps with a resolution of 1024□×□1024 pixels. The bowl-shaped arena was covered with Sigmacote (Sigma) coated glass plate to prevent walking on the ceiling. A group of ∼25 flies was loaded into the arena and allowed to walk for 60□s before the first optogenetic simulation. A red-light source at 627□nm wavelength illuminated the entire arena uniformly to activate CsChrimson. Each red-light stimulus was 5□s in duration and at 60-s intervals (onset to onset). For Fig. 1c-e and Fig. S2, male flies were given nine episodes of red-light stimuli in one experiment with three consecutive episodes for each of three intensity levels from low to high: 0.158, 0.288, and 0.456□mW/mm^2^. Experiments in Fig. 1u-v were conducted in a different replicate of Fly Bowl with female flies given nine identical red light optogenetic stimuli at 0.145 mW/mm^2^, otherwise the same as above.

Experiments in Fig. 2h-i were performed in a Fly Bowl illuminated with 525nm green light. Single female flies were assayed through two sequential sessions before dissected. The first session comprises three of 60-s green-light stimuli at 0.033 mW/mm^2^ interspersed with 60-s light-off periods. The second session comprises 24 of 5-s green-light stimuli at 0.125 mW/mm^2^ interspersed with 5-s light-off period.

Experiments in Fig. 1f-g were conducted in a 90-mm diameter circular arena^34^ with single flies. To prevent thigmotactic behaviour the walls were heated up 37.5 °C by an insulated nichrome wire (Pelican Wire P2128N60TFEWT). The arena was covered with a glass plate pre-treated with Sigmacote. For optogenetic activation experiments, flies were stimulated with repeated red-light pulses (617 nm, 0.012 mW/mm^2^) of 50 ms, 100 ms, or 200 ms in duration, each followed by a no-stimulus interval such that the full cycle lasted 2 s; each stimulus duration was presented 10 times per block, and the full stimulation sequence was repeated across the recording (30 min). Because qualitatively similar effects were observed for all three stimulus durations, we only plotted responses to the 100-ms stimuli. For silencing experiments, individual flies were assayed for 18 minutes in darkness.

#### Miniature wind-tunnel assays

Experiments for odour-offset search were conducted in a miniature wind-tunnel as described previously^29^. Briefly, single flies were placed in rectangular arenas (14 by 4 by 0.17 cm), where they were exposed to a constant flow of filtered, humidified air, defining the wind direction. Into this airflow we injected pulses of odour with rapid onset and offset kinetics. A 525-nm blue LED light (NFLS-B300X3, SuperBrightLEDs) was used for optogenetic stimulation. Flies were imaged from below the arena using a monochrome USB 3.0 camera (Basler: acA1920-155um) and a 12 mm 2/3’’ lens (Computar: M1214-MP2). Imaging and stimulus delivery were controlled by custom software written in Labview (National Instruments, Austin, TX). Tracking code was written in Labview to extract position and heading at 50Hz. We used SS lines that were backcrossed for 7 generations to a *w^1118^, norpA^36^* strain, as flies with this genetic background showed robust activity in the assay^29^. Flies were starved 18-24 hr prior to the experiment, and were tested for approximately 3 hr (from ZT 0–4), in a series of 70-s trials under three conditions (15-s light; 70-s light and light off; intensity: 45 μW /mm^2^) with blank (wind only) and odour (10-s 1% ACV) trials randomly interleaved.

#### Spherical treadmill walking assays

The spherical treadmill was based on a polypropylene ball (PP ball, 6 mm in diameter, Spherotech GmbH, Germany) supported by a custom-built ball holder (Electronics workshop, Zoological Institute, University of Cologne, Germany) as described previously^97^. Compressed air was carefully adjusted to the minimal level that can spin the ball. Six Basler acA1920-155um USB3 cameras equipped with Infinistix 1□×□94□mm lens (with a 5.8□mm aperture retainer ring and an infrared filter; Infinity Photo-optical Company, CO) were placed on the side of the ball at the same height as indicated in Fig. 6a. Another Basler acA1920-150um USB3 camera equipped with a Computar MLH_10X lens (CBC America, NC, with an infrared filter) was placed on top of the ball. A custom-built infrared-LED ring (wavelength: 880□nm, Electronics workshop, Zoological Institute, University of Cologne, Germany) was positioned around the top camera lens for illumination. Custom C++ software was used to record synchronized videos from all the cameras. Videos with a frame size of 960 × 480 pixels were recorded at 200 fps. A custom built 660-nm diode laser with fibre coupling (Electronics workshop, Zoological Institute, University of Cologne, Germany) was used to deliver the red-light stimulus for CsChrimson. A fibre coupled 470□nm LED (M470F1, Thorlabs, NJ) controlled by a custom LED driver (QBI Workshop, University of Queensland, Australia) was used to deliver the blue-light stimulus for GtACR2. Multimode optical fibres (Ø200 µm, 0.39 NA for red light; Ø200□µm, 0.50 NA for blue light; Thorlabs, NJ) were used to deliver light to the flies (both with SMA coupling to source and clean-cut bare end pointing to the fly). The red-light intensity was ∼0.73□mW/mm2 and the blue-light intensity was ∼0.11□mW/mm2 as measured at the fly’s distance. An Arduino UNO (Arduino.cc) was used to trigger and synchronize the cameras and optogenetic stimuli.

Before an experiment, a fly was anaesthetized using a Peltier cold block (QBI Workshop, University of Queensland, Australia) set to 0°C and placed in a fly-sized rectangular groove. A drop of adhesive “Not-A-Glue” (Bondic, Canada) was applied to the notum of the fly using a thin metal wire. A bent metal pin fixed on one end of a rod was lowered by a manual manipulator to gently touch the adhesive. Then the adhesive was cured by LED light (provided with the adhesive). The tethered fly was then transferred to the ball setup and positioned to walk comfortably on the ball under video guidance. The experiments were performed at room temperature (∼23°C) in very dim ambient light.

For the SPARC stochastic activation experiments, a fly underwent 10 sessions of recording before dissection to check the expression pattern. Each session lasted 7 s (5-s constant red-light stimulus in the middle with 1-s light off before and after) and neighbouring sessions were spaced by 1 to 2 minutes. All flies were randomly selected and the expression patterns were determined post hoc.

For the leg amputation experiments, pre-selected flies of *DNa11>SPARC-D-CsChrimson* that reliably turned to one side were used. As DNa11 unilateral activation led to robust turning phenotypes that were visible under light from a dissection microscope, pre-selection of turning flies by visual observation was efficient and led to accurate prediction of the side of DNa11 labelling, which was subsequently verified by immunostainings. These flies were divided to six cohorts as described in the main text and the number of left and right turning flies roughly matched for each cohort. Each fly first underwent 10 sessions of recording while their legs were intact. The flies were then cold anesthetized and 5 of the 6 legs were amputated at the femur. Each of the amputated flies underwent another 10 sessions of recording with identical protocols before being dissected.

For the neuronal epistasis experiments, both males and females were used in a 1:1 ratio for each genotype. Each fly underwent 10 sessions of recording. Each session lasted 27 s and the neighbouring sessions were spaced by at least 2 minutes. For each session, a 25-s constant red-light stimulus was applied from 1 s after the recording start until 1 s before the recording end. Two 5-s constant blue light stimuli were applied 5 s and 15 s after the onset of red light, as indicated in Fig. 2e.

#### Tethered flight assays

The tethered flight assays were performed on the same treadmill setup, but with the fly raised so that it cannot reach the ball with its legs. The ball provided a uniform background for camera recording of the wing movement. Only the top camera was used, focused on the fly, with the ball out of focus. An infrared-LED ring light was set to be constantly on during the recording. The camera shutter time was set to the maximum allowed by the recording frame rate. Videos of a frame size 960 × 480 pixels were recorded at 400 fps. Under such conditions, the motion blur of fly wings during flight could be captured to extract the wingbeat amplitude. A gentle blow was applied to motivate the flight if the fly stopped flying. The experiments were performed at room temperature (∼23°C) in very dim ambient light.

A later subset of the *LAL013/DNa03>SPARC-D-CsChrimson* flies that underwent the treadmill walking assay were used for the flight assays. Those flies performed 10 flight sessions spaced by 1 to 2 minutes after 10 walking sessions and before being dissected. Red optogenetic stimulation and session duration were identical in the flight assays as in the walking assays.

#### Virtual reality assays (goal-directed navigation)

The same ball holder and 6-mm PP ball as described above were used for the virtual reality menotaxis assays. Custom hardware (Electronics workshop, Zoological Institute, University of Cologne, Germany) based on two motion sensors and 785nm infrared laser illumination was used to track the ball rotation as described previously^97^. The ball rotation was measured and recorded at 50Hz and sent to a computer via serial ports through USB connections. Two VuePix SF4 bendable LED panels, along with a N300 Nova MSD 300 sender box (Universal Lighting & Audio Pty Ltd, Australia) were used to display the visual stimuli. Each of these rectangular LED panels had 128 × 64 pixels. They were bent to form a cylinder surrounding the fly covering 240° and a gap of 120° was left open behind the fly. A bright blue vertical bar of 10 × 64 pixels (∼9.4° wide and spanning the full height of the panels) was displayed on a dark background. In closed loop, the bar position was controlled by fly’s rotational velocity with a gain of 0.7 (ref^23^). A third virtual panel on the back of the fly covers the 120° gap when computing the bar’s position (virtually 384 × 64 pixels surrounding the fly). The experiments were performed at room temperature (∼24°C) in a dark enclosure in which the LED panels were the only light source. The tethered flies were lightly starved (1-3 hours in a chamber with wet tissues) before experiments. After being carefully positioned on the ball, they were given half an hour to acclimate. If the fly showed reasonable walking activity, it entered the perturbation sessions, otherwise it was given another half an hour in closed loop before entering the perturbation sessions. A perturbation session lasted 31 minutes and a bar jump alternating to the left or right by 90 pixels (∼84.4°, designated as 90° trials) or 180 pixels (∼168.8°, designated as 180° trials) were performed every minute. After a bar jump the fly was given control of the bar position immediately. A fly typically started with a 90° perturbation session (90° trials × 30) and then entered a 180° perturbation session (180° trials × 30) and continued to alternate between 90° and 180° sessions until the walking activity significantly dropped (typically after 3-6 sessions). A custom MATLAB (Mathworks, MA) program was used to read the ball tracking signals, update the display on the LED panels (using functions of Psychophysics Toolbox Version 3, ref^98^) and record the data.

#### Virtual reality assays (path tortuosity modulation in novel environment)

A virtual reality (VR) setup was used as previously described^54^. Briefly, flies were positioned on a spherical treadmill with integrated in-line heating, in front of a panoramic screen. We used the VR system to simulate three visual environments consisted of a different cylindrical, grey-scale panorama spanning 360° in azimuth^54^: Disk stimulus (“Disk”): white 6°-wide circular disk positioned just above the horizon line on a dark background. Full gradient stimulus: gradient between white and black, following a sinusoidal profile along the azimuth. Half gradient stimulus: same profile as the full gradient, but with half the modulation depth, spanning between light grey and dark grey. The visual cues were simulated as having infinite distance to the animal, allowed flies to change their azimuthal orientation relative to the visual panorama but not their distance.

For the spherical treadmill we used of a polyurethane foam ball (8.8mm diameter, FR-7110, Last-A-Foam, General Plastics Manufacturing Company, Tacoma, WA, USA) floating freely on an aircushion on top of a ball holder machined from aluminium. The aircushion was maintained through a constant airflow controlled through a flow meter (Alicat Scientific, Tucson, AZ, USA) at 0.74 L/min. To control the temperature of the fly during experiments, the airstream was heated through a coil wrapped around the ball holder. The temperature was controlled in closed loop using a thermocouple attached to the ball holder and connected to a controller, which also reported the current temperature such that the experimenter could wait with the experiment until the desired temperature was reached. The ball was illuminated with infrared LEDs and its motion was captured by a high-speed camera (Basler daA1920-30um) at 130 Hz. FicTrac^99^ was used to track the ball motion and the tracking data was sent via a socket port to the VR software, where the 130 Hz Fictrac updates were integrated into VR updates at 120 Hz.

A triangular screen used for visual display was made from two faces (5.8-cm wide and 9.5-cm high each), arranged in a 90° angle to the front of the fly. The fly was positioned 2 cm in front of the point where the two screens meet, such that the edges of the screen extended to 117° to the left and right, respectively, resulting in a 234° panoramic display. The fly was positioned at 3.16 cm relative to the bottom edge of the screen, such that the ball obscured the bottom edge, and the top edge of the screen extended 53.93°-77.44° from the horizon depending on the azimuthal direction. The screens were made from a diffuser sheet (V-HHDE-PM06-S01-D01, BrightView Technologies) and images were back-projected onto the screen using two DLP projectors (DLPDLCR2010EVM, DLP® LightCrafter™ Display 2010 Evaluation Module, Texas Instruments). To correct for the in-built gamma correction of the projectors, we measured the output brightness with a power meter and calculated a correction function for each projector. This function was applied to texture images before loading them into our VR environment to ensure accurate brightness and contrast.

All experiments were performed at 30°C and 35-50% relative humidity. At least 3 days before the experiment, 4 to 6-day-old mated female flies were split into groups and phase-shifted by different amounts to ensure that flies could be measured during their evening activity period throughout the day. To increase motivation for walking, flies were also wet starved for 24 h prior to the experiment. Thirty minutes before the experiment, flies were cold-anaesthetized on ice and glued to a thin tungsten tether at their thorax using UV-curable glue (BONDIC, Kranzberg, Germany). After tethering, flies were allowed to recover for 5 minutes before being placed on the ball. To ensure active walking behaviour during the experiment, the flies were allowed to acclimate to the ball for 15 minutes before the start of the experiment. Each experiment began with a priming trial with the disk stimulus, followed by three further visual conditions (disk, full and half gradient) presented in randomized order. Each trial, including the priming trial, lasted 8 minutes. During the priming trial, flies got accustomed to the VR setup and likely learned to associate the provided visual stimuli with their head direction estimate^54^. If a fly showed no walking activity during the priming trial, the experiment was aborted, and a new fly was prepared. On average, this led to the exclusion of 1 animal per experimental group.

### Two-photon calcium imaging

Two-photon calcium imaging was performed on a Thorlabs Bergamo II microscope equipped with a Galvo-Resonant scanner (Thorlabs, NJ). A piezo objective focus module with 400□µm travel distance (Physik Instrumente GmbH & Co. KG, Germany) was used to control an Olympus XLUMPLFLN 20× (2.0 mm working distance, 1.0 NA) or a Nikon N40X-NIR 40× (3.5 mm working distance, 0.8 NA) water immersion objective with. A Mai Tai DeepSee Ti:Sapphire laser (Spectra-Physics, CA) was tuned to 920nm or 960□nm for two-photon stimulation of GCaMP and mScarlet at the same time. We used 960 nm for the initial single-cell-type imaging dataset but 920nm for all the dual-cell-type imaging datasets to prioritize the signal for GCaMP. A Pockels Cell (Conoptics Inc., CT) was used to control the laser intensity. A 635□nm long pass dichroic was used to pass the stimulation laser and reflect the fluorescence from the sample to a pair of GaAsP PMTs (Hamamatsu, Japan). A 562□nm long pass dichroic was placed between the two PMTs. A 525/50□nm and a 607/70□nm band pass filter were placed in front of the PMTs that were used to detect green and red fluorescence respectively. During imaging, maximally 20 mW laser power was allowed at the sample. ThorImage software (Thorlabs, NJ) was used to control the microscope and record the images.

The same spherical treadmill, tracking system and visual display for the virtual reality experiments (goal-directed navigation) were used for two-photon imaging. The same vertical blue bar as described above was used in closed loop or open loop with fly’s rotation in a subset of EPG/ExR7 dual imaging experiments. In those cases, the LED screen for visual display was covered with four layers of blue gel filter (R381 Baldassari Blue Rosco Filter) to isolate the light from PMTs. In all other experiments, imaging was performed in the dark. A GS3-U3-23S6M-C USB3 camera (FLIR, Canada) equipped with a Computar MLM3X-MP75□lens (CBC America, NC, with an infrared filter) was used to monitor the fly’s behaviour during imaging. A custom-built infrared-LED ring (wavelength: 850□nm, Electronics workshop, Zoological Institute, University of Cologne, Germany) was positioned around the camera lens for illumination. Spinnaker software (FLIR, Canada) was used to control the camera and record videos at 200 fps with a resolution of 640 × 480 pixels. An Arduino UNO (Arduino.cc) was used to trigger and synchronize two-photon imaging, ball tracking and the camera to monitor fly behaviour. The timing of each frame start for two-photon imaging, ball tracking and camera video were recorded with the ThorSync software (Thorlabs, NJ).

Experiments were performed at room temperature on predominantly female flies. Before an experiment, a fly was anaesthetized using a Peltier cold block (QBI Workshop, University of Queensland, Australia) set to 0°C. We used fly holders with a 3D printed plastic frame and a metal plate glued together^100^. The metal plate was slightly modified from the original design to accommodate the 2mm working distance of the Olympus lens for imaging flies walking on the treadmill. The fly’s head and anterior notum were glued to the fly holder using the adhesive “Not-A-Glue”. Ice-cold extracellular solution comprising (in millimoles): 103 NaCl, 3 KCl, 5 N-Tris(hydroxymethyl)methyl-2-aminoethanesulfonic acid (TES), 8 trehalose, 10 glucose, 26 NaHCO3, 1 NaH2PO4, 2 CaCl2, and 4 MgCl2 (pH near 7.3 when bubbled with 95% (vol/vol) O2 and 5% (vol/vol) CO2 (carbogen)) was applied to the chamber of the fly holder to immerse the top of the head from the opening in the metal plate. Then a hole was opened on the cuticle of the dorsal anterior side of the head using a sharp needle. A pair of fine forceps (Fine Science Tools, Germany) were used to dissect out the tissues in the hole to expose the brain. After surgery, the fly was given about half an hour to recover before being placed on the treadmill underneath microscope lens. The position of the fly was carefully adjusted using the camera preview so that it can comfortably walk on the ball. The fly was allowed to acclimate for 15 minutes to half an hour while we set up the recording environment. A typical session lasted about two and half minutes. As relatively strong laser power was used (∼10-20mW) to ensure sufficient signal strength, the fly was usually given several minutes to rest between sessions. Multiple sessions were performed on each fly until the fly’s condition deteriorated. Volumetric imaging of dual colours was performed on a rectangular frame (512-1024 × 256 pixels) exposing the neurites of a neuron type in both brain hemisphere. We typically acquired 8 optical slices spaced by 6 µm and this resulted in an imaging speed of ∼7 volumes per second. With RSET-jGCaMP8m, we could achieve a higher signal-to-noise ratio and therefore used higher frame rate of 55Hz, but staying at a fixed Z plane where we can best separate DNa03 and DNa11 neurites.

To image the neural activity during flight, the spherical treadmill was removed and a mirror was placed underneath the fly holder. The same FLIR camera and lens were used to take bottom view of the fly at 100 fps through the mirror to monitor the wing beat. The Nikon objective with 3.5 mm working distance and the fly holder of the original design^100^ was used to allow more space to accommodate wing beat. Other procedures were identical to imaging walking flies but the front legs were removed and minimal or no glue was applied on the notum to facility flight. A brief air puff was used to initiate flight before an imaging session.

### Electrophysiology

Dissection and mounting of flies on an air-suspended ball was performed as previously described^101,102^ on male flies 1-2-day post-elcosion. Experiments were performed at room temperature, and the brain was perfused in carbogenated external saline (103 mM NaCl, 3 mM KCl, 5 mM TES, 8 mM trehalose, 10 mM glucose, 26 mM NaHCO3, 1 mM NaH2PO4, 4 mM MgCl2 and 1.5 mM CaCl2, adjusted to ∼275 mOsm). Patch pipettes (6-10 MΩ) were made from borosilicate glass (1.5 mm OD, 1.12 mm ID, World Precision Instruments, TW150F-3) using a P-97 Micropipette Puller (Sutter) and filled with internal solution (120 mM aspartic acid, 10 mM HEPES, 1 mM EGTA, 1 mM KCl, 4 mM MgATP, 0.5 mM Na3GTP, 13 mM biocytin hydrazide (Biotium, #90060), pH 7.3, adjusted to ∼265 mOsm).

To visualize brain tissue and track ball movements, an infrared LED array (780 nm, Roithner Laser Technik, LED780-66-60) was placed behind the fly, at a 45° angle relative to the optical axis of the two tracking cameras. Ball movements were tracked at 4 kHz. GFP fluorescence was excited using a M617F2 LED (Thorlabs), connected to an optic fibre (M28L02, Thorlabs). Whole-cell patch clamp recordings in vivo were performed under a custom-built upright microscope^103^ using a 40x water-immersion objective lens (Zeiss). Current clamp data was filtered at 3 kHz before digitization, sampled at 10 kHz, recorded with a MultiClamp700B amplifier (Molecular Devices), digitized using a BNC-2110 and PCIe 6321 (both National Instruments), and acquired using the custom-written FlyVRena software^104^. The liquid junction potential (13 mV) was post hoc subtracted from all data.

We recorded from 12 LAL013 neurons, with 4 of them passing our stringent quality criteria regarding the behaviour of the fly (>3-s bouts of continuous locomotion) and the stability of the recording (average membrane potential of quality-controlled recordings: -61.2 +/- 6.5 mV), and 19 DNa03 neurons, with 5 meeting our quality criteria (average membrane potential: -50.3 +/- 7.5 mV).

### Data analysis

#### Neuron segmentation and rendering

Using the software VVD Viewer^105^, we rendered confocal image stacks registered to the JRC2018_Unisex brain and VNC templates^106^ in 3D and manually masked other neurons co-labelled in the image and segmented out the neuron of interest. The segmented images and electron microscopy images downloaded as .swc flies were overlaid together with the templates in VVD Viewer to render the images shown in Fig. 1b and Fig. 2a.

#### Connectomic analysis

The hemibrain v1.2.1^41^ and FlyWire v783 datasets^42,43^ were used to analyse the brain connectome for Fig. 1t, Fig. 2b, Fig. 3a, Fig. 4a-b, Fig. 6l, and Fig. S10. The MANC v1.2.1 dataset^64-66^ was used to analyse the VNC connectome for Fig. 6l. For the brain, three independent sets of weights were available for the connection between each pair of cell types (right brain in hemibrain and both hemispheres in FlyWire). They generally showed close agreement with each other, so we only presented the weights of hemibrain in Fig. 1t, Fig. 2b and Fig. 6l and weights of Flywire in Fig. 3a and Fig S10. The schematic in Fig. 4a does not show the exact connectivity, but a summary based on patterns seen in the hemibrain. The connection weights for FlyWire were synapse counts grouped by cell type and side and averaged across both brain hemispheres. But in case one hemisphere was clearly less well reconstructed, only the side with better reconstruction was used. For the VNC, the central connections in both hemispheres were in close agreement but the motor nerves were degraded to various extents for different legs^65^, so we used the mean weights for the central connections and the maximal weights for connections to motor neurons across the two hemispheres. The weights presented were summed synapse numbers across the neurons of a cell type. The cell type names considered existing literatures^10,40,107^, both brain datasets and MANC. We followed the nomenclatures that were temporarily used in FlyWire and adopted in literatures for DNae014 (ref^8^, current FlyWire type DNa15) and DNae003 (earlier version of this study, current FlyWire type DNae001). The transmitter prediction^108^ was based on FlyWire and MANC. The neuron types, weights and transmitter types were imported into Cytoscape^109^ for plotting the wiring diagrams. To analyse the downstream premotor pathways of DNa11 in the VNC, we first extracted all the bi-synaptic pathways from DNa11 to motor neurons and then ranked them by the indirect relative weights from DNa11 to each type of motor neurons in each leg. The indirect relative weights were calculated as the product of relative weights for DNa11-to-premotor-neuron connections and premotor-to-motor-neuron connections, in which the relative weight between two neurons was defined as the geometric mean of the pre-synaptically normalized weights and post-synaptically normalized weights between the two neurons. The top bi-synaptic pathways for each leg and the premotor neurons mediating them were selected for further analysis and plotting the diagrams in Fig. 6l. To generate Fig. S10, those with direct inputs of at least 150 synapses were included for DNa03 upstream neurons and only descending neurons within two hops and synapse counts no smaller than 50 (at least in one hemisphere) for each hop were included for DNa03 downstream neurons. See Supplementary Table 2 for cross-reference of neuron IDs, cell types, transmitter type predictions and connection weights in different connectomes and the initial analysis of bi-synaptic pathways from DNa11 to motor neurons.

#### Data analysis for Fly Bowl experiments

The fly position and orientation were tracked frame by frame using the analysis pipeline described previously^110,111^. The raw data of angular velocity and forward velocity for each fly were first ‘lowess’ smoothed with a window of 5 frames (0.2 s) using the MATLAB function ‘smoothdata’. The absolute angular velocity (angular speed) and forward velocity 5 s before, during and after (in total 450 frames) each of the 9 optogenetic stimuli were averaged across the 9 stimuli for each fly (we pooled the data from the three intensity levels of optogenetic stimuli as they didn’t show obvious differences). These per fly data were further averaged across flies or across the time window of optogenetic stimuli (and subtracting the baseline which is 5 s before stimuli onset) to plot the time courses or quantify the light-induced changes in Fig. 1d-e and Fig. 1u-v. Turning events were detected in the entire recording session for each fly using the criteria of the angular velocity crossing a threshold of 45°/s. For each of these turning events, the average angular velocity was colour coded to plot Fig. S2a,g. The absolute translational velocity (ground speed) and angular velocity data across all flies of a genotype and 9 optogenetic stimuli in a 5-s time window either before or during the stimuli were pooled to calculate the distribution and plot Fig. S2b-c and Fig. S2h-i. The autocorrelation was performed on the angular velocity data (without smoothing) of each fly in each 5-s time window before or during the stimuli using the MATLAB function ‘crosscorr’. Per-fly data were first calculated by averaging across nine stimuli for each fly and then these per-fly data were used to calculate the mean and standard error across flies to plot Fig. S2d-f and Fig. S2j-l.

For the experiment in Fig. 2h-i, the two recording sessions were concatenated resulting in a total of 300-s light-on duration for each fly. Angular velocity data were partitioned into light-on and light-off (the same duration flanking the light-on periods) windows and averaged for each light condition. The flies were initially grouped by their labelling of the target neuron (the left, right, neither or both neurons labelled). The groups for labelling the left or right neuron were combined as a unilateral group (with the sign of angular velocity flipped for one group) and statistical tests were performed on the unilateral group across genotypes.

#### Saccade rate analysis for open arena assays

The fly tracking and saccade detection were performed using computational pipelines described previously^34^. Briefly, for each walking bout, the continuous wavelet transform of the angular velocity was computed using Gaussian wavelets. After the local maxima of the frequency signal and of the absolute value of the angular velocity were matched, the putative turns were filtered first to remove small local maxima that did not disrupt the forward velocity of the fly (variance of forward speed < 3mm/s), and then to roughly match the signal with a template obtained via PCA on a subset of very prominent, spike-like events. For each fly, saccade rate was then computed during active locomotion only (forward velocity > 2 mm/s). The number of saccades with absolute rotational velocity greater than 200 °/s at peak occurring during active locomotion, was divided by the total active time in seconds. Saccade rate was thus expressed in Hz. For activation experiments, saccades were counted in 400-ms windows before and after stimulation onset to calculate the changes in saccade rate. To plot the time course, for each fly, at each frame in a peri-stimulus window (from -0.5 s pre- to 1 s post-stimulus), the binary saccade score (0 or 1) was summed across trials and divided by the number of stimulus presentations.

#### Data analysis for miniature wind-tunnel assays

The analysis for Fig. 1i-m was performed in MATLAB using the pipeline described previously^29^. Position and orientation data were low-pass filtered at 2.5 Hz using a 2-pole Butterworth filter before further analysis. Angular velocity was calculated as the absolute value of the derivative of the filtered unwrapped orientation (i.e. orientation with phases corrected to be continuous beyond 0° or 360°) divided by the time interval of 20 ms and filtered by the criteria of ground speed no less than 1 mm/s as previously described^29^. Average angular velocity within a time window of 5 s immediately after odour offset was calculated for the same fly across trials of the same condition. The resultant data for each fly from light-on versus light-off conditions were compared using paired t-tests.

#### Data analysis for spherical treadmill walking assays

A custom MATLAB script was used to track the ball rotation post hoc using the top view images of the ball for all the optogenetic experiments with flies walking on the treadmill. Briefly, the MATLAB function ‘imregtform’ was used to calculate a linear transformation between the images in neighbouring frames for two small squares (50 by 50 pixels) at fixed positions on the ball. Considering the positions of the two squares on the ball and the linear transformations obtained, the ball rotation speed in three axes can be calculated. The high frame rate (200 fps) ensures that the ball moves only a few pixels in neighbouring frames and therefore a good linear transformation can almost always be found. A ball painted with black and white patterns was used to obtain simultaneous tracking data from this method and Fictrac^99^ for comparison. Instantaneous ball rotation speed aligned well between the two methods.

The fly’s angular velocity was calculated as the ball’s rotational velocity around the vertical axis in the opposite sign. The raw data were first smoothed using a moving average of 0.2 s and then averaged across 10 trials for each fly. These per fly data were used to calculate the mean and standard error across flies and plot the time series graphs in Fig. 1o and Fig. 2c,e. Such per fly data were averaged across a 2-s time window after the optogenetic stimuli onset for each fly to calculate the per fly mean and used to plot the quantifications across flies in Fig. 1p and Fig. 2c. The raw angular velocity data in a 2-s time window after the optogenetic stimuli onset were pooled across trials and flies for each neuron labelling category to plot the angular velocity distributions in Fig. S3. For Fig. 2f,g, the two 5-s windows of blue light on and those of red light only immediately after the blue light were used to calculate the per fly mean.

#### Data analysis for tethered flight assays

The videos of the top view (400 fps) were down-sampled to 40 fps to ensure each frame covered at least an entire wing-beat cycle. These videos were further processed to enhance the contrast and cropped to 480 × 480 pixels in MATLAB. A DeepLabCut^112^ model was trained using a dataset of manually annotated ground truth frames to label the wing hinges and the most anterior and posterior points at the outer edge of the wings (Supplementary Video 2). With the triangle of the three points for each wing, the wingbeat amplitude can be calculated as the internal angle at the vertex of the wing hinge. The difference of wingbeat amplitude between the left and right wings were averaged across 10 trials for each fly and used to calculate the mean and standard error across flies and plot the time series graphs in Fig. 1r and Fig. 2d. Such per-fly data were averaged across a 2-s time window after the optogenetic stimuli onset before subtracting the baseline (average across the 1-s time window before the optogenetic stimuli onset) to calculate the per fly mean and used to plot the quantifications across flies in Fig. 1s and Fig. 2d.

#### Data analysis for virtual reality assays (goal-directed navigation)

Bar position (0-360° in world coordinates; fly’s front is designated as 120°) and fly’s forward, sideways and angular velocity were sampled at 50 Hz. Virtual trajectories of flies considering the bar at a fixed position were calculated to determine whether the fly was performing menotaxis and plot Fig. 3m and Fig. S4g. Menotaxis bouts were extracted from the virtual trajectories using the criteria of continuous straight trajectories of at least 200 mm with no more than 25mm deviation, following ref^22^. Bar jumps that fell into a menotaxis bout but not within 5 s from either end of the bout and fulfilled a walking criterion (the fly walked faster than 1.5 mm/s in at least 20% of the time of a 60-s window centred at the bar jump) were considered menotaxis trials. For simplicity, we focused our analysis on 90° menotaxis trials. The bar position and fly locomotion data in a 60-s window centred at the bar jump of all 90° menotaxis trials (-90° trials flipped) were extracted and aligned to plot Fig. 3n. The circular mean of the aligned bar position data was calculated using the Circstat MATLAB toolbox^113^ to plot Fig. 3o. For each trial, a “time to return” was calculated as the time required for the fly to return the bar to within 30° from the pre-jump position (longer than 30 s counted as 30 s) to plot Fig. 3q. A per fly median of the “time to return” across trials was used to plot Fig. 3p. The baseline (the mean of the 30-s window before bar jump) subtracted angular speed and forward velocity were Gaussian smoothed in a 2-s window and averaged across trials to plot Fig. 3r-s.

#### Data analysis for virtual reality assays (tortuosity modulation in novel environment)

We focused our analysis on comparing walking behaviour during the first and second presentation of the disk stimulus. Trajectories were analysed using custom scripts written in python 3.10.13. For each fly and trial, we computed the mean translational velocity, the mean absolute rotational velocity and the path lengths as the accumulated movement. Further, we computed the local path tortuosity as the ratio of the path length and the total displacement within a moving window of 20 s, and then calculated the median per fly. For the statistical evaluation we used the python packages statsmodels and scipy to perform unpaired Wilcoxon ranked sums tests comparing the silenced and control genotypes for each tested driver line.

#### Data analysis for calcium imaging experiments

Ball tracking data (50 Hz) were resampled to match the sampling rate and synchronized with the calcium imaging (∼7 Hz or ∼55 Hz) using the ThorSync-registered timestamps for both imaging and behavioural data. For flight, the wing beat amplitude was extracted by custom MATLAB scripts post hoc from videos and synchronized with imaging data in the same way.

The maximal projections across optical slices in depth were calculated for both colour channels for volumetric imaging. Motion corrections were performed on those images using the NoRMCorre algorithm implemented in MATLAB^114^. A ratio image of GCaMP / mScarlet was obtained by image division. A free shape ROI was manually drawn for each neuron on the neurites in the mean image across time of the green / red ratio images using the MATLAB function ‘drawfreehand’. The ROIs of EPG neurons were determined by manually defining the centre, two semiaxes and the rotation of two images.roi.Ellipse objects for the outer and inner edges of ellipsoid body respectively and partitioning the space in between into 16 wedges with evenly distributed angles. The background grey values of blank regions were uniformly subtracted from images. Average intensity for each ROI was determined in each frame of the background subtracted images for both colour channels. A GCaMP / mScarlet ratio was calculated by dividing the average intensity of an ROI in green channel by red channel and designated as F_t_ for the frame t. Average F_t_ from the lowest 5% frames in a session was designated as F_0_. To standardize the neuronal activity across different trials and flies, we calculated a normalized ΔF / F_0_ as (F_t_ - F_0_) / (F_t_ - F_0_) _max_, in which (F_t_ - F_0_) _max_ was the maximum of ΔF observed for a ROI in a session. For the *LAL121, ExR7>jGCaMP7f and DNa03*, *DNa11>RSET-jGCaMP8m* datasets, we still recorded the red channel for quality control but only used the green channel to calculate the ΔF / F_0_ as we found sample movement was minimal in most cases and the division with red channel would introduce extra noise. For the *EPG, ExR7>jGCaMP8m* and *DNa03, DNa11>jGCaMP8m* datasets, we didn’t normalize the ΔF / F_0_ to [0 to 1] as that would introduce distortions on the ratios between different EPG ROIs or between DNa03 and DNa11 during walking or flight. All example traces were plotted at a 7-Hz sampling frequency, so the *DNa03/DNa11>RSET-jGCaMP8m* dataset imaged at 55 Hz were smoothed with an 8-frame window while other datasets were not smoothed.

For the heatmaps of the single-cell-type imaging data, neural activity and angular velocity were ‘lowess’ smoothed with a window of 2 s. Data were pooled across trials and flies of a genotype and partitioned by 100 bins for X and Y axes respectively, to plot the heatmap in Fig. 5a. For the 3-way correlation heatmaps of the *EPG, ExR7>jGCaMP8m* dataset, all three variables were smoothed with a moving window of 1 s. Data were pooled across trials or flies and partitioned by 50 by 50 bins for [0 1] range of EPG mean and PVA amplitude respectively, and average ExR7 activity was calculated within each bin. The same data were partitioned into 7 bins by EPG mean in the range [0.2 0.55] and for each bin the correlation coefficient between ExR7 activity and PVA amplitude was computed with the MATLAB function ‘corr’, which also gives the Pearson’s one-tailed *p* value for the null hypothesis of the population correlation coefficient ρ ≥ 0.

For the turning event-triggered analysis with angular velocity peaks, the 7-Hz imaging data without smoothing were used to best preserve the temporal structure. Peak detection was performed on the angular velocity data using the MATLAB function ‘findpeaks’ for both left and right turns using a ‘minPeakDistance’ parameter of 0.3 s to reduce the subpeaks during a turn to be included. For all the turning events falling into the bins of peak angular velocity across trials and flies in each genotype, time series of activity in both neurons and the angular velocity of the fly flanking the peak angular velocity in 10 s on each side were aligned to plot Fig. 5b and Fig. S8. The angular velocity at each peak and the average neuronal activity across 3 consecutive frames centred at the angular velocity peak were used to quantify the turning events in the plots on the right of Fig. S8. The 95% confidence interval was calculated as mean ± T-score * s.e.m, in which the T-score was calculated using the MATLAB function ‘tinv’. The LAL121 transients in Fig. 3i were detected by the same peak detection procedure but using neural signals as input and a threshold of 0.2 for both peak height and prominence.

For the cross-correlation analysis, time series were partitioned into 22-s segments (a typical 153-s session contains 7 segments) before the MATLAB function ‘xcorr’ was applied on the mean-subtracted data. An active-walking criteria of mean ground speed larger than 0.5 mm/s was applied to exclude quiescent segments for the correlation between bilateral neural activity and angular velocity in LAL121, DNa03 and DNa11. For flight experiments, as the flight durations were variable and very often shorter than 22 s, the segments were taken as continuous flight time within a session. The resultant cross-correlation for segments were averaged first for each fly then across flies for each genotype. For the single-cell-type imaging data, neural activity and angular velocity were ‘lowess’ smoothed with a window of 2 s before computing the cross-correlation. To compare between different neurons in Fig. 5d and Fig. S7e-f, the correlation coefficient or lag at peak correlation for each fly was used as data replicates for each genotype and 1-way ANOVA was performed. Only for the lag, genotype had a significant effect and post hoc multi-comparison with Tukey-Kramer method was performed between each pair using the MATLAB function ‘multcompare’ and the results were also used to plot the 95% intervals in Fig. 5d. Interpolation was performed on the average cross-correlation curves for each neuron in Fig. S7b, using MATLAB function ‘interp1’ in the ‘spline’ mode to get a smoothed curve and the lag at peak for each neuron was detected on this curve. For all other datasets, no smoothing or very light smoothing (0.14-s window for flight data) were applied before cross-correlation.

For the *DNa03/DNa11>RSET-jGCaMP8m* dataset, deconvolution of the calcium imaging data was performed using the deconvolution module in the CaImAn-MATLAB package^115^ on each neuron’s ΔF / F_0_. Welch’s method was used for the frequency domain analysis (Fig. S11h-i), which involves dividing signals into segments, windowing them, and averaging their spectral estimates. We used the MATLAB function ‘pwelch’ to compute the power spectral density estimate, ‘tfestimate’ for the magnitude and phase of the transfer function and ‘mscohere’ for the coherence. A 1024-point hamming window, a 512-point overlap and a 1024-point sampling for discrete Fourier transform were used as input parameters for all three functions on the neural activity data sampled at 55 Hz. For the analysis of transients (Fig. 5i), the ΔF / F_0_ data were z-scored (to account for the difference in the absolute ΔF / F_0_ between DNa03 and DNa11) and subtracted from a local baseline (moving 20-percentile of a 3-s window) before peak detection (with a height threshold of 2) was performed using MATLAB function ‘findpeaks’. The start and end of each transient were determined by searching downwards threshold-crossings from peaks. The linear filters (Fig. 5j and Fig. S12i-j) were computed following ref^8,9^ by computing the first-order Wiener kernel. Briefly, the kernels were computed in the frequency domain by taking the cross-spectrum of the input and response signals and normalizing it by the power spectrum of the input. The resulting filters were then smoothed using an exponential low-pass filter with a cutoff of 4 Hz to remove high-frequency noise, before transforming back to the time domain. The neural activity and angular velocity data were resampled to 100 Hz by interpolation and then divided into 8-s segments. The filters were computed on these segments before averaged.

#### Data analysis for electrophysiology

Data analysis was performed using MATLAB. Electrophysiology and treadmill data were downsampled to 400 Hz. Active bouts were defined as at least one of the three velocities being above threshold for a minimum duration of 250 ms. Thresholds were 10 °/s (rotational velocity) and 0.1 mm/s (forward and side velocity). If two active bouts were flanking an instance of inactivity shorter than 500 ms, both bouts and the inactivity in between were together considered as one single bout. For cross-correlation analysis, the initial 50 ms of each bout were removed to avoid correlations with potential state transitions in the membrane potential influencing the analysis. Chance level was calculated through bootstrapping: to preserve temporal dynamics, 100-ms segments of the electrophysiology data, approximating the average length of a step cycle^116,117^, were cut and shuffled. Then the cross-correlation analysis between this artificial electrophysiology dataset and the walking data was run. We repeated this procedure 1000 times to calculate the shuffled data. Cross-correlation peaks were calculated by detecting absolute extrema +/- 0.2 s around 0-s lag, if no extrema were detected within that boundary, data was excluded from lag analysis. To test whether the detected lags were significantly different from 0-s lag, we again applied a bootstrapping procedure (10000 samples, resampling with replacement from data), took the median of each sample to calculate the 95% confidence interval of the bootstrapped distribution and tested whether the interval included zero. The turning-event triggered alignments were performed on turns with minimal peak amplitude of 20 °/s, with ipsilateral or contralateral turns separated relative to the recorded neuron. These events were aligned with the peak angular velocity before membrane potential and angular velocity were averaged.

#### 3D leg joint kinematics

Using the DeepFly3D package^60^, a neural network for marking all the leg joints and tarsal tips in the 2D images from 6 different camera views were trained based on a dataset of manually annotated ground truth frames. The synchronized videos from 6 cameras for flies of DNa11 unilateral activation were analysed using this network. 3D triangulation was then performed based on the 2D annotations to generate a 3D reconstruction of all legs of the walking fly in DeepFly3D. The 2D annotations and the 3D reconstruction were used to generate a preview video (Supplementary Video 3) of the leg skeletons using a DeepFly3D function. The 3D reconstructions from 10 trials, each from an individual fly labelling either the left or right DNa11, were selected based on the tracking quality for further analysis. A Python script was used to align the 3D coordinates from different experiments and extract the stance and swing phase for each leg. First, the fly’s midlines determined by the average of the body-coxa joints across both body sides were aligned across experiments by rotating the coordinates around the raw and pitch axes. Then the median of tarsal tip positions across all time points for the two middle legs were levelled by rotating the coordinates around the roll axis. To determine the stance and swing phase, a PCA analysis was performed on the 3D positions of tarsal tips of each leg. The first component of the PCA should signify the main movement direction of the tarsal tip. Peak detections (both peaks and counter peaks) were performed on the first component of PCA to detect the extreme positions using the ‘find_peaks’ function from scipy.signal. The shorter lags between neighbouring peaks were assumed to be swing phases. A MATLAB script was used to calculate the average steps and plot the 3D graphs in Fig. 6b-c. We first segmented the steps for each leg based on the stance/swing timing. We then flipped the data from flies of left DNa11 labelling to make the turning directions uniform across flies and resampled the position of each joint during each stance phase with a uniform sample number for each stance phase. These resampled joint positions were averaged across experiments to plot the average steps in Fig. 6b-c.

#### Data analysis for the leg amputation and DNa06 activation experiments

A DeepLabCut model was trained to track the tarsal tips, antennae and posterior edges of the head in the videos of the top view. A MATLAB script was used to determine the swing and stance phase based on the tarsal tip data. For each leg, whether the main movement direction was in the X or Y axis was empirically determined. Then we detected peaks and counter peaks in the data for the main movement direction using the MATLAB function ‘findpeaks’. Those peaks were considered stance/swing transition points. The shorter lags between neighbouring peaks were assumed to be swing phases. The stepping frequency and tarsal tip trajectories were then calculated based on these stance/swing phase timing and positions for Fig. 6f-k. The data for flies of left DNa11 labelled were flipped so that they can be merged with the flies of right neuron labelled in the same cohort. As the wings partially blocked the view of legs in the posterior of the fly, tracking of the contralateral hind leg tarsal tips were less accurate than the other 5 legs. But as the extreme positions in a step are mostly outside the blind spot, the averaged stance movement vector for the contralateral leg largely reflected the contralateral hind leg’s movement direction observed from the raw videos.

The posterior edge of the head was used to determine the head direction change and plot Fig. S14a,e,g. Ball rotation data were obtained from the same top-view videos as described above to plot Fig. S14b-d,f,h.

#### Statistics

Statistical analyses were performed in GraphPad Prism10 (GraphPad Software, MA), MATLAB (Mathworks, MA) or Python. Significance level was 0.05 for all statistical tests and the *p* values were two-tailed when applicable, except for the directed Pearson’s correlation test in Fig. 4e. The *p* value summaries were described in the figures and figure legends. Details for each statistical test and the exact *p* value wherever applicable were described in Supplementary Table 3.

## Data and code availability

The datasets generated in the current study and the analysis code are available from the corresponding authors on request.

## Acknowledgements

We thank Masayoshi Ito, Kei Ito and Gerry Rubin for sharing *SS31864* before publication; Vivek Jayaraman, Rachel Wilson, Glenn Turner, Gwyneth Card, Gerry Rubin and BDSC for fly strains; Tom Clandinin, Clarissa Whitmire and Leandro Scholz for discussion and comments on the manuscript; Janelia FlyLight and FlyCore for technical assistance in generating the split-GAL4 lines; Janelia FlyEM for providing the access to the MANC connectome before publication; Flywire and the community for sharing the FlyWire connectome and cell type annotations; Samuel Kelly for technical assistance in the camera recording software and DeepFly3D implementation; Danielle Christesen, Rob Sullivan, QBI Media Kitchen, QBI Workshop, QBI Advanced Microscopy for technical assistance. C.G. is supported by HORIZON-MSCA-2022-PF grant 101108661. M.E.C. is supported by European Research Council Consolidator Grant ERC-2022-CoC-101088936. R.T. and H.H. were supported by the Emmy-Noether grant 513850388 from the Deutsche Forschungsgemeinschaft (DFG, German Research Foundation). This research was supported by ARC grants DP220103391 and DP260103520 to B.J.D and K.F.

## Author contributions

K.F., conceptualization, data acquisition, data analysis, supervision and funding acquisition. R.T., data acquisition and analysis under H.H.’s supervision. C.G and V.P., data acquisition and analysis under M.E.C.’s supervision. Y.Z. and K.A.v.H., data acquisition and analysis under K.N.’s supervision. M.K. and A.M., data acquisition under K.F.’s supervision. R.M., split-GAL4 screening. M.N.V.D.P. and B.v.S., help with visual display setup. B.J.D., supervision and funding acquisition. K.F. and B.J.D., writing, in coordination with all other authors.

---

**Fig. S1:**
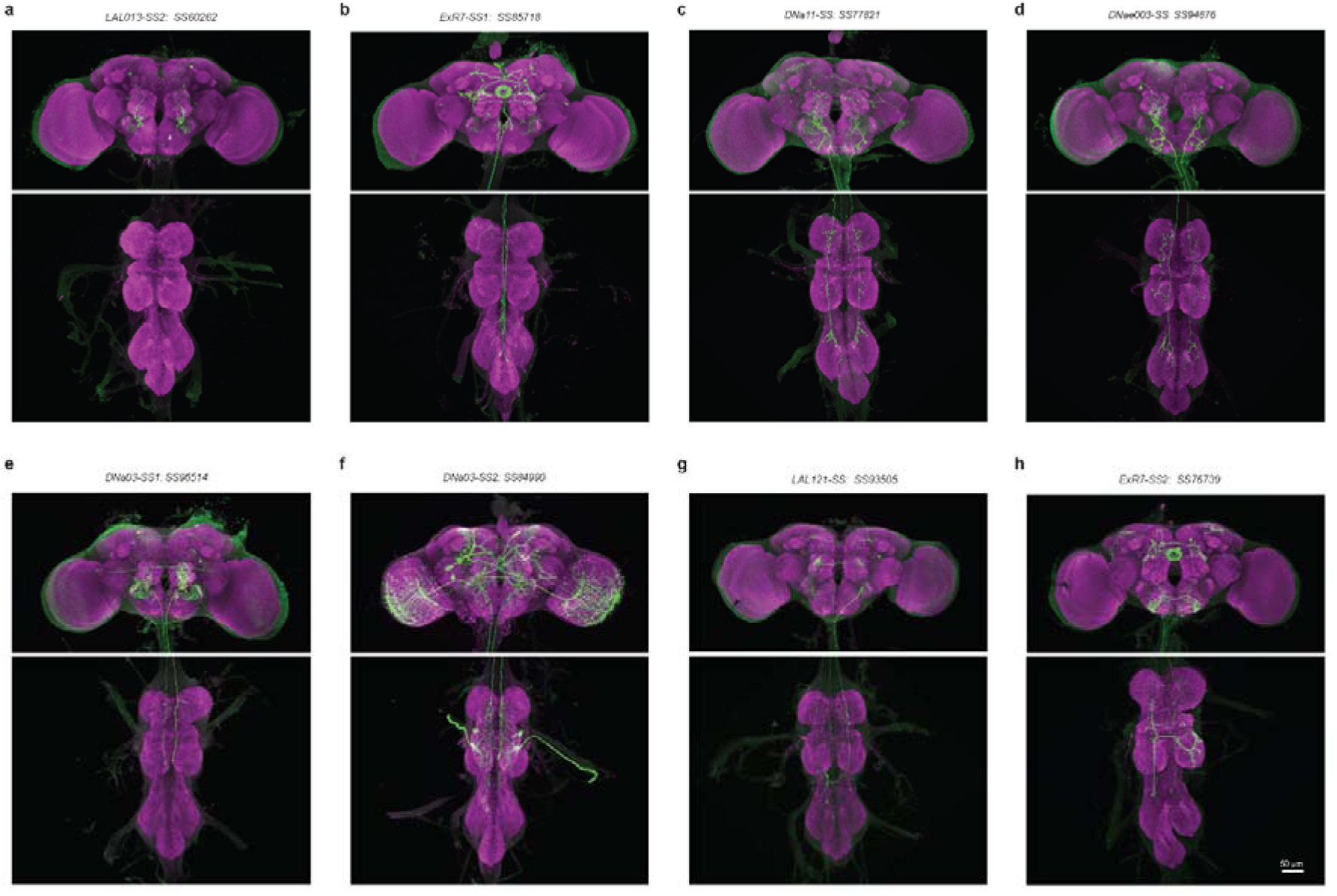
Expression pattern of *SS-GAL4* lines. **a-h**, Maximal projection of confocal stacks showing the expression pattern of each stable split-GAL4 line. Green, anti-GFP labelling to visualize mVenus reporter; magenta, nc82 labelling to visualize all synapses.

**Fig. S2:**
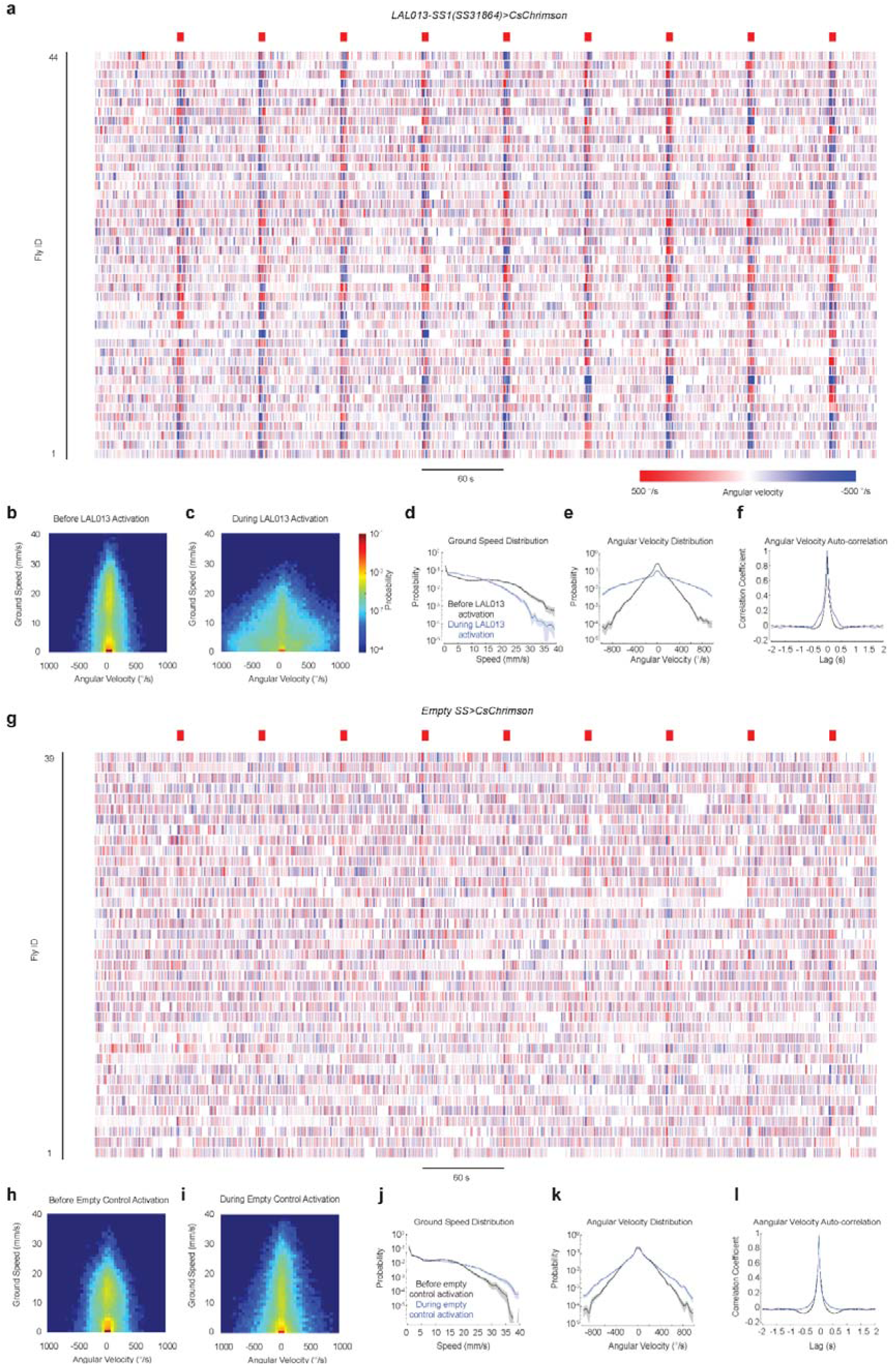
LAL013 bilateral activation elicits prolonged turning to an arbitrary direction and slower ground speed. **a**, Turning bouts (angular speed crossing a threshold of 45°/s) for each fly upon bilateral LAL013 activation. Red bars (top) indicate optogenetic stimuli. Average angular velocity during each turning bout is colour-coded. **b**, **c**, Distribution of ground speed and angular velocity in 5-second time windows before (b) and during (c) red-light stimulation of *LAL013-SS1>CsChrimson* flies. **d-f**, Ground speed distribution (d), angular velocity distribution (e) angular velocity autocorrelation (f) of *LAL013-SS1>CsChrimson* flies before (black) and during (blue) red-light stimuli. **g-l**, The same analyses for negative control flies (*Empty>CsChrimson)* as in (a-f). Data in (d-f) and (j-l) are mean ±Ls.e.m. across flies.

**Fig. S3:**
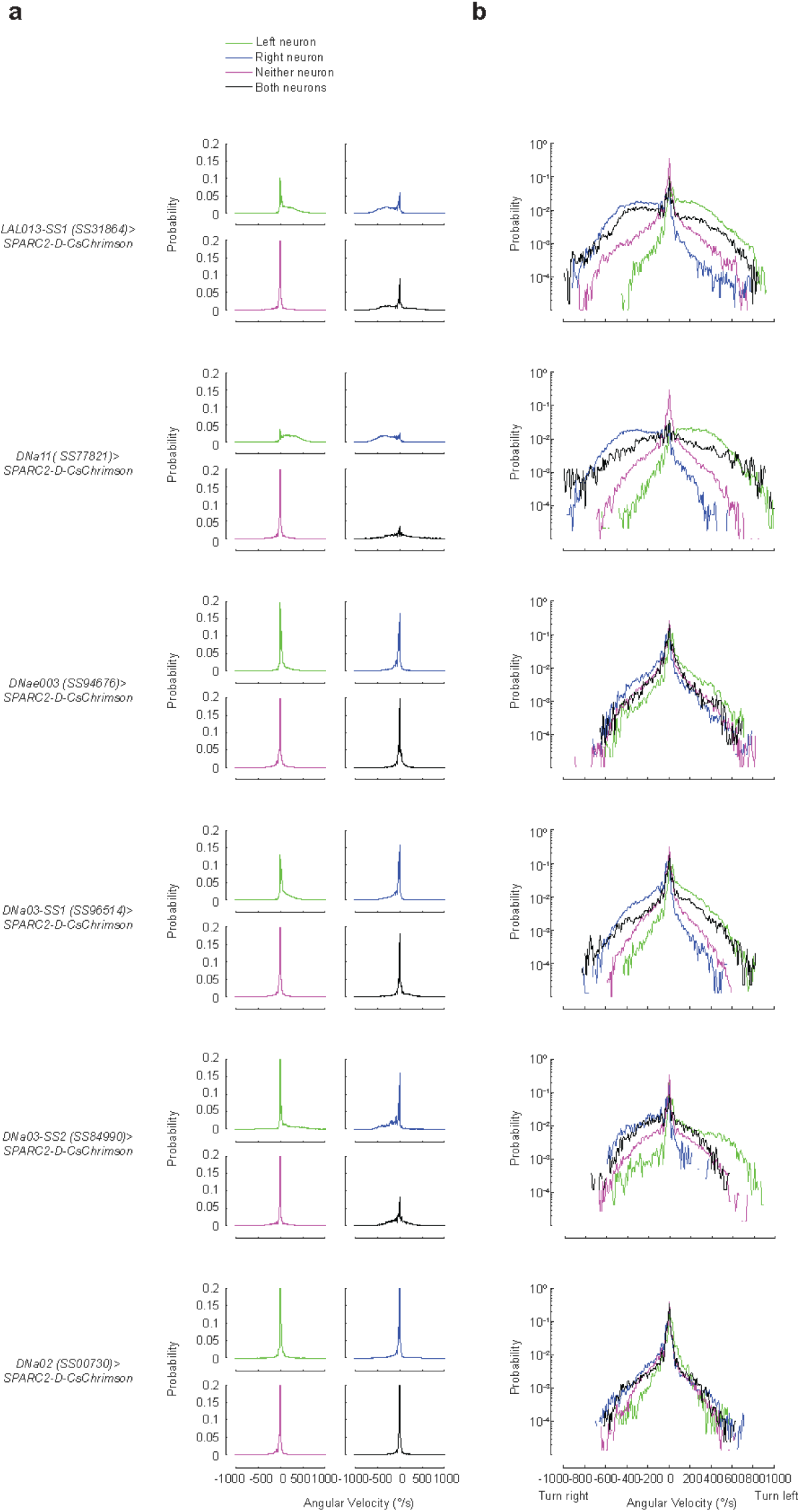
Angular velocity distribution during stochastic activation of steering neurons. **a**, Angular velocity distribution during the first 2 seconds of optogenetic activation, grouped by labelling of each cell type. **b**, Angular velocity distributions as in (a), plotted on a semi-logarithmic scale.

**Fig. S4:**
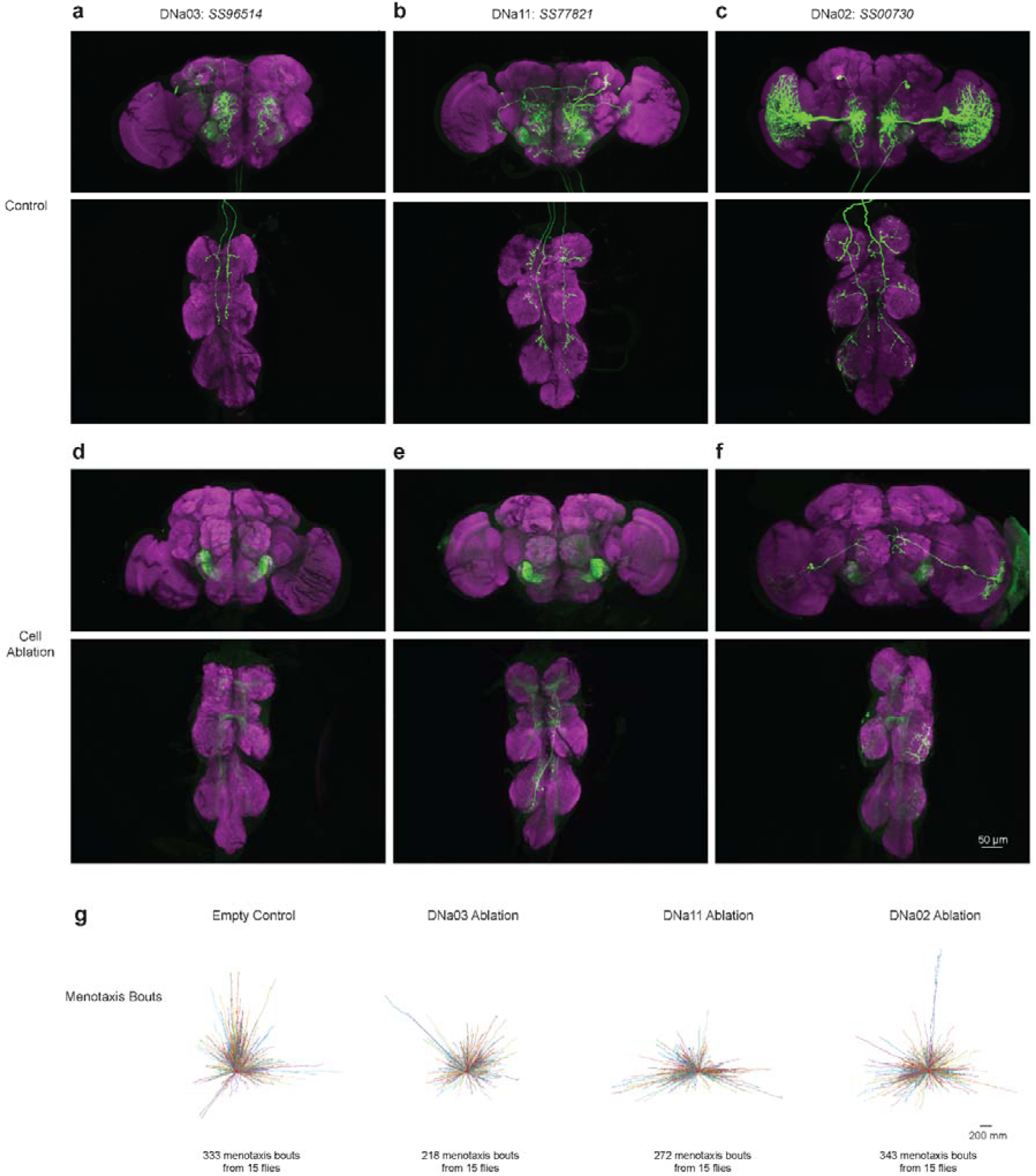
DNa03, DNa11 and DNa02 ablation in the menotaxis assay. **a-c**, Control images of *SS96514>CsChrimson-mVenus* labelling DNa03 (a), *SS77821>CsChrimson-mVenus* labelling DNa11 (b), and *SS00730>CsChrimson-mVenus* labelling DNa02 (c). **d**, Sample image of *SS96514>CsChrimson-mVenus, reaper, grim* to ablate DNa03. DNa03 was bilaterally ablated in 10/10 flies examined. **e**, Sample image of *SS77821>CsChrimson-mVenus, reaper, grim* to ablate DNa11. DNa11 was bilaterally ablated in 8/8 flies examined. **f**, Sample image of *SS00730>CsChrimson-mVenus, reaper, grim* to ablate DNa02. DNa02 was ablated in 8/8 flies examined. Note that the neuron-shaped signals in DNa11- and DNa02-ablated samples in (e) and (f) represent off-target cells co-labelled by the driver lines. **g**, Fictive trajectories of menotaxis bouts from control and DN-ablated flies, with the criteria of a minimal length of 200mm and a maximal deviation of 25mm from the course during each bout. Red dots indicate fly’s starting points. The bar is fixed at north (top on graph) to the fly’s starting point to compute the fictive trajectories.

**Fig. S5:**
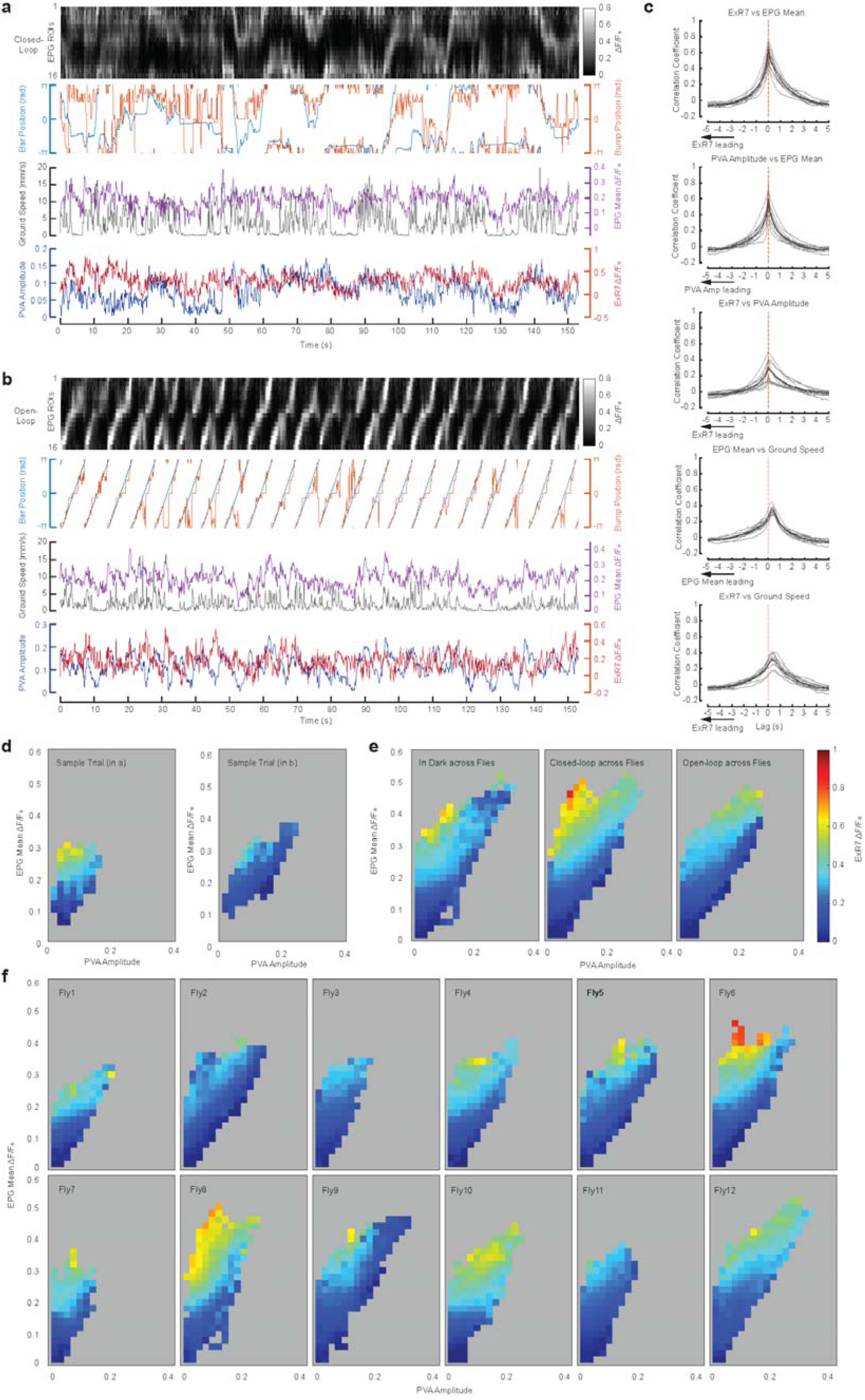
ExR7 activity correlates with uncertainty of heading estimate. **a**, Example traces of EPG, ExR7 dual imaging experiments, when the fly was in closed loop with a vertical bright bar. Bar and EPG Bump’s angular position shown in the second row. Bar position -120° to 0° were behind the fly and invisible. **b**, Same as (a) for a trial in open loop, where the bar’s movement is controlled by experimenter independent of fly’s locomotion. **c**, The cross-correlations between ExR7, EPG mean, PVA amplitude and ground speed. Each grey line represents one fly and the thick black line represents mean across flies (n = 12 flies). Note ExR7 and PVA amplitude have low and variable correlations, although either of the two correlates well with EPG mean. **d**, Colour-coded ExR7 activity binned by EPG mean and PVA amplitude, for the sample trials in (a-b). **e, f**, Same as (d), but grouped by three conditions across flies (e) or grouped by flies across conditions (f).

**Fig. S6.**
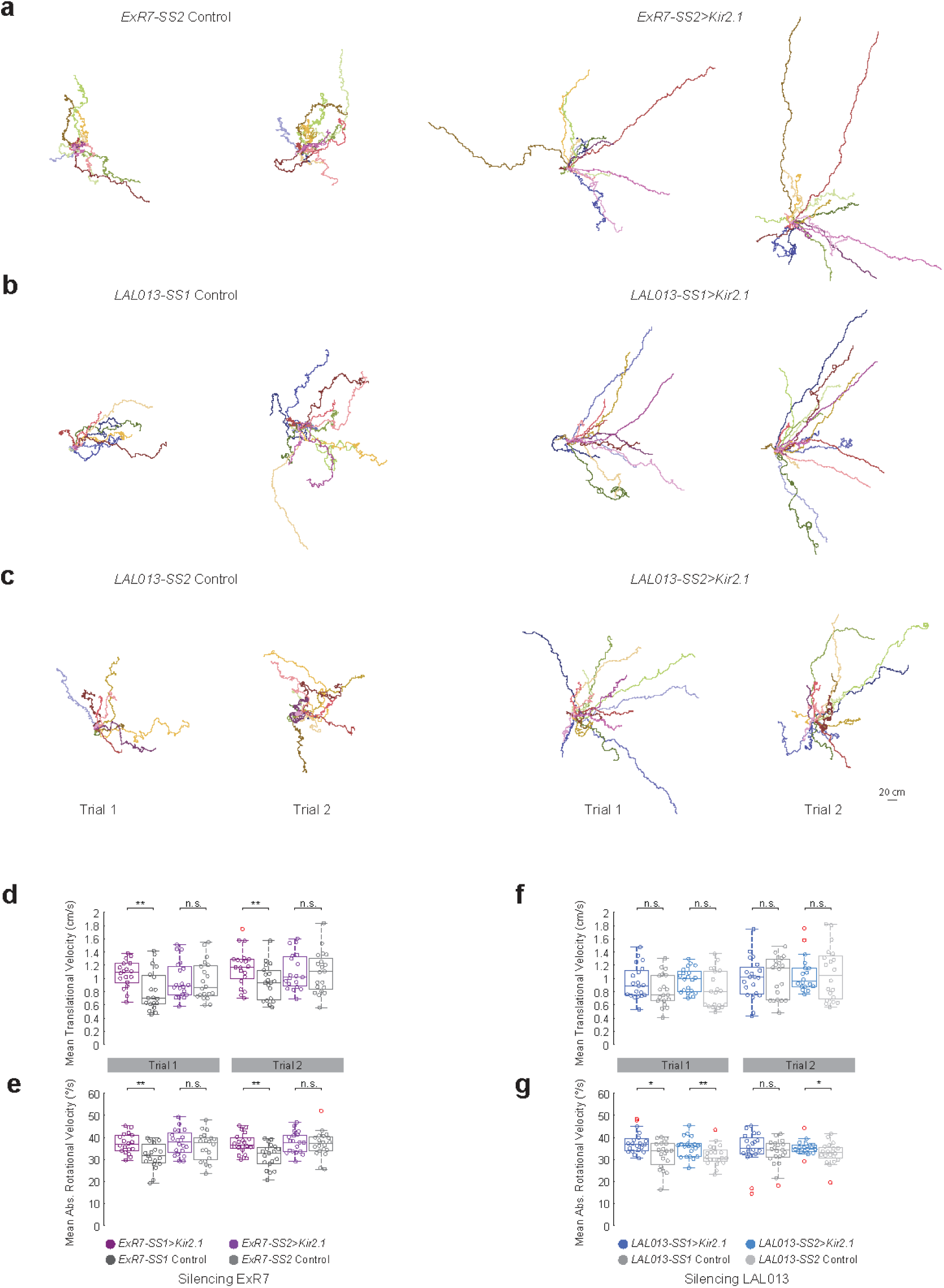
ExR7 and LAL013 silencing in a novel environment: trajectories and velocity data. **a**-**c**, Trajectories of individual flies in closed loop with sun disks. **d**-**g**, Boxplots depicting the mean translational velocity and absolute rotational velocity for ExR7 silencing and controls (d, e) and for LAL013 silencing and controls (f, g). The box shows the interquartile range (IQR). The line inside the box marks the median. Outliers are determined by boundaries of 1.5 × IQR from the first and third quartiles. Whiskers extend to the most remote non-outliner data points. **, *p*<0.01; *, *p*<0.05, n.s., *p*>0.05; Wilcoxon tests comparing the silenced and control genotypes.

**Fig. S7:**
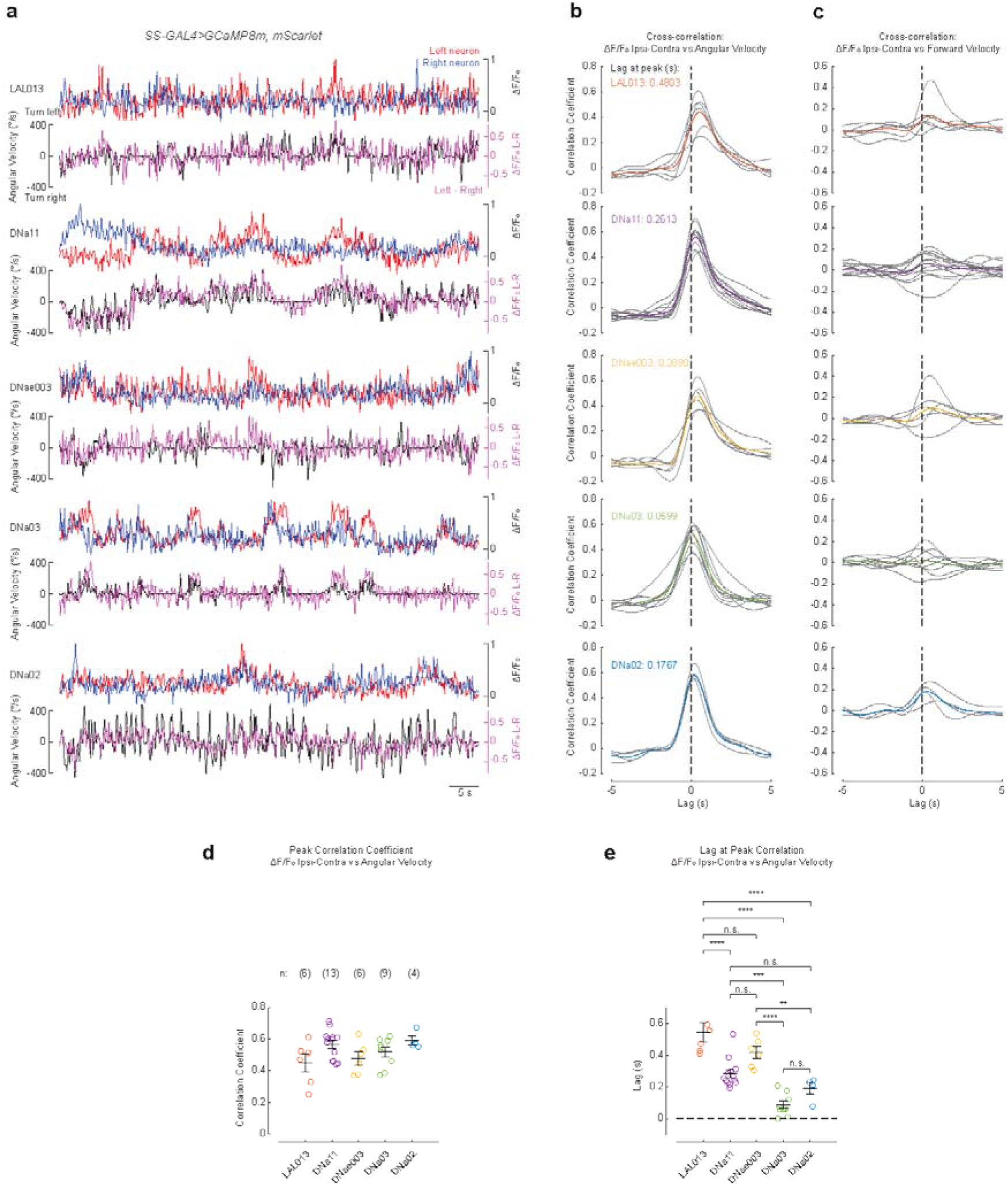
Steering neurons show different dynamics in activity during turning. **a**, Example traces for left and right neuron activity overlaid (red and blue lines in the top graphs) and the bilateral difference in neuronal activity overlaid with fly’s angular velocity (purple and black lines in the bottom graphs), for each cell type. **b**, Cross-correlation between bilateral differences of neural activity in 5 neuron types and angular velocity. Grey lines are means across all 22-s trials for each individual fly; thick coloured lines are mean across flies for each genotype. **c**, Same as (b) for forward velocity. **d**, Quantification of correlation coefficient at peaks. Not significant in 1-way ANOVA test. **e**, Quantification of lag at cross-correlation peaks. ****, *p*<0.0001; ***, *p*<0.001, **, *p*<0.01; n.s., *p*>0.05; Tukey-Kramer tests between each pair of neurons post-hoc to 1-way ANOVA.

**Fig. S8:**
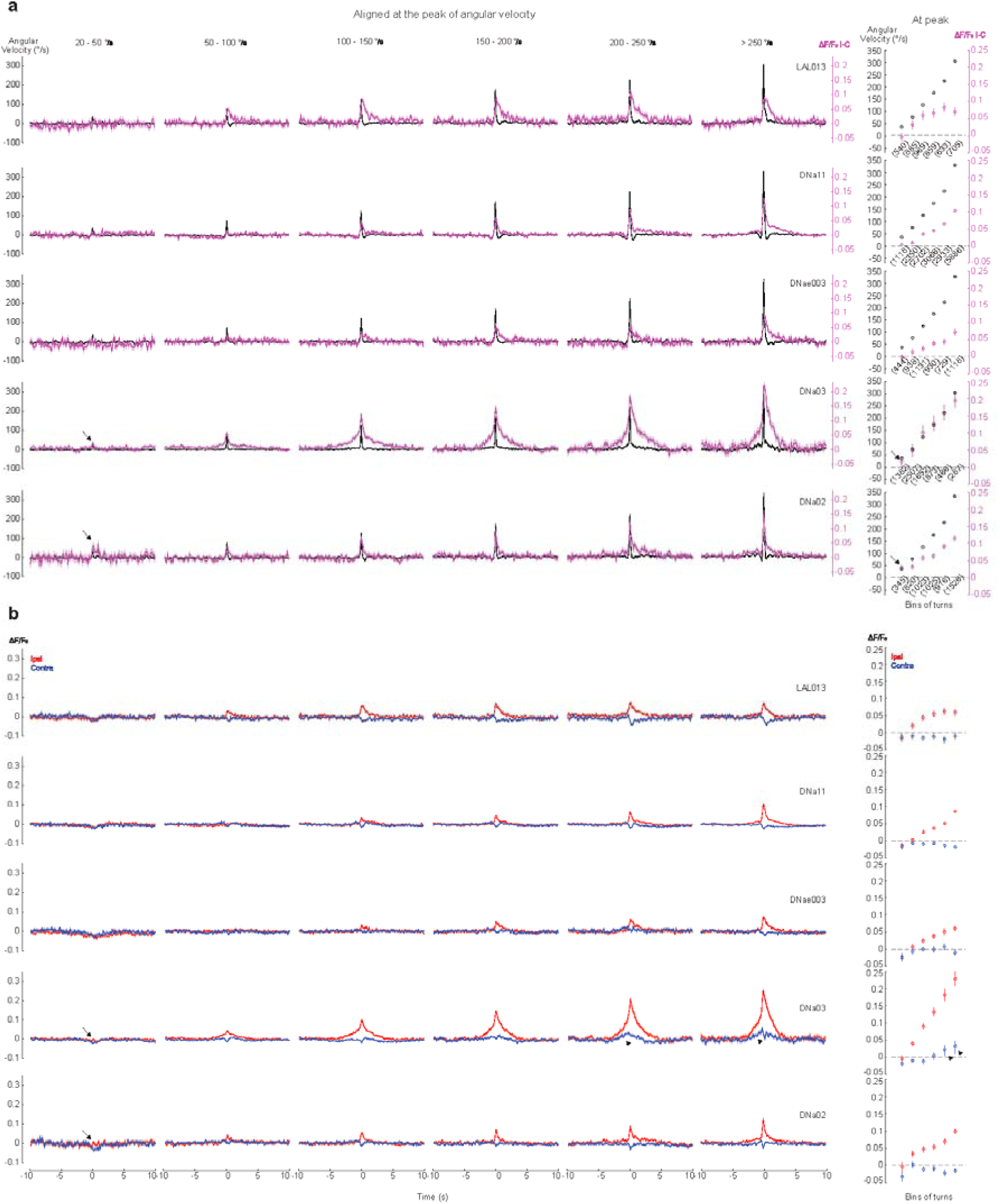
Angular velocity tuning in steering neurons. **a**, Left: angular velocity (black) and bilateral activity difference (purple) aligned at the peaks of angular velocity during turns, binned by the peak angular velocity. Data are mean ±Ls.e.m. across turning events. Right: average peak angular velocity (black) and bilateral activity difference at the peak angular velocity (purple) across turning events in each bin. Numbers in brackets are n of turning events in each bin. Error bars indicate 95% confidence intervals. **b**, Left: neuronal activity in ipsilateral (red) and contralateral (blue) neurons (relative to the turning direction) overlaid, as aligned in (a) for the same bins. Right: average activity in ipsilateral (red) and contralateral (blue) neurons at the peak angular velocity across the turning events in each bin. Arrows, only DNa03 and DNa02 showed responses during turns of 20-50°/s; arrowheads, the contralateral neuron of DNa03 ramps up activity before turns greater than 200°/s.

**Fig. S9:**
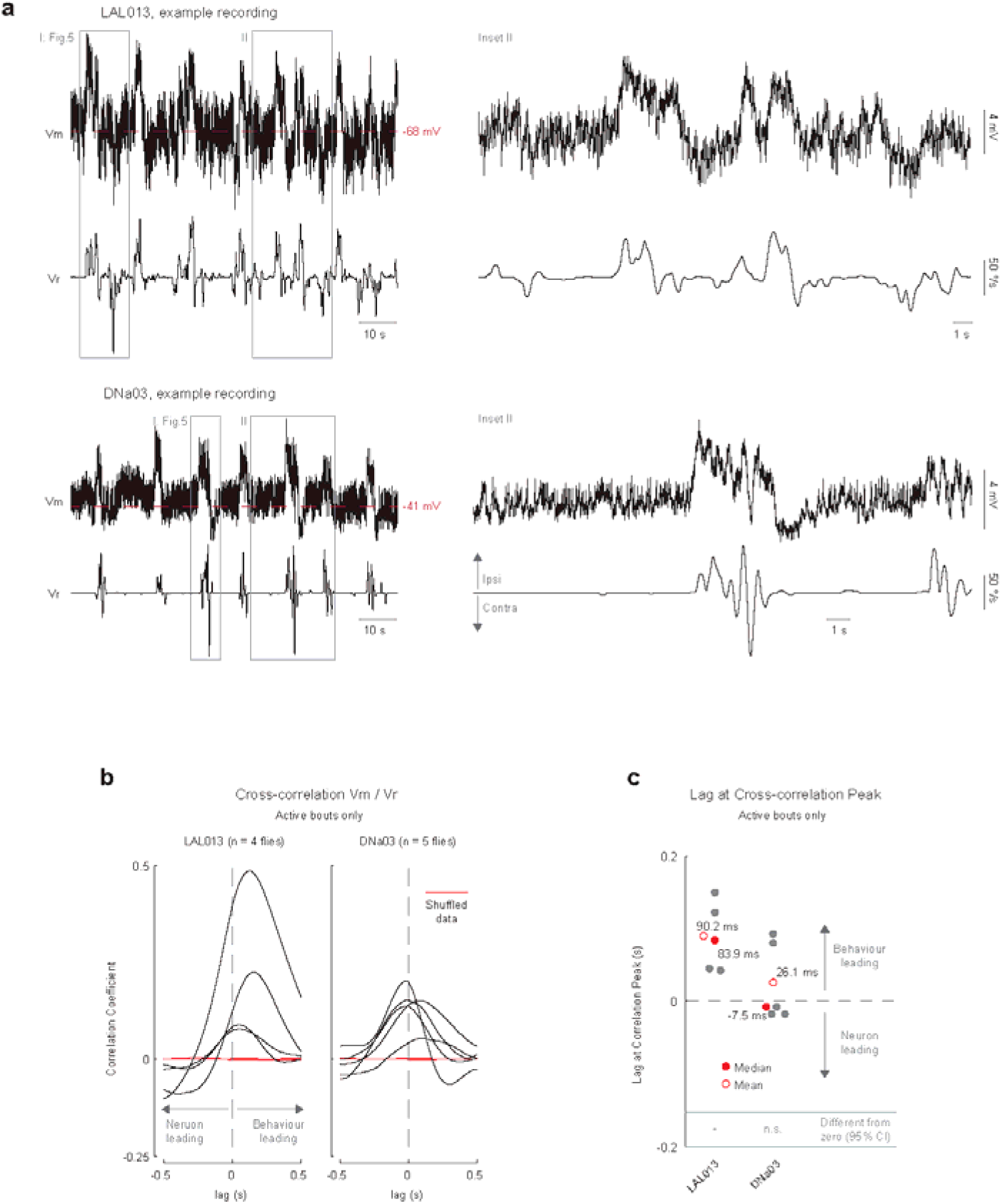
Patch clamp recordings of LAL013 and DNa03 neurons. **a**, Example traces of LAL013 and DNa03 recordings. **b**. Cross-correlation between membrane potentials and angular velocity during active walking bouts. Each black line represents a single fly. Red lines represent shuffled data. **c**. Lag at peak-correlation for both neurons. *, *p* < 0.05; n.s., *p* ≥ 0.05 for median different from 0 by bootstrapping (see Methods).

**Fig. S10:**
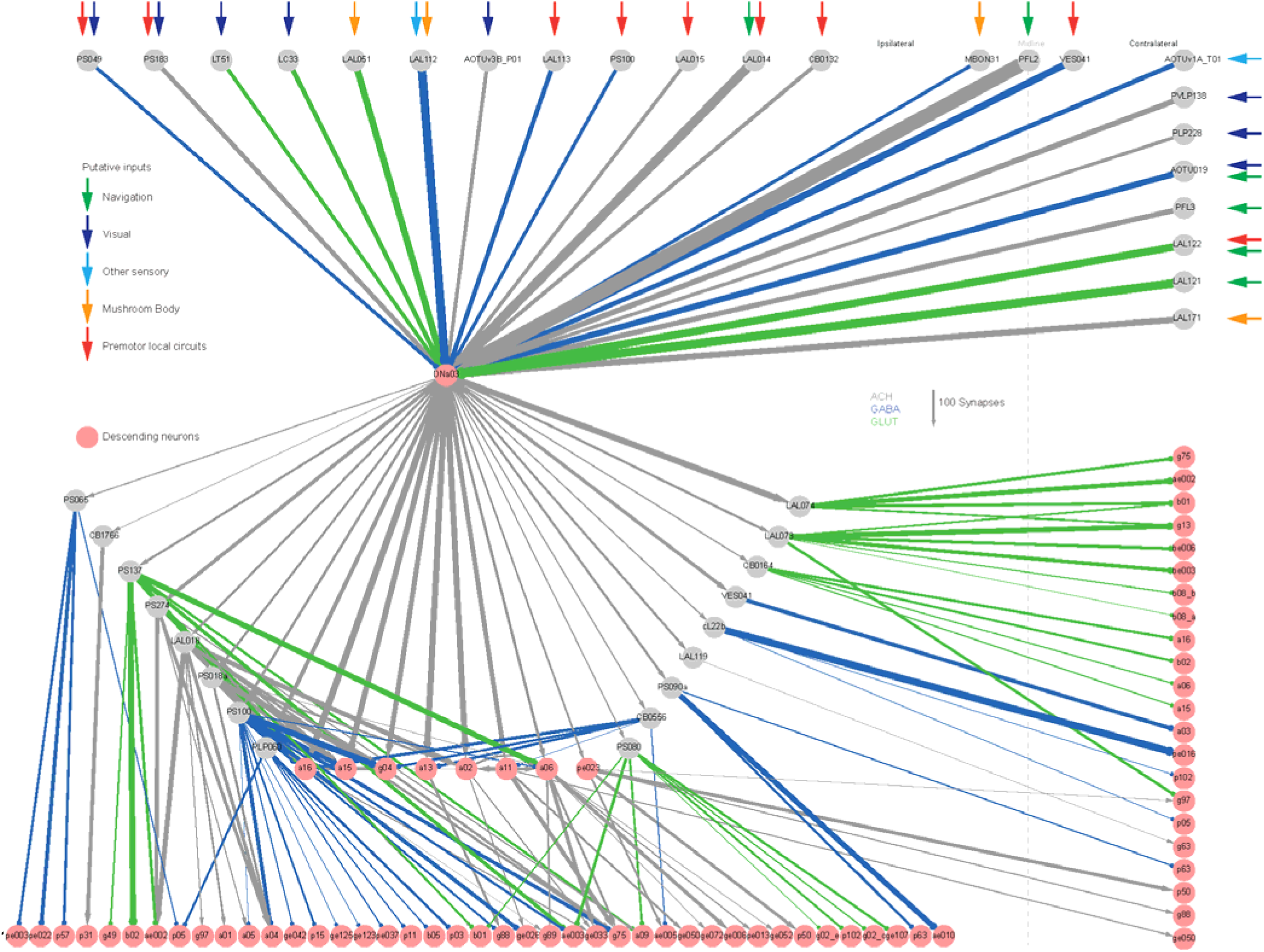
DNa03 upstream and downstream network. DNa03 upstream and downstream network, with synaptic weights from FlyWire. Edge colours represent predicted transmitter types and edge widths represent synapse counts. For upstream neurons, top direct inputs with synapse counts larger than 150 were shown and putative inputs to these neurons were manually annotated with reference to the connectome and literatures. For downstream neurons, direct and bi-synaptic connections to descending neurons (pink nodes) with synapse counts larger than 50 in each leg of connection were shown. See Methods for details.

**Fig. S11:**
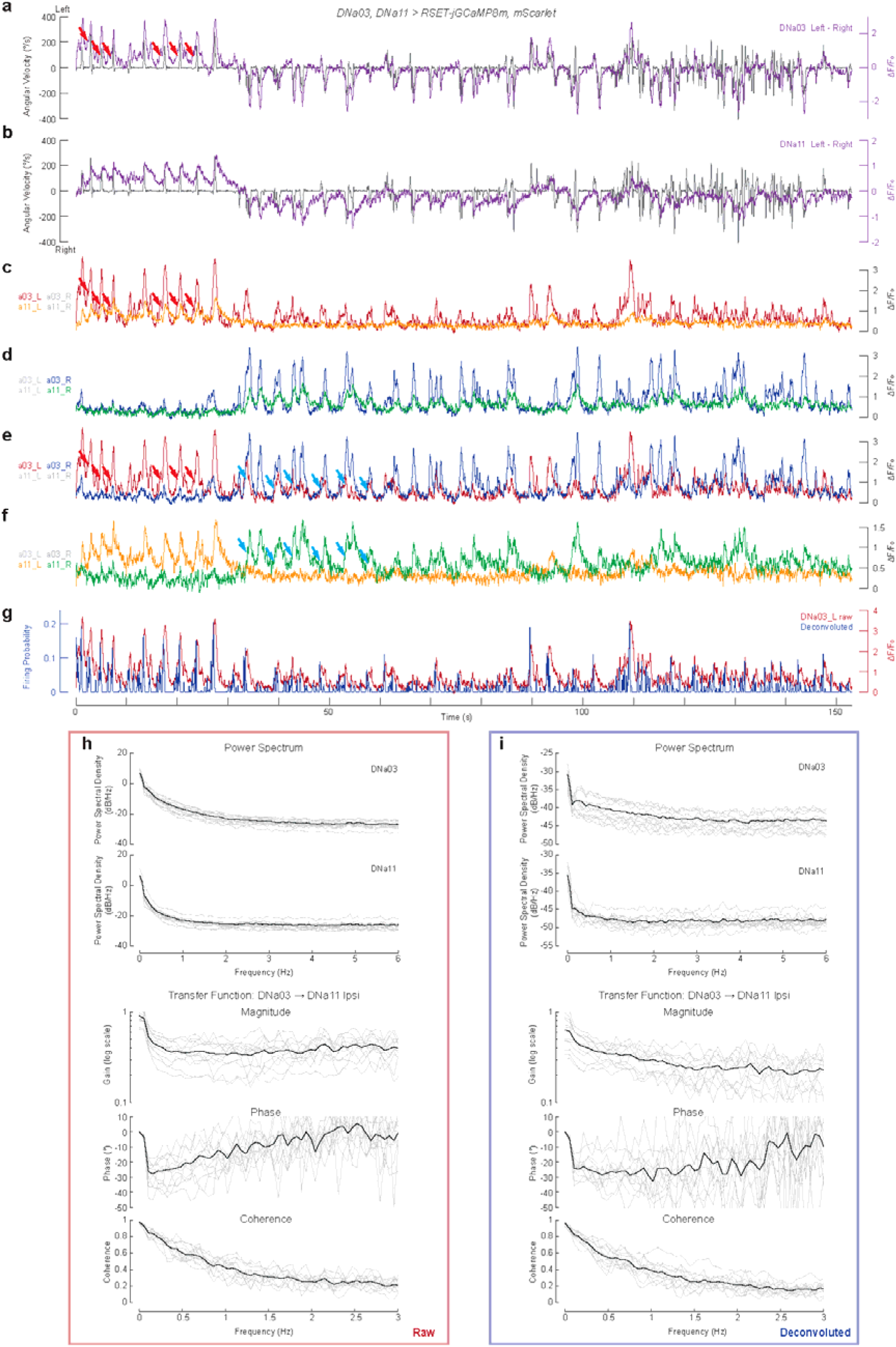
The transformation between DNa03 and DNa11 during walking. **a**, Example traces from DNa03, DNa11 dual imaging experiments showing DNa03 bilateral differences overlaid with angular velocity. Arrows indicate DNa03 ramping up activity prior to the angular velocity changes. **b**, Same as (a) for DNa11 in the same trial. **c**, **d**, Overlay of the ipsilateral DNa03 and DNa11 neurons on the left (c) or right (d). **e**, **f**, Overlay of both DNa03 neurons (e) or both DNa11 neurons (f). **g**, Example trace of deconvolved versus raw signal (left DNa03 neuron). The deconvolved signal represents putative spiking probability. **h**, **i**, Frequency domain analyses of DNa03-DNa11 transformation: the power spectra of both neurons and transfer function between the ipsilateral DNa03 and DNa11. The transfer function magnitude (in logarithmic scale), phase and coherence were shown from top to bottom. Grey lines represent single pairs of neurons and black lines represent mean across neuron pairs. N = 6 flies and n = 12 pair of neurons.

**Fig. S12:**
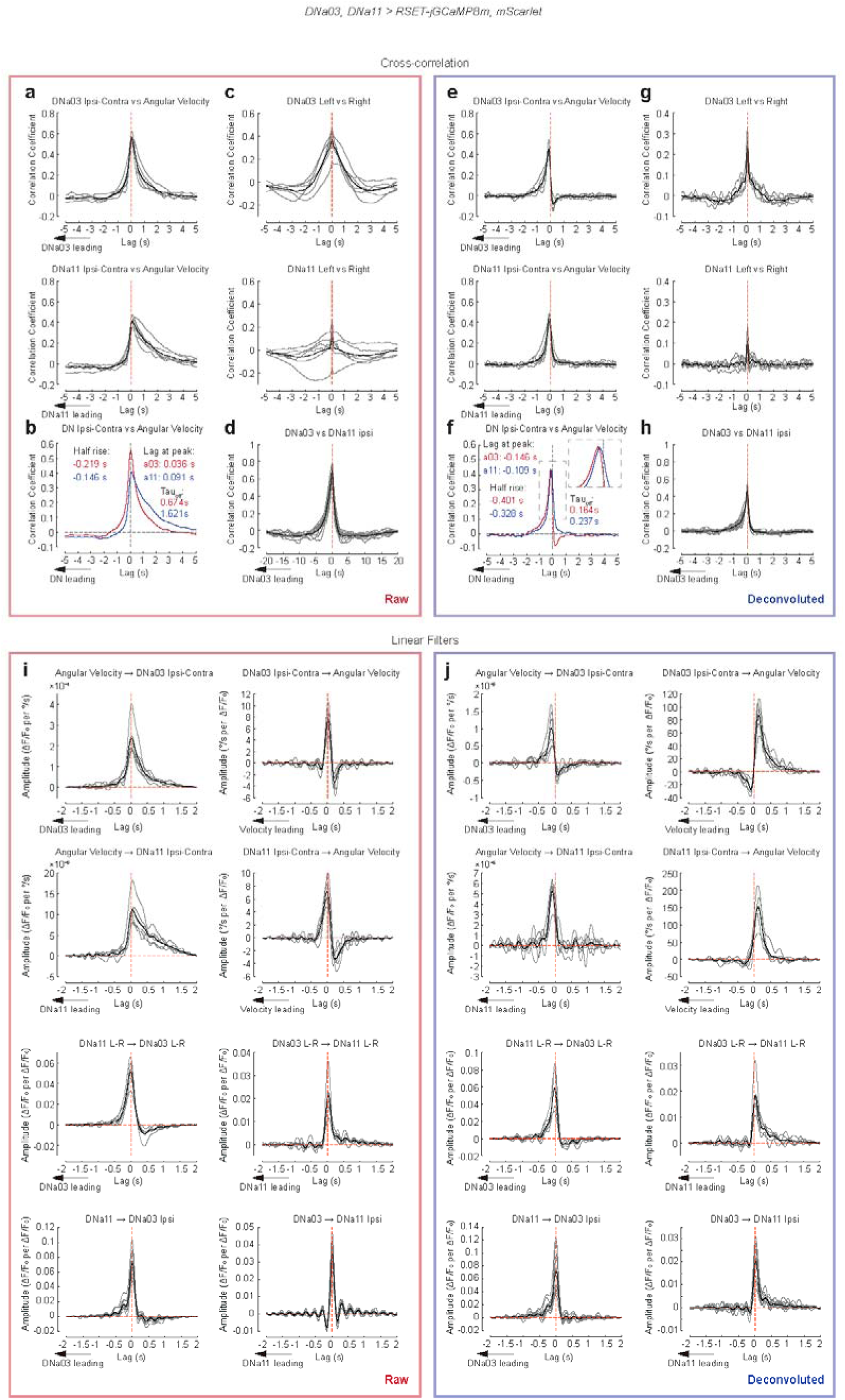
Cross-correlation and linear filters between DNa03, DNa11 and angular velocity in walking flies. **a**, Cross-correlation between bilateral differences in DNa03 (top) or DNa11 (bottom) activity and angular velocity. For each fly, 22-s segments of time series were used to calculate cross-correlation before averaging across segments. **b**, Overlay of the average cross-correlation across flies for DNa03 and DNa11 from (a). The lags at half rise or peak and the off time constant are shown on the graph. **c**, Cross-correlation between the left and right DNa03 (top) or DNa11 (bottom) neurons. Note the sharp peak at 0 present in both neurons is only 1-frame wide, likely imaging noise. **d**, Cross-correlation between DNa03 and ipsilateral DNa11 neurons. Note the skewed shape towards DNa03 leading. **e-h**, Same as (a-d), but for deconvolved neural activity. Note the scale differences between (h) and (d) in time. **i**, Linear filters for the inputs and outputs indicated in the title of each graph. Note the scales in time are different from cross-correlations. As Linear filters normalize the cross-correlation with input’s autocorrelation, the peaks are generally narrower. **j**, Same as (i) but for deconvolved neural activity signals. Grey lines represent individual flies (N = 6) or neuron pairs (n = 12). Thick black lines represent the mean across flies or neuron pairs.

**Fig. S13:**
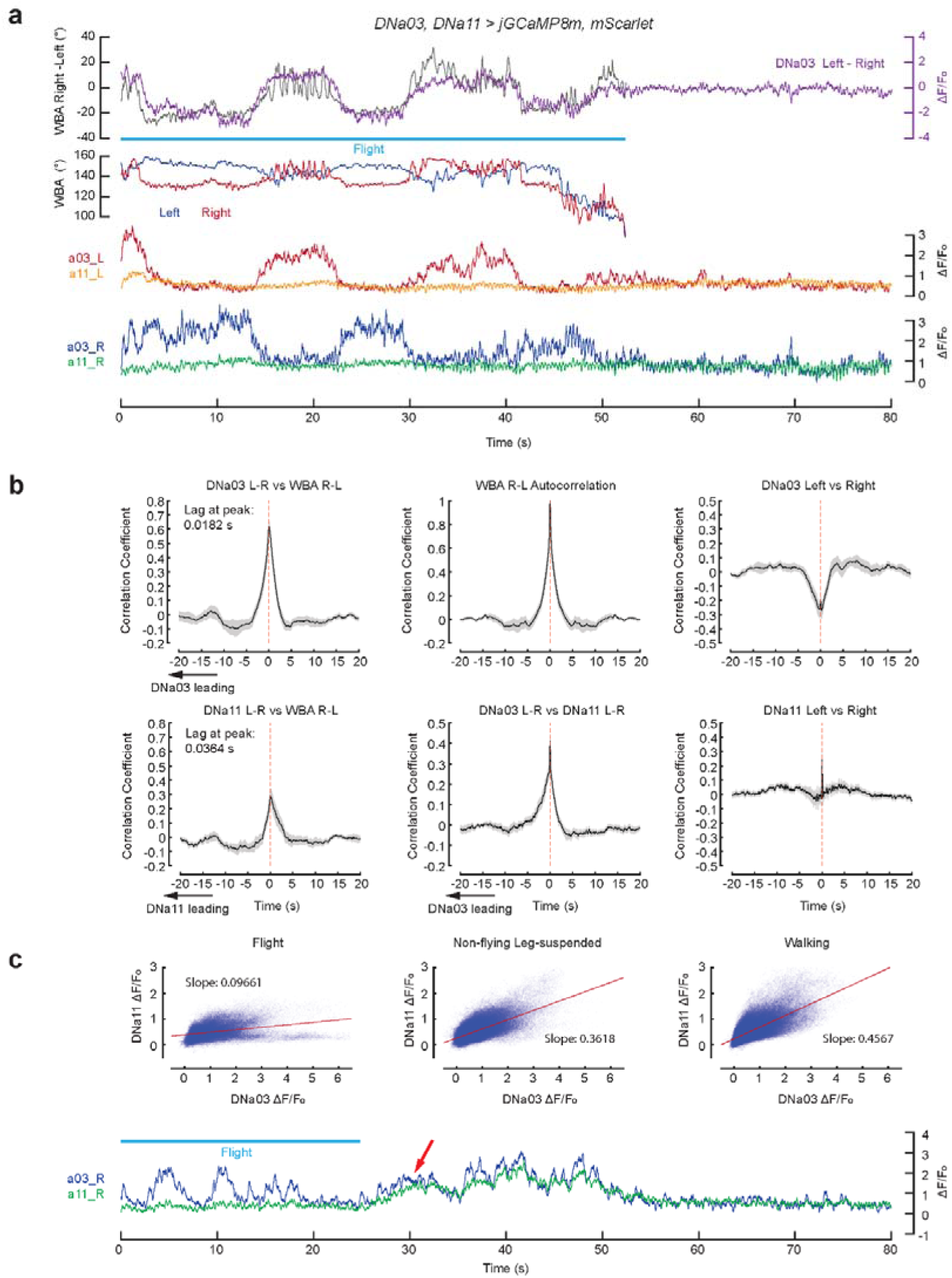
The transformation between DNa03 and DNa11 during flight. **a**, Example traces from DNa03, DNa11 dual imaging experiments during flight. **b**, Cross-correlations between variables as indicated in the graphs. The thick black line and grey shades represent mean ±Ls.e.m. across flies (n = 13). The peaks between bilateral differences in neuronal activity and angular velocity are wider than during walking, likely due to the wider autocorrelation of behaviour. Note the correlation between left and right DNa03 neurons was negative, different from the positive correlation seen during walking. **c**, Top: regression between DNa03 and ipsilateral DNa11 activity during flight, non-flying suspended state or walking. Fitted slopes are shown in graphs. Bottom: an example trace of DNa03 and DNa11 pair during flight. The light blue bar shows the period of flight. The arrow indicates that shortly after flight stopped, DNa11 started to trail DNa03, by contrast to the flattened trace during flight.

**Fig. S14:**
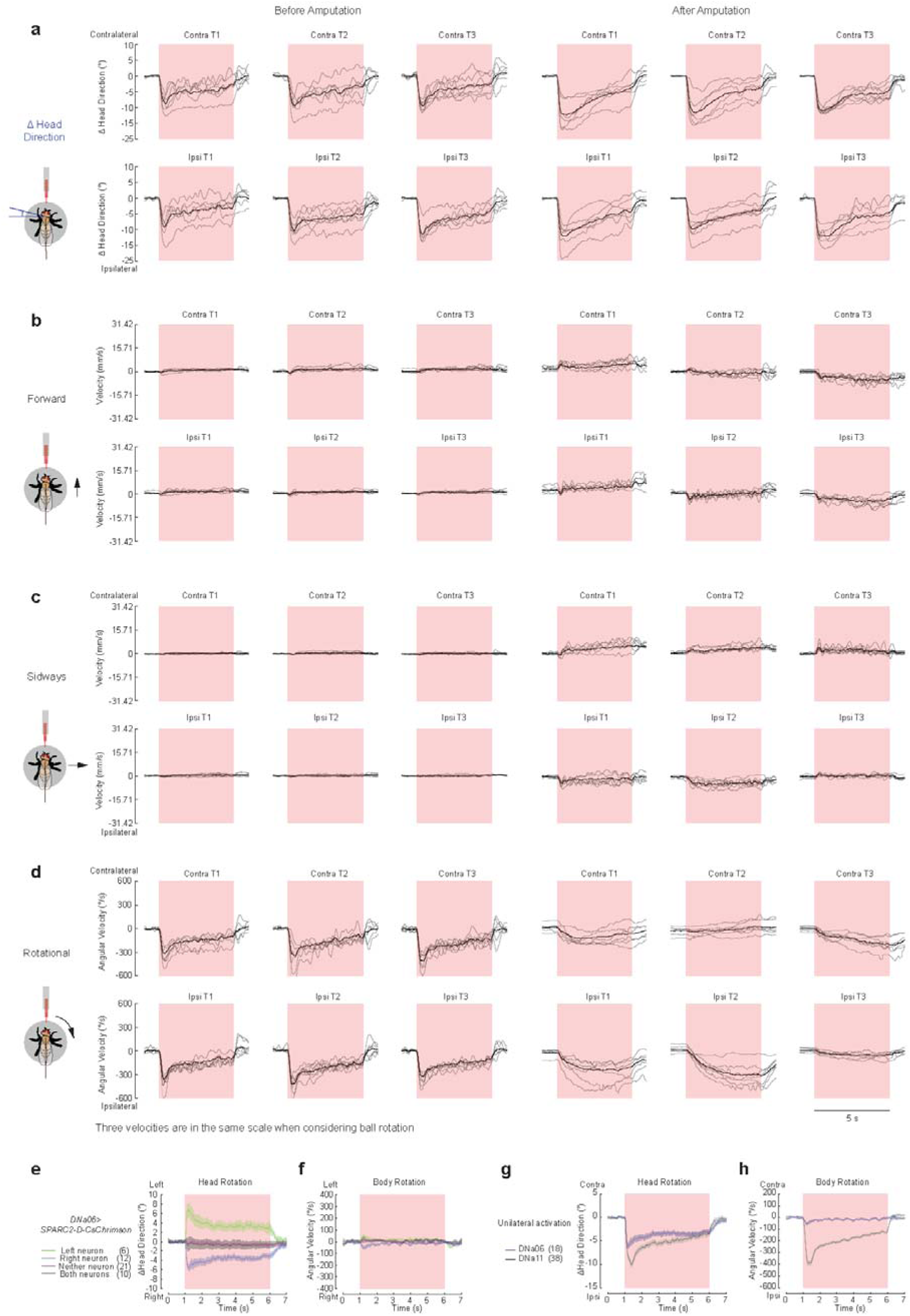
DNa11 controls coordinated head and leg movements during turning and DNa06 primarily controls head orientation. **a**, Head direction changes from the baseline (1s window before red-light onset), **b**, forward velocity, **c**, sideways velocity and **d**, rotational velocity, for each fly before (left) and after (right) leg amputations. Grey lines indicate the mean per fly across 10 trials. Black lines indicate the mean across flies. Red squares indicate optogenetic stimuli. The three velocities are in the same scale when considering ball rotation. **e**, **f**, The head rotation (e) and angular velocity (f) of *DNa06>SPARC2-D-CsChrimson* flies upon 5-s red light stimuli grouped by labelling of left, right, neither or both DNa06 neurons. **g**, **h**, The head rotation (g) and angular velocity (h) of flies with DNa06 (blue) or DNa11 (black) unilaterally activated. Data in (e-g) are mean ±Ls.e.m. across flies.

