## Supplementary Information and Tables for "Hierarchical Transformation of Navigational Variables into Coordinated Turning"

**Supplementary Table 1: The genotypes for each data item.**
The full genotypes of the flies used in each data item.

**Supplementary Table 2: Connectome analyses.**
The IDs and other information related to neurons included in the connectome analyses.

**Supplementary Table 3: Details of the statistical tests.**
Exact *p* values and other details of the statistical tests reported in this study.

**Supplementary Video 1: LAL013 bilateral activation elicits turning.**
Optogenetic activation of *SS31864>CsChrimson* flies (LAL013 bilaterally labelled). White indicator light at right bottom indicates the time of optogenetic stimulation. Recorded and played back at 30 fps.

**Supplementary Video 2: Unilateral DNa03 activation elicits ipsilateral flight turns.**
Optogenetic activation of a *SS96514>SPARC2-D-CsChrimson* fly with the right DNa03 labelled. Red squares on the top left indicates the time of optogenetic stimulation. The coloured dots indicate body parts labelled by a DeepLabCut model. Originally recorded at 400 fps; down sampled to and played back at 40 fps.

**Supplementary Video 3: Unilateral DNa11 activation elicits saccadic walking turns.**
Optogenetic activation of a *SS77821>SPARC2-D-CsChrimson* fly with the right DNa11 labelled. The top row are three views from the ipsilateral side and the middle row are three views from the contralateral side, relative to DNa11 labelling. DeepFly3D was used to track the joints (labelling in the top two rows) and reconstruct the 3D model displayed at the bottom row. Red squares on the top left indicates the time of optogenetic stimulation. Recorded at 200 fps and played back at 40 fps.

**Supplementary Video 4: Unilateral DNa11 activation before amputation.**
Optogenetic activation of six *SS77821>SPARC2-D-CsChrimson* flies with the left DNa11 labelled. These flies were assigned to six different cohorts for later amputations (the remaining leg as the text indicated). Red squares on the top left indicates the time of optogenetic stimulation. Recorded at 200 fps and played back at 40 fps.

**Supplementary Video 5: Unilateral DNa11 activation after amputation.**
Optogenetic activation of the same *SS77821>SPARC2-D-CsChrimson* flies with the left DNa11 labelled as shown in Supplementary Video S4, but after amputation of 5 legs (the remaining leg as the text indicated). Red squares on the top left indicates the time of optogenetic stimulation. Recorded at 200 fps and played back at 40 fps.

**Supplementary Video 6: Decapitated fly turning upon unilateral DNa11 activation.**
Optogenetic activation of a *SS77821>SPARC2-D-CsChrimson* fly with the right DNa11 labelled. Top view is from the ipsilateral side and bottom view is from the contralateral side, relative to DNa11 labelling. Red squares on the top left indicates the time of optogenetic stimulation. Recorded and played back at 200 fps. 8/8 decapitated flies examined turned towards the ipsilateral side upon unilateral DNa11 activation.

Supplementary Table 1

| Data items | Designation | Genotype |
|---|---|---|
| Fig. 1b | LM: LAL013 MCFO segmented | R57C10- Flp2::PEST in attP18, w / w; brp::Snap / +; pJFRC201-10XUASFRT >STOP > FRT-myr::smGFP-HA in VK00005, pJFRC240-10XUAS-FRT>STOP > FRT-myr::smGFP-V5-THS-10XUASFRT>STOP>FRT-myr::smGFP-FLAG in su(Hw)attP1 / VT002063-GAL4 in attP2 |
| Fig. 1d-e; Fig. S2a-f; Supp Video 1 | LAL013-SS1>CsChrimson | UAS-CsChrimson-mVenus in attP18, w / w; + / R83H09-AD; + / R37G12-DBD |
| Fig. 1d-e; Fig. S2g-l | Empty>CsChrimson | UAS-CsChrimson-mVenus in attP18, w / w; + / BPp65ADZp; + / BPZpGDBD |
| Fig. 1f | LAL013-SS2>CsChrimson | w / w; + / R83H09-AD; UAS-CsChrimson-mVenus in attP2 / VT002063-DBD |
| Fig. 1g | LAL013-SS2 Silencing | w / w; Otd-nls:FLPo / R83H09-AD; UAS>Stop>eGFPKir2.1 / VT002063-DBD |
| Fig. 1g | LAL013-SS2 Control | w / w; + / R83H09-AD; UAS>Stop>eGFPKir2.1 / VT002063-DBD |
| Fig. 1g-h | Empty Silencing | w / w; Otd-nls:FLPo / BPp65ADZp; UAS>Stop>eGFPKir2.1 / BPZpGDBD |
| Fig. 1h | LAL013-SS1 Silencing | w / w; Otd-nls:FLPo / R83H09-AD; UAS>Stop>eGFPKir2.1 / R37G12-DBD |
| Fig. 1h | LAL013-SS1 Control | w / w; + / R83H09-AD; UAS>Stop>eGFPKir2.1 / R37G12-DBD |
| Fig. 1i-m | LAL013-SS1>GtACR1 | w / NorpA $^{36}$, w; UAS-GtACR1-EYFP in attP40 / R83H09-AD, + / R37G12-DBD |
| Fig. 1i-m | LAL013-SS2>GtACR1 | w / NorpA $^{36}$, w; UAS-GtACR1-EYFP in attP40 / R83H09-AD, + / VT002063-DBD |
| Fig. 1i-m | Empty>GtACR1 | w / NorpA $^{36}$, w; UAS-GtACR1-EYFP in attP40 / BPp65ADZp, + / BPZpGDBD |
| Fig. 1n-s; Fig. S3a-b | LAL013-SS1>SPARC-D-CsChrimson | nSyb-PhiC31 in attP18, w / w; UAS-SPARC2-D-CsChrimson / R83H09-AD; + / R37G12-DBD |
| Fig. 1u-v | ExR7-SS1>CsChrimson | UAS-CsChrimson-mVenus in attP18, w / w; + / VT049727-AD; + / VT009668-DBD |
| Fig. 2a; Fig. 6a | DNa11 MCFO segmented | R57C10- Flp2::PEST in attP18, w / w; brp::Snap / R76B02-AD; pJFRC201-10XUASFRT >STOP > FRT-myr::smGFP-HA in VK00005, pJFRC240-10XUAS-FRT>STOP > FRT-myr::smGFP-V5-THS-10XUASFRT>STOP>FRT-myr::smGFP-FLAG in su(Hw)attP1 / VT040347-DBD |
| Fig. 2a | DNae003 MCFO segmented | R57C10- Flp2::PEST in attP18, w / w; brp::Snap / VT050660-AD; pJFRC201-10XUASFRT >STOP > FRT-myr::smGFP-HA in VK00005, pJFRC240-10XUAS-FRT>STOP > FRT-myr::smGFP-V5-THS-10XUASFRT>STOP>FRT-myr::smGFP-FLAG in su(Hw)attP1 / R22G01-DBD |
| Fig. 2a | DNa03 MCFO segmented | R57C10- Flp2::PEST in attP18, w / w; brp::Snap / VT025717-AD; pJFRC201-10XUASFRT >STOP > FRT-myr::smGFP-HA in VK00005, pJFRC240-10XUAS-FRT>STOP > FRT-myr::smGFP-V5-THS-10XUASFRT>STOP>FRT-myr::smGFP-FLAG in su(Hw)attP1 / VT028555-DBD |
| Fig. 2c; Fig. 6b-k; Fig. S3a-b; Fig. S14a-d,g-h; Supp Videos 3-6 | DNa11>SPARC2-D-CsChrimson | nSyb-PhiC31 in attP18 / w; UAS-SPARC2-D-CsChrimson in attP40 / R76B02-AD; + / VT040347-DBD |
| Fig. 2c; Fig. S3a-b | DNae003>SPARC2-D-CsChrimson | nSyb-PhiC31 in attP18 / w; UAS-SPARC2-D-CsChrimson in attP40 / VT050660-AD; + / R22G01-DBD |
| Fig. 2c-d; Fig. S3a-b; Supp Video 2 | DNa03-SS1>SPARC2-D-CsChrimson | nSyb-PhiC31 in attP18 / w; UAS-SPARC2-D-CsChrimson in attP40 / VT025717-AD; + / VT028555-DBD |

| | | |
|---|---|---|
| Fig. S3a-b | *DNa03-SS2>SPARC2-D-CsChrimson* | *nSyb-PhiC31 in attP18 / w; UAS-SPARC2-D-CsChrimson in attP40 / VT033616-AD; + / VT025718-DBD* |
| Fig. 2c; Fig. S3a-b | *DNa02>SPARC2-D-CsChrimson* | *nSyb-PhiC31 in attP18 / w; UAS-SPARC2-D-CsChrimson in attP40/ R75C10-AD; + / R87D07-DBD* |
| Fig. 2e-g | *LAL013-LexA>CsChrimson, Empty>GtACR2* | *LexAop2-CsChrimson-tdT in attp18, w / w; VT002063-LexAGADfl in attP40 / BPp65ADZp; UAS-GtACR2 in attP2 / BPZpGDBD* |
| Fig. 2e-g | *LAL013-LexA>CsChrimson, LAL013-SS1>GtACR2* | *LexAop2-CsChrimson-tdT in attp18, w / w; VT002063-LexAGADfl in attP40 / R83H09-AD; UAS-GtACR2 in attP2 / R37G12-DBD* |
| Fig. 2e-g | *LAL013-LexA>CsChrimson, DNa11>GtACR2* | *LexAop2-CsChrimson-tdT in attp18, w / w; VT002063-LexAGADfl in attP40 / R76B02-AD; UAS-GtACR2 in attP2 / VT040347-DBD* |
| Fig. 2e-g | *LAL013-LexA>CsChrimson, DNae003>GtACR2* | *LexAop2-CsChrimson-tdT in attp18, w / w; VT002063-LexAGADfl in attP40 / VT050660-AD; UAS-GtACR2 in attP2 / R22G01-DBD* |
| Fig. 2e-g | *LAL013-LexA>CsChrimson, DNa03-SS1>GtACR2* | *LexAop2-CsChrimson-tdT in attp18, w / w; VT002063-LexAGADfl in attP40 / VT025717-AD; UAS-GtACR2 in attP2 / VT028555-DBD* |
| Fig. 2e-g | *LAL013-LexA>CsChrimson, DNa02>GtACR2* | *LexAop2-CsChrimson-tdT in attp18, w / w; VT002063-LexAGADfl in attP40 / R75C10-AD; UAS-GtACR2 in attP2 / R87D07-DBD* |
| Fig. 2h | *DNa03>SPARC2-D-GtACR1* | *nSyb-PhiC31 in attP18 / w; UAS-SPARC2-D-GtACR1 in attP40 / VT025717-AD; + / VT028555-DBD* |
| Fig. 2h | *DNa03>SPARC2-I-GtACR1* | *nSyb-PhiC31 in attP18 / w; UAS-SPARC2-I-GtACR1 in attP40 / VT025717-AD; + / VT028555-DBD* |
| Fig. 2h | *DNa11>SPARC2-D-GtACR1* | *nSyb-PhiC31 in attP18 / w; UAS-SPARC2-D-GtACR1 in attP40 / R76B02-AD; + / VT040347-DBD* |
| Fig. 2h | *LAL013>SPARC2-D-GtACR1* | *nSyb-PhiC31 in attP18 / w; UAS-SPARC2-D-GtACR1 / R83H09-AD; + / R37G12-DBD* |
| Fig. 2h | *LAL013>SPARC2-I-GtACR1* | *nSyb-PhiC31 in attP18 / w; UAS-SPARC2-I-GtACR1 / R83H09-AD; + / R37G12-DBD* |
| Fig. 3b-i | *LAL121, ExR7 > Syt-jGCaMP7f, mScarlet* | *UAS-mScarlet in su(Hw)attP8, UAS-mScarlet in JK16L, w / w; VT049727-AD, UAS-IVS-Syn21-Synaptotagmin opGCaMP7f-p10 in JK22C / R65H04-AD; VT009668-DBD / VT061910-DBD, UAS-IVS-Syn21-Synaptotagmin opGCaMP7f-p10 in VK00005* |
| Fig. 3m-s; Fig. S4g | Empty control | *UAS-CsChrimson-mVenus in attP18, w / w; UAS-grim in su(Hw)attP6 / BPp65ADZp; UAS-reaper / BPZpGDBD* |
| Fig. 3n-s; Fig. S4d,g | DNa03 ablation | *UAS-CsChrimson-mVenus in attP18, w / w; UAS-grim in su(Hw)attP6 / VT25717-AD; UAS-reaper / VT028555-DBD* |
| Fig. 3n-s; Fig. S4e,g | DNa11 ablation | *UAS-CsChrimson-mVenus in attP18, w / w; UAS-grim in su(Hw)attP6 / R76B02-AD; UAS-reaper / VT040347-DBD* |
| Fig. 3n-s; Fig. S4f-g | DNa02 ablation | *UAS-CsChrimson-mVenus in attP18, w / w; UAS-grim in su(Hw)attP6 / R75C10-AD; UAS-reaper / R87D07-DBD* |

Supplementary Table 1

| | | |
|---|---|---|
| Fig. 4a-f; Fig. S5a-f | *EPG, ExR7 > jGCaMP8m, mScarlet* | *UAS-IVS-jGCaMP8m in su(Hw)attP8, UAS-mScarlet in JK16L, w / 20XUAS-IVS-jGCaMP8m in su(Hw)attP8, UAS-mScarlet in JK16L, w; R19G02-AD / VT049727-AD; R15C03-DBD / VT009668-DBD* |
| Fig. 4i-k; Fig. S6d-e | *ExR7-SS1>Kir2.1* | *w / +; VT049727-AD / +; VT009668-DBD / UAS-Kir2.1 in WTB background* |
| Fig. 4i-k; Fig. S6d-e | *ExR7-SS1* Control | *w / +; VT049727-AD / +; VT009668-DBD / + in WTB background* |
| Fig. 4j-k; Fig. S6a,d,e | *ExR7-SS2>Kir2.1* | *w / +; R65A07-AD / +; VT049427-DBD / UAS-Kir2.1 in WTB background* |
| Fig. 4j-k; Fig. S6a,d,e | *ExR7-SS2* Control | *w / +; R65A07-AD / +; VT049427-DBD / + in WTB background* |
| Fig. 4l-m; Fig. S6b,f-g | *LAL013-SS1>Kir2.1* | *w / +; R83H09-AD / +; R37G12-DBD / UAS-Kir2.1 in WTB background* |
| Fig. 4l-m; Fig. S6b,f-g | *LAL013-SS1* Control | *w / +; R83H09-AD / +; R37G12-DBD / + in WTB background* |
| Fig. 4l-m; Fig. S6c,f-g | *LAL013-SS2>Kir2.1* | *w / +; R83H09-AD / +; VT002063-DBD / UAS-Kir2.1 in WTB background* |
| Fig. 4l-m; Fig. S6c,f-g | *LAL013-SS2* Control | *w / +; R83H09-AD / +; VT002063-DBD / + in WTB background* |
| Fig. 5a-d; Fig. S7a-e; Fig. S8a-b | *LAL013>jGCaMP8m, mScarlet* | *UAS-jGCaMP8m in su(Hw)attP8, UAS-mScarlet in JK16L, w / w; UAS-jGCaMP8m in su(Hw)attP5 / R83H09-AD; UAS-jGCaMP8m in VK00005 / R37G12-DBD* |
| Fig. 5a-d; Fig. S7a-e; Fig. S8a-b | *DNa11>jGCaMP8m, mScarlet* | *UAS-jGCaMP8m in su(Hw)attP8, UAS-mScarlet in JK16L, w / w; UAS-jGCaMP8m in su(Hw)attP5 / R76B02-AD; UAS-jGCaMP8m in VK00005 / VT040347-DBD* |
| Fig. 5a-d; Fig. S7a-e; Fig. S8a-b | *DNae003>jGCaMP8m, mScarlet* | *UAS-jGCaMP8m in su(Hw)attP8, UAS-mScarlet in JK16L, w / w; UAS-jGCaMP8m in su(Hw)attP5 / VT050660-AD; UAS-jGCaMP8m in VK00005 / R22G01-DBD* |
| Fig. 5a-d; Fig. S7a-e; Fig. S8a-b | *DNa03>jGCaMP8m, mScarlet* | *UAS-jGCaMP8m in su(Hw)attP8, UAS-mScarlet in JK16L, w / w; UAS-jGCaMP8m in su(Hw)attP5 / VT033616-AD; UAS-jGCaMP8m in VK00005 / VT025718-DBD* |
| Fig. 5a-d; Fig. S7a-e; Fig. S8a-b | *DNa02>jGCaMP8m, mScarlet* | *UAS-jGCaMP8m in su(Hw)attP8, UAS-mScarlet in JK16L, w / w; UAS-jGCaMP8m in su(Hw)attP5 / R75C10-AD; UAS-jGCaMP8m in VK00005 / R87D07-DBD* |
| Fig. 5e-f; Fig. S9a-c | *LAL013-SS1>6xGFP* | *wDL; R83H09-AD / +; R37G12-DBD / 20XUAS-6XGFP* |
| Fig. 5e-f; Fig. S9a-c | *DNa03-SS1>6xGFP* | *wDL; VT025717-AD / +; VT028555-DBD / 20XUAS-6XGFP* |
| Fig. 5g-l; Fig. S11a-l; Fig. S12a-j; Fig. S13a-c | *DNa03, DNa11 > RSET-jGCaMP8m, mScarlet* | *UAS-jGCaMP8m in su(Hw)attP8, UAS-mScarlet in JK16L, w / UAS-RSET-jGCaMP8m in su(Hw)attP8, w; VT025717-AD, UAS-RSET-jGCaMP8m in su(Hw)attP5 / R76B02-AD, UAS-RSET-jGCaMP8m in su(Hw)attP5; VT028555-DBD, UAS-RSET-jGCaMP8m in VK00005 / VT040347-DBD* |
| Fig. S1a | *LAL013-SS2: SS60262* | *UAS-CsChrimson-mVenus in attP18, w / w; + / R83H09-AD; + / VT002063-DBD* |
| Fig. S1b | *ExR7-SS1: SS85718* | *UAS-CsChrimson-mVenus in attP18, w / w; + / VT049727-AD; + / VT009668-DBD* |
| Fig. S1c; Fig. S4b | *DNa11-SS: SS77821* | *UAS-CsChrimson-mVenus in attP18, w / w; + / R76B02-AD; + / VT040347-DBD* |
| Fig. S1d; Fig. S4c | *DNae003-SS: SS94676* | *UAS-CsChrimson-mVenus in attP18, w / w; + / VT050660-AD; + / R22G01-DBD* |
| Fig. S1e; Fig. S4a | *DNa03-SS1: SS96514* | *UAS-CsChrimson-mVenus in attP18, w / w; + / VT025717-AD; + / VT028555-DBD* |
| Fig. S1f | *DNa03-SS2: SS84990* | *UAS-CsChrimson-mVenus in attP18, w / w; + / VT033616-AD; + / VT025718-DBD* |

Supplementary Table 1

| | | |
|---|---|---|
| Fig. S1g | *LAL121-SS: SS93505* | *UAS-CsChrimson-mVenus in attP18, w / w; + / R65H04-AD; + / VT061910-DBD* |
| Fig. S1h | *ExR7-SS2: SS76739* | *UAS-CsChrimson-mVenus in attP18, w / w; + / R65A07-AD; + / VT049427-DBD* |
| Fig. S14e-h | *DNa06> SPARC2-D-CsChrimson* | *nSyb-PhiC31 in attP18 / w; UAS-SPARC2-D-CsChrimson in attP40 / R54E05-AD; + / VT025718-DBD* |

Supplementary Table 2

**Hemibrain v1.2.1 & FlyWire v783 & MANC v1.2.1**

Upstream of LAL013 (Fig. 1t)

| | hemibrain_id_post | | type_post | | | | nt_post | FlyWire_LAL013_R | | FlyWire_LAL013_L | |
|---|---|---|---|---|---|---|---|---|---|---|---|
| | 894702888 | | LAL013 | | | | ACH | 720575940630675151 | | 720575940612424362 | |
| rank | hemibrain_id_pre | weight | type_pre | hemibrain_type | FlyWire_type | other_type | nt_pre | FlyWire_id_pre | R_weight | FlyWire_id_pre | L_weight |
| 1 | 667241165 | 96 | SMP192 | SMP192 | | | ACH | 720575940621526911 | 62 | 720575940613859379 | 84 |
| 2 | 1105882358 | 86 | ExR7 | ExR7 | | | ACH | 720575940612937073 | 62 | 720575940650785401 | 15 |
| 3 | 5812987684 | 81 | ExR7 | ExR7 | | | ACH | 720575940650785401 | 33 | 720575940612937073 | 65 |
| 4 | 1259386264 | 76 | ExR7 | ExR7 | | | ACH | 720575940631626491 | 54 | 720575940645057828 | 51 |
| 5 | 544637659 | 69 | SMP192 | SMP192 | | | ACH | 720575940613859379 | 45 | 720575940621526911 | 47 |
| 6 | 700002191 | 58 | mALB5 | mALB5 | | | GABA | 720575940645989655 | 45 | 720575940629728811 | 91 |
| 7 | 2003459912 | 56 | CB0556 | | CB0556 | | GABA | 720575940626138701 | 10 | 720575940631283512 | 34 |
| 8 | 1353375816 | 54 | AN_multi_11 | | AN_multi_11 | | GABA | 720575940629486714 | 33 | 720575940610640482 | 63 |
| 9 | 1005710800 | 51 | ExR7 | ExR7 | | | ACH | 720575940645057828 | 47 | 720575940631626491 | 30 |
| 10 | 1344858721 | 50 | LAL139 | LAL139 | | | GABA | 720575940640053181 | 31 | 720575940631043139 | 64 |
| 11 | 1687321237 | 43 | AN_multi_11 | | AN_multi_11 | | GABA | 720575940610640482 | 45 | 720575940629486714 | 33 |
| 12 | 1882040130 | 42 | CB0149 | | CB0149 | | GLUT | 720575940622140109 | 15 | 720575940618722542 | 13 |
| 13 | 1529073617 | 37 | PS196a | PS196 | PS196a | | ACH | 720575940629229020 | 40 | 720575940625082969 | 35 |
| 14 | 1292367728 | 33 | mALD4 | mALD4 | | | GABA | 720575940639542691 | 24 | 720575940631365260 | 20 |
| 15 | 1848277193 | 25 | CB0625 | | CB0625 | | GABA | 720575940621174337 | 17 | 720575940639278781 | 24 |
| 16 | 1934975261 | 23 | CB0663 | | CB0663 | | GLUT | 720575940631812141 | 12 | 720575940615032130 | 18 |
| 17 | 1201296549 | 20 | LAL190 | LAL190 | | | ACH | 720575940614619282 | 19 | 720575940632469964 | 15 |
| 18 | 1634140884 | 19 | LAL113 | LAL113 | | | GABA | 720575940610075346 | 17 | 720575940625048250 | 9 |
| 19 | 1635833170 | 19 | LAL113 | LAL113 | | | GABA | 720575940640608219 | 12 | 720575940620431798 | 14 |
| 20 | 1967637295 | 17 | LAL124 | | LAL124 | | GLUT | 720575940622281904 | 40 | 720575940621768436 | 24 |

Downstream of LAL013 (Fig. 1t)

| | hemibrain_id_pre | | type_pre | | | | nt_pre | FlyWire_LAL013_R | | FlyWire_LAL013_L | |
|---|---|---|---|---|---|---|---|---|---|---|---|
| | 894702888 | | LAL013 | | | | ACH | 720575940630675151 | | 720575940612424362 | |
| rank | hemibrain_id_post | weight | type_post | hemibrain_type | FlyWire_type | other_type | nt_post | FlyWire_id_post | R_weight | FlyWire_id_post | L_weight |
| 1 | 5813069484 | 175 | **DNae014** | PS017 | **DNa15** | **DNae014*** | ACH | 720575940631759663 | 129 | 720575940609488942 | 150 |
| 2 | 5813057245 | 173 | DNa11 | VES008 | DNa11 | | ACH | 720575940611735514 | 102 | 720575940623019544 | 100 |
| 3 | 1170939344 | 92 | **DNae003** | DNa01 | **DNae001** | **DNae003*** | ACH | 720575940618167579 | 58 | 720575940626328707 | 77 |
| 4 | 5813069496 | 89 | LAL074 | LAL084 | LAL074 | | GLUT | 720575940610211779 | 73 | 720575940619866027 | 86 |
| 5 | 2003459912 | 88 | CB0556 | | CB0556 | | GABA | 720575940626138701 | 77 | 720575940631283512 | 78 |
| 6 | 5813057723 | 85 | PS019 | PS019 | | | ACH | 720575940625561406 | 63 | 720575940629786624 | 65 |
| 7 | 1139909038 | 85 | DNa03 | DNa03 | | | ACH | 720575940620918789 | 48 | 720575940630085583 | 59 |
| 8 | 1812488920 | 72 | PS019 | PS019 | | | ACH | 720575940640205501 | 39 | 720575940625477322 | 49 |
| 9 | 1383111671 | 71 | PS274 | PS274 | | | ACH | 720575940624323110 | 61 | 720575940623992317 | 72 |
| 10 | 1201296549 | 68 | LAL190 | LAL190 | | | ACH | 720575940614619282 | 102 | 720575940632469964 | 122 |
| 11 | 1572390197 | 57 | LAL074 | LAL074 | | | GLUT | 720575940616807897 | 57 | 720575940617226141 | 80 |
| 12 | 1622540849 | 46 | CB0901 | | CB0901 | Ruby | ACH | 720575940622361270 | 9 | 720575940622168052 | 21 |
| 13 | 1629383757 | 39 | CB0584 | | CB0584 | | GABA | 720575940632042148 | 45 | 720575940633901395 | 45 |
| 14 | 5813047453 | 37 | LAL001 | LAL001 | | | GLUT | 720575940628453690 | 13 | 720575940635937678 | 32 |
| 15 | 1448575109 | 30 | LNO2 | LNO2 | | | GABA | 720575940622945037 | 35 | 720575940615620002 | 9 |
| 16 | 1140245595 | 29 | DNa02 | DNa02 | | | ACH | 720575940629327659 | 51 | 720575940604737708 | 41 |
| 17 | 1140249804 | 28 | LAL018 | LAL018 | | | ACH | 720575940611385806 | 9 | 720575940620686964 | 21 |
| 18 | 1874217622 | 27 | DNg75 | | DNg75 | | ACH | 720575940614730914 | 14 | 720575940612603289 | 25 |
| 19 | 1477239390 | 24 | DNb02 | DNb02 | | | GLUT | 720575940616936226 | 14 | 720575940622540899 | 17 |
| 20 | 1534270924 | 24 | VES041 | VES041 | | | GABA | 720575940627253604 | 57 | 720575940635202204 | 67 |

***FlyWire legacy names in early 2024; DNae014 adopted in Ros et al. (2024) and DNae003 in Feng et al. (2024); "DNae003" is now used for another neuron in FlyWire**

DNa03-LAL013 hierarchy and central complex inputs (Fig. 2b)

| type_pre | nt | weight | type_post |
|---|---|---|---|
| LAL013 | ACT | 173 | DNa11 |
| LAL013 | ACT | 85 | DNa03 |
| LAL013 | ACT | 29 | DNa02 |
| LAL013 | ACT | 92 | DNae003 |
| LAL013 | ACT | 175 | DNae014 |

Supplementary Table 2

| | | | | | | |
|---|---|---|---|---|---|---|
| DNa11 | ACT | 21 | DNa03 | | | |
| DNa11 | ACT | 185 | DNa02 | | | |
| DNa11 | ACT | 30 | DNae003 | | | |
| DNa11 | ACT | 19 | DNae014 | | | |
| DNa03 | ACT | 16 | LAL013 | | | |
| DNa03 | ACT | 200 | DNa11 | | | |
| DNa03 | ACT | 271 | DNa02 | | | |
| DNa03 | ACT | 243 | DNae014 | | | |
| PFL3_L | ACT | 239 | DNa03 | | | |
| PFL3_L | ACT | 275 | DNa02 | | | |
| PFL3_R | ACT | 1805 | LAL121_L | | | |
| PFL3_R | ACT | 837 | AOTU019_L | | | |
| PFL3_R | ACT | 855 | LAL040_L | | | |
| ExR7 | ACT | 294 | LAL013 | | | |
| PFL2 | ACT | 808 | DNa03 | | | |
| LAL121_L | VGLUT | 382 | DNa03 | | | |
| AOTU019_L | GABA | 65 | DNa11 | | | |
| AOTU019_L | GABA | 352 | DNa03 | | | |
| AOTU019_L | GABA | 192 | DNa02 | | | |
| AOTU019_L | GABA | 19 | DNae003 | | | |
| AOTU019_L | GABA | 362 | DNae014 | | | |
| LAL040_L | GABA | 60 | DNa11 | | | |
| LAL040_L | GABA | 108 | DNa03 | | | |
| LAL040_L | GABA | 139 | DNa02 | | | |
| LAL040_L | GABA | 84 | DNae003 | | | |
| type | hemibrain_id | | PFL3_R | ExR7_L&R | PFL2_L&R | PFL3_L |
| LAL013_R | 894702888 | | 787374226 | 1105882358 | 481112669 | 1004700437 |
| DNa03_R | 1139909038 | | 910447181 | 5812987684 | 635234035 | 941939879 |
| DNa11_R | 5813057245 | | 757694775 | 1259386264 | 760094565 | 666994301 |
| DNae014_R | 5813069484 | | 911134017 | 1005710800 | 850194651 | 1258686925 |
| DNa02_R | 1140245595 | | 1097718659 | | 880875840 | 789130596 |
| DNae003_R | 1170939344 | | 941132430 | | 882239837 | 912925080 |
| LAL121_L | 1510311956 | | 1008028537 | | 916284630 | 850855138 |
| AOTU019_L | 1605518663 | | 1258073453 | | 948333783 | 666308023 |
| LAL040_L | 796978752 | | 942172835 | | 979752365 | 911569552 |
| PFL3_R | see right | | 912488890 | | 1167282562 | 851493896 |
| ExR7_L&R | see right | | 1134253374 | | 1200053009 | 944262351 |
| PFL2_L&R | see right | | 1200032115 | | 1381484065 | 880875927 |
| PFL3_L | see right | | | | | |
| Upstream of DNa03 in the brain (Fig. 3a) | | | | | | |
| | Pre_type | | Post_type | | | |
| | PFL3 | | LAL121_Right | LAL121_Left | DNa03_Right | DNa03_Left |
| | FlyWire_id | | weight | weight | weight | weight |
| | 720575940616654370 | | 0 | 183 | 0 | 21 |
| | 720575940619802800 | | 0 | 126 | 0 | 11 |
| | 720575940635430030 | | 2 | 191 | 0 | 29 |
| | 720575940618759361 | | 167 | 0 | 18 | 0 |
| | 720575940623430621 | | 185 | 0 | 22 | 0 |
| | 720575940612870952 | | 0 | 225 | 0 | 29 |
| | 720575940625308162 | | 1 | 216 | 0 | 25 |
| | 720575940628587202 | | 183 | 0 | 19 | 0 |
| | 720575940635481262 | | 0 | 154 | 0 | 13 |
| | 720575940631151663 | | 1 | 220 | 0 | 35 |
| | 720575940644706083 | | 0 | 205 | 0 | 26 |
| | 720575940612936154 | | 166 | 0 | 20 | 0 |
| | 720575940630653604 | | 182 | 0 | 22 | 0 |
| | 720575940605640905 | | 0 | 128 | 0 | 11 |
| | 720575940638901875 | | 0 | 126 | 0 | 19 |
| | 720575940619648116 | | 178 | 1 | 43 | 0 |

Supplementary Table 2

| | | | | | | |
|---|---|---|---|---|---|---|
| | 720575940620202266 | | 149 | 1 | 13 | 0 |
| | 720575940622234326 | | 201 | 1 | 15 | 0 |
| | 720575940636868186 | | 0 | 119 | 0 | 25 |
| | 720575940619006087 | | 0 | 198 | 0 | 33 |
| | 720575940624320318 | | 160 | 1 | 21 | 0 |
| | 720575940628457448 | | 122 | 0 | 16 | 0 |
| | 720575940630728588 | | 134 | 0 | 18 | 0 |
| | 720575940622727629 | | 207 | 0 | 22 | 0 |
| | Sum by side | | 2034 | 2091 | 249 | 277 |
| | | | | | DNa03_Right | DNa03_Left |
| | LAL121 | | | | 720575940630085583 | 720575940620918789 |
| | 720575940617374242 | | | | 1 | 467 |
| | 720575940623689959 | | | | 456 | 0 |
| EPG to ExR7 connectivity (Fig. 4a for illustration of the all-to-all connectivity pattern) | | | | | | |
| | Pre_type | | Post_type | | | |
| | EPG | | ExR7 | | | |
| | hemibrain_id | | 1005710800 | 1105882358 | 1259386264 | 5812987684 |
| | 387364605 | | 11 | 14 | 21 | 15 |
| | 416642425 | | 15 | 13 | 9 | 11 |
| | 449438847 | | 16 | 16 | 14 | 20 |
| | 478375456 | | 15 | 19 | 16 | 18 |
| | 541118908 | | 20 | 10 | 20 | 22 |
| | 541870397 | | 15 | 19 | 9 | 13 |
| | 572870540 | | 16 | 21 | 18 | 20 |
| | 632544268 | | 12 | 18 | 12 | 10 |
| | 633546217 | | 17 | 17 | 14 | 10 |
| | 634962055 | | 6 | 10 | 5 | 6 |
| | 665314820 | | 1 | 16 | 8 | 7 |
| | 694920753 | | 5 | 11 | 13 | 8 |
| | 695629525 | | 9 | 15 | 8 | 8 |
| | 695956656 | | 8 | 10 | 10 | 7 |
| | 696362840 | | 5 | 6 | 6 | 6 |
| | 697001770 | | 10 | 14 | 14 | 11 |
| | 725951521 | | 7 | 8 | 5 | 7 |
| | 758419409 | | 13 | 21 | 26 | 17 |
| | 788794171 | | 10 | 19 | 11 | 8 |
| | 789126240 | | 3 | 9 | 6 | 14 |
| | 819828986 | | 8 | 19 | 10 | 15 |
| | 910438331 | | 7 | 16 | 6 | 11 |
| | 912545106 | | 8 | 17 | 6 | 13 |
| | 912601268 | | 9 | 12 | 9 | 12 |
| | 941132434 | | 4 | 12 | 8 | 10 |
| | 942491983 | | 9 | 11 | 2 | 11 |
| | 1002507159 | | 17 | 13 | 14 | 12 |
| | 1002852791 | | 11 | 13 | 11 | 2 |
| | 1004017998 | | 13 | 16 | 14 | 16 |
| | 1034219901 | | 8 | 19 | 19 | 25 |
| | 1035045015 | | 15 | 15 | 11 | 17 |
| | 1065410141 | | 9 | 13 | 10 | 10 |
| | 1125964814 | | 6 | 17 | 7 | 10 |
| | 1126647624 | | 8 | 13 | 7 | 4 |
| | 1127476096 | | 9 | 18 | 5 | 5 |
| | 1167995070 | | 16 | 19 | 12 | 18 |
| | 1168664495 | | 26 | 20 | 11 | 17 |
| | 1447576662 | | 19 | 15 | 7 | 19 |
| | 5813012006 | | 22 | 16 | 17 | 24 |
| | 5813014873 | | 20 | 35 | 22 | 33 |
| | 5813022281 | | 8 | 15 | 5 | 14 |
| | 5813027103 | | 5 | 14 | 9 | 15 |

| | | | | | | | | | |
|---|---|---|---|---|---|---|---|---|---|
| | 5813040233 | | 9 | 18 | 12 | 17 | | | |
| | 5813061251 | | 6 | 6 | 11 | 15 | | | |
| | 5813077544 | | 14 | 16 | 13 | 18 | | | |
| | 5813080838 | | 8 | 17 | 16 | 9 | | | |
| | ExR7 | | | | | | | | |
| | 1005710800 | | 0 | 74 | 39 | 45 | | | |
| | 1105882358 | | 40 | 0 | 46 | 49 | | | |
| | 1259386264 | | 27 | 79 | 0 | 44 | | | |
| | 5812987684 | | 52 | 75 | 58 | 0 | | | |
| Upstream of DNa03 in the brain (Fig. S10) | | | | | | | | | |
| | | | Post_type | | | | | | |
| | | | DNa03_Right | | DNa03_Left | | | | |
| Pre_type | FlyWire_id | | 720575940630085583 | | 720575940620918789 | | nt_pre | connection_side | mean_weight |
| PFL2 | sum of 12, see below | | 726 | | 809 | | ACH | Bilateral | 767.5 |
| LAL014 | 720575940623632904 | | 456 | | 520 | | ACH | Ipsi | 488 |
| LAL121 | 720575940617374242 | | 456 | | 467 | | GLUT | Contra | 461.5 |
| LAL112 | sum of 2, see below | | 410 | | 488 | | GABA | Ipsi | 449 |
| LAL122 | 720575940638677411 | | 357 | | 436 | | GLUT | Contra | 396.5 |
| VES041 | sum of 2, see below | | 345 | | 366 | | GABA | Bilateral | 355.5 |
| LAL051 | 720575940623067389 | | 319 | | 366 | | GLUT | Ipsi | 342.5 |
| LAL171 | sum of 2, see below | | 274 | | 376 | | ACH | Contra | 325 |
| AOTU019 | 720575940631517251 | | 307 | | 305 | | GABA | Contra | 306 |
| PVLP138 | 720575940641706184 | | 247 | | 308 | | ACH | Contra | 277.5 |
| PFL3 | sum of 12, see above | | 249 | | 277 | | ACH | Contra | 263 |
| AOTUv1A_T01 | 720575940608339209 | | 214 | | 250 | | GABA | Contra | 232 |
| LC33 | 720575940633801261 | | 190 | | 259 | | GLUT | Ipsi | 224.5 |
| LAL113 | sum of 2, see below | | 220 | | 219 | | GABA | Ipsi | 219.5 |
| PS183 | 720575940615359794 | | 203 | | 231 | | ACH | Ipsi | 217 |
| CB0132 | 720575940625172750 | | 180 | | 189 | | ACH | Ipsi | 184.5 |
| AOTUv3B_P01 | 720575940633268575 | | 186 | | 176 | | ACH | Ipsi | 181 |
| LT51 | sum of 4, see below | | 188 | | 164 | | GLUT | Ipsi | 176 |
| LAL015 | 720575940615755147 | | 145 | | 186 | | ACH | Ipsi | 165.5 |
| PLP228 | 720575940629768722 | | 167 | | 160 | | ACH | Contra | 163.5 |
| PS100 | 720575940633308371 | | 172 | | 153 | | GABA | Ipsi | 162.5 |
| PS049 | 720575940623533749 | | 147 | | 172 | | GABA | Ipsi | 159.5 |
| MBON31 | 720575940644615716 | | 157 | | 158 | | GABA | Bilateral | 157.5 |
| PFL2 | 720575940620657140 | | 37 | | 56 | | | | |
| | 720575940635467150 | | 52 | | 88 | | | | |
| | 720575940635990640 | | 67 | | 75 | | | | |
| | 720575940624754981 | | 71 | | 46 | | | | |
| | 720575940629922934 | | 79 | | 72 | | | | |
| | 720575940620952865 | | 83 | | 73 | | | | |
| | 720575940623585255 | | 63 | | 69 | | | | |
| | 720575940623937976 | | 47 | | 49 | | | | |
| | 720575940638678616 | | 75 | | 67 | | | | |
| | 720575940635153527 | | 57 | | 88 | | | | |
| | 720575940617498688 | | 52 | | 59 | | | | |
| | 720575940626763688 | | 43 | | 67 | | | | |
| LAL112 | 720575940629393207 | | | | 271 | | | | |
| | 720575940629377132 | | | | 217 | | | | |
| | 720575940623426013 | | 211 | | | | | | |
| | 720575940643551392 | | 199 | | | | | | |
| VES041 | 720575940635202204 | | 186 | | 216 | | | | |
| | 720575940627253604 | | 159 | | 150 | | | | |
| LAL171 | 720575940624891941 | | | | 192 | | | | |
| | 720575940659019649 | | | | 184 | | | | |
| | 720575940615651670 | | 151 | | | | | | |
| | 720575940645716771 | | 123 | | | | | | |
| AOTUv1A_T0 | 720575940608339209 | | | | 126 | | | | |

Supplementary Table 2

| | | | | | | | | | | | |
|---|---|---|---|---|---|---|---|---|---|---|---|
| | 720575940628842240 | | | | 124 | | | | | | |
| | 720575940614386942 | | 124 | | | | | | | | |
| | 720575940620283307 | | 90 | | | | | | | | |
| LAL113 | 720575940610075346 | | | | 111 | | | | | | |
| | 720575940640608219 | | | | 108 | | | | | | |
| | 720575940620431798 | | 128 | | | | | | | | |
| | 720575940625048250 | | 92 | | | | | | | | |
| MBON31 | 720575940644615716 | | 64 | | 109 | | | | | | |
| | 720575940636992368 | | 93 | | 49 | | | | | | |
| Downstream DNs of DNa03 in the brain (Fig. S10) | | | | | | | | | | | |
| Left | DNa03: 720575940620918789 | | | | | | Right | DNa03: 720575940630085583 | | | |
| Pre_type | FlyWire_weight | Post_type | FlyWire_ID | | | | Pre_type | FlyWire_weight | Post_type | FlyWire_ID | |
| DNa03 | 315 | **DNa16** | 720575940632962786 | | | | DNa03 | 365 | **DNa16** | 720575940621519130 | |
| | 309 | **DNa02** | 720575940629327659 | | | | | 301 | **DNa02** | 720575940604737708 | |
| | 290 | **DNa15** | 720575940631759663 | | | | | 290 | **DNa15** | 720575940609488942 | |
| | 211 | **DNa11** | 720575940611735514 | | | | | 256 | **DNa11** | 720575940623019544 | |
| | 174 + 159 | **DNa13** | 720575940616286175 | | 720575940613408552 | | | 170 + 129 | **DNa13** | 720575940637607646 | 720575940625502666 |
| | 145 + 138 | **DNg04** | 720575940624082931 | | 720575940643817888 | | | 224 + 64 | **DNg04** | 720575940621325034 | 720575940624968013 |
| | 130 | **DNa06** | 720575940624986919 | | | | | 119 | **DNa06** | 720575940640325429 | |
| | 75 | **DNpe023** | 720575940616604817 | | | | | 74 | **DNpe023** | 720575940633011256 | |
| | | | 720575940624816115 | | | | | | | | |
| Pre_type | FlyWire_weight | 1st_type | FlyWire_weight | 2nd_type | | | Pre_type | FlyWire_weight | 1st_type | FlyWire_weight | 2nd_type |
| DNa03 | 315 | **DNa16** | 57 | **DNge033** | | | DNa03 | 365 | **DNa16** | 81 | **DNge033** |
| | | **DNa16** | 25 | **DNg89** | | | | | **DNa16** | 53 | **DNg89** |
| DNa03 | 309 | **DNa02** | 62 | **DNa06** | | | DNa03 | 301 | **DNa02** | 106 | **DNa06** |
| | | **DNa02** | 61 | **DNa16** | | | | | **DNa02** | 41 | **DNge033** |
| | | **DNa02** | 53 | **DNge033** | | | | | **DNa02** | 79 | **DNge026** |
| | | **DNa02** | 39 | **DNge026** | | | | | **DNa02** | 73 | **DNa16** |
| DNa03 | 211 | **DNa11** | 190 | **DNa02** | | | DNa03 | 256 | **DNa11** | 200 | **DNa02** |
| | | **DNa11** | 186 | **DNg75** | | | | | **DNa11** | 202 | **DNg75** |
| | | **DNa11** | 120 | **DNa06** | | | | | **DNa11** | 193 | **DNa06** |
| | | **DNa11** | 108 | **DNge050** | | | | | **DNa11** | 55 | **DNge050** |
| | | **DNa11** | 52 | **DNge050** | | | | | **DNa11** | 76 | **DNge050** |
| DNa03 | 185 | PLP060 | 194 | **DNp03** | | | DNa03 | 211 | PLP060 | 19 | **DNp03** |
| | | PLP060 | 113 | **DNp05** | | | | | PLP060 | 75 | **DNp05** |
| | | PLP060 | 67 | **DNb05** | | | | | PLP060 | 57 | **DNb05** |
| | | PLP060 | 59 | **DNp11** | | | | | PLP060 | 53 | **DNp11** |
| | | PLP060 | 52 | **DNpe037** | | | | | PLP060 | 62 | **DNpe037** |
| DNa03 | 333 | **DNa13** | 109 | **DNg88** | | | DNa03 | 299 | **DNa13** | 107 | **DNg88** |
| DNa03 | 174 | PS274 | 216 | **DNae002** | | | DNa03 | 164 | PS274 | 163 | **DNae002** |
| | | PS274 | 154 | **DNa04** | | | | | PS274 | 153 | **DNa04** |
| | | PS274 | 72 | **DNa15** | | | | | PS274 | 72 | **DNa15** |
| | | PS274 | 51 | **DNa05** | | | | | PS274 | 77 | **DNa05** |
| | | PS274 | 47 | **DNa09** | | | | | PS274 | 68 | **DNa09** |
| | | PS274 | 44 | **DNb01** | | | | | PS274 | 59 | **DNb01** |
| DNa03 | 169 | LAL073 | 255 | **DNg13** | | | DNa03 | 34 | LAL073 | 262 | **DNg13** |
| | | LAL073 | 191 | **DNbe003** | | | | | LAL073 | 176 | **DNbe003** |
| | | LAL073 | 132 | **DNbe006** | | | | | LAL073 | 156 | **DNbe006** |
| | | LAL073 | 123 | **DNg97** | | | | | LAL073 | 116 | **DNg97** |
| | | LAL073 | 82 | **DNb08_b** | | | | | LAL073 | 35 | **DNb08_b** |
| | | LAL073 | 58 | **DNb08_a** | | | | | LAL073 | 32 | **DNb08_a** |
| | | LAL073 | 64 | **DNb01** | | | | | LAL073 | 84 | **DNb01** |
| DNa03 | 151 | VES041 | 159 | **DNa03** | | | DNa03 | 163 | VES041 | 216 | **DNa03** |
| | | VES041 | 150 | **DNa03** | | | | | VES041 | 186 | **DNa03** |
| DNa03 | 282 | LAL074 | 326 | **DNae002** | | | DNa03 | 343 | LAL074 | 142 | **DNae002** |
| | | LAL074 | 147 | **DNb01** | | | | | LAL074 | 149 | **DNb01** |
| | | LAL074 | 131 | **DNg75** | | | | | LAL074 | 139 | **DNg75** |
| | | LAL074 | 61 | **DNg13** | | | | | LAL074 | 110 | **DNg13** |
| DNa03 | 283 | **DNg04** | 154 | **DNa16** | | | DNa03 | 224 | **DNg04** | 192 | **DNa16** |
| | | **DNg04** | 112 | **DNg75** | | | | | **DNg04** | 113 | **DNg75** |

Supplementary Table 2

| | | | | | | | | | | | |
|---|---|---|---|---|---|---|---|---|---|---|---|
| DNa03 | 130 | **DNa06** | 46 | **DNge006** | | | DNa03 | 119 | **DNa06** | 279 | **DNge006** |
| | | **DNa06** | 173 | **DNge033** | | | | | **DNa06** | 181 | **DNge033** |
| | | **DNa06** | 95 | **DNg75** | | | | | **DNa06** | 89 | **DNg75** |
| | | **DNa06** | 55 | **DNpe013** | | | | | **DNa06** | 58 | **DNpe013** |
| | | **DNa06** | 53 | **DNg89** | | | | | **DNa06** | 101 | **DNg89** |
| | | **DNa06** | 51 | **DNa16** | | | | | **DNa06** | 54 | **DNa16** |
| | | **DNa06** | 46 | **DNa02** | | | | | **DNa06** | 70 | **DNa02** |
| | | **DNa06** | 29 | **DNge052** | | | | | **DNa06** | 67 | **DNge052** |
| | | **DNa06** | 45 | **DNge072** | | | | | **DNa06** | 57 | **DNge072** |
| DNa03 | 115 | PS100 | 323 | **DNa16** | | | DNa03 | 120 | PS100 | 351 | **DNa16** |
| | | PS100 | 192 | **DNa15** | | | | | PS100 | 208 | **DNa15** |
| | | PS100 | 165 | **DNa04** | | | | | PS100 | 186 | **DNa04** |
| | | PS100 | 164 | **DNg75** | | | | | PS100 | 178 | **DNg75** |
| | | PS100 | 157 | **DNge026** | | | | | PS100 | 204 | **DNge026** |
| | | PS100 | 153 | **DNa03** | | | | | PS100 | 172 | **DNa03** |
| | | PS100 | 301 | **DNg04** | | | | | PS100 | 305 | **DNg04** |
| | | PS100 | 109 | **DNae003** | | | | | PS100 | 89 | **DNae003** |
| | | PS100 | 68 | **DNa05** | | | | | PS100 | 38 | **DNa05** |
| | | PS100 | 68 | **DNg88** | | | | | PS100 | 92 | **DNg88** |
| | | PS100 | 67 | **DNge042** | | | | | PS100 | 60 | **DNge042** |
| | | PS100 | 61 | **DNa06** | | | | | PS100 | 72 | **DNa06** |
| | | PS100 | 51 | **DNp15** | | | | | PS100 | 104 | **DNp15** |
| | | PS100 | 41 | **DNge123** | | | | | PS100 | 64 | **DNge123** |
| | | PS100 | 37 | **DNge125** | | | | | PS100 | 50 | **DNge125** |
| DNa03 | 95 | LAL018 | 249 | **DNa15** | | | DNa03 | 100 | LAL018 | 248 | **DNa15** |
| | | LAL018 | 241 | **DNa02** | | | | | LAL018 | 221 | **DNa02** |
| | | LAL018 | 229 | **DNae002** | | | | | LAL018 | 265 | **DNae002** |
| | | LAL018 | 147 | **DNa04** | | | | | LAL018 | 181 | **DNa04** |
| | | LAL018 | 144 | **DNa16** | | | | | LAL018 | 160 | **DNa16** |
| | | LAL018 | 98 | **DNg75** | | | | | LAL018 | 122 | **DNg75** |
| | | LAL018 | 87 | **DNa01** | | | | | LAL018 | 80 | **DNa01** |
| | | LAL018 | 69 | **DNg97** | | | | | LAL018 | 55 | **DNg97** |
| | | LAL018 | 97 | **DNg04** | | | | | LAL018 | 130 | **DNg04** |
| DNa03 | 95 | PS090a | 272 | **DNae010** | | | DNa03 | 106 | PS090a | 253 | **DNae010** |
| | | PS090a | 88 | **DNp63** | | | | | PS090a | 118 | **DNp63** |
| | | PS090a | 44 | **DNp63** | | | | | PS090a | 79 | **DNp63** |
| DNa03 | 82 | PS018a | 187 | **DNa16** | | | DNa03 | 108 | PS018a | 198 | **DNa16** |
| | | PS018a | 172 | **DNa15** | | | | | PS018a | 233 | **DNa15** |
| | | PS018a | 242 | **DNg04** | | | | | PS018a | 276 | **DNg04** |
| | | PS018a | 111 | **DNa04** | | | | | PS018a | 113 | **DNa04** |
| | | PS018a | 85 | **DNa02** | | | | | PS018a | 114 | **DNa02** |
| | | PS018a | 57 | **DNa06** | | | | | PS018a | 47 | **DNa06** |
| DNa03 | 75 | **DNpe023** | 193 | **DNp50** | | | DNa03 | 74 | **DNpe023** | 153 | **DNp50** |
| | | **DNpe023** | 84 | **DNg88** | | | | | **DNpe023** | 48 | **DNg88** |
| | | **DNpe023** | 39 | **DNg97** | | | | | **DNpe023** | 50 | **DNg97** |
| | | **DNpe023** | 82 | **DNae005** | | | | | **DNpe023** | 82 | **DNae005** |
| | | **DNpe023** | 140 | **DNp50** | | | | | **DNpe023** | 131 | **DNp50** |
| DNa03 | 68 | CB0556 | 137 | **DNa16** | | | DNa03 | 69 | CB0556 | 152 | **DNa16** |
| | | CB0556 | 115 | **DNa03** | | | | | CB0556 | 110 | **DNa03** |
| | | CB0556 | 120 | **DNa13** | | | | | CB0556 | 120 | **DNa13** |
| | | CB0556 | 56 | **DNae005** | | | | | CB0556 | 67 | **DNae005** |
| | | CB0556 | 103 | **DNg04** | | | | | CB0556 | 103 | **DNg04** |
| | | CB0556 | 50 | **DNa11** | | | | | CB0556 | 33 | **DNa11** |
| DNa03 | 68 | cL22b | 391 | **DNpe016** | | | DNa03 | 59 | cL22b | 336 | **DNpe016** |
| | | cL22b | 55 | **DNp102** | | | | | cL22b | 49 | **DNp102** |
| | | cL22b | 44 | **DNp05** | | | | | cL22b | 53 | **DNp05** |
| DNa03 | 100 | PS137 | 244 | **DNa06** | | | DNa03 | 98 | PS137 | 302 | **DNa06** |
| | | PS137 | 176 | **DNae003** | | | | | PS137 | 156 | **DNae003** |
| | | PS137 | 181 | **DNa16** | | | | | PS137 | 243 | **DNa16** |
| | | PS137 | 288 | **DNb02** | | | | | PS137 | 327 | **DNb02** |
| | | PS137 | 120 | **DNae002** | | | | | PS137 | 142 | **DNae002** |

Supplementary Table 2

| | | | | | | | | | | | |
|---|---|---|---|---|---|---|---|---|---|---|---|
| | | PS137 | 129 | **DNg49** | | | | | PS137 | 14 | **DNg49** |
| | | PS137 | 96 | **DNa09** | | | | | PS137 | 116 | **DNa09** |
| | | PS137 | 62 | **DNb01** | | | | | PS137 | 113 | **DNb01** |
| DNa03 | 54 | PS065 | 160 | **DNpe022** | | | DNa03 | 82 | PS065 | 204 | **DNpe022** |
| | | PS065 | 125 | **DNp57** | | | | | PS065 | 108 | **DNp57** |
| | | PS065 | 115 | **DNpe003** | | | | | PS065 | 110 | **DNpe003** |
| | | PS065 | 46 | **DNp05** | | | | | PS065 | 87 | **DNp05** |
| DNa03 | 40 | PS080 | 138 | **DNae003** | | | DNa03 | 67 | PS080 | 127 | **DNae003** |
| | | PS080 | 98 | **DNge107** | | | | | PS080 | 105 | **DNge107** |
| | | PS080 | 85 | **DNp102** | | | | | PS080 | 79 | **DNp102** |
| | | PS080 | 111 | **DNa09** | | | | | PS080 | 78 | **DNa09** |
| | | PS080 | 74 | **DNb01** | | | | | PS080 | 55 | **DNb01** |
| | | PS080 | 85 | **DNg02_c** | | | | | PS080 | 53 | **DNg02_c** |
| | | PS080 | 60 | **DNg02_e** | | | | | PS080 | 37 | **DNg02_e** |
| DNa03 | 64 | CB0164 | 111 | **DNa16** | | | DNa03 | 69 | CB0164 | 121 | **DNa16** |
| | | CB0164 | 94 | **DNb02** | | | | | CB0164 | 108 | **DNb02** |
| | | CB0164 | 67 | **DNa15** | | | | | CB0164 | 53 | **DNa15** |
| | | CB0164 | 62 | **DNa06** | | | | | CB0164 | 49 | **DNa06** |
| DNa03 | 105 | LAL119 | 20 | **DNg63** | | | DNa03 | 86 | LAL119 | 55 | **DNg63** |
| DNa03 | 34 | CB1766 | 189 | **DNp31** | | | DNa03 | 67 | CB1766 | 183 | **DNp31** |
| Downstream of DNa11 in the brain (Fig. 6I) | | | | | | | | | | | |
| | hemibrain_id_pre | | type_pre | | | | nt_pre | FlyWire_DNa11_R | | FlyWire_DNa11_L | |
| | 5813057245 | | DNa11 | | | | ACH | 720575940611735514 | | 720575940623019544 | |
| rank | hemibrain_id_pre | weight | type_pre | hemibrain_type | FlyWire_type | | nt_pre | FlyWire_id_pre | R_weight | FlyWire_id_pre | L_weight |
| 1 | 1140245595 | 185 | DNa02 | DNa02 | | | ACH | 720575940629327659 | 149 | 720575940604737708 | 185 |
| 2 | 1963523237 | 123 | DNa06 | PS039 | DNa06 | | ACH | 720575940624986919 | 76 | 720575940640325429 | 132 |
| 3 | 1874217622 | 114 | DNg75 | | DNg75 | | ACH | 720575940614730914 | 120 | 720575940612603289 | 127 |
| Downstream of DNa11 in the VNC (Fig. 6I) | | | | | | | | | | | |
| | weight = average (left, right) | | | | PMN->MN weight = max (left, right) | | | | | | |
| type_pre | weight | post_id | type_post | PMN neuropil | PMN->MN weight | MN type | | MN neuropil | | | |
| DNa11 | 84.5 | IN03A010 | ACH | T1_ipsi | 401 | Fe reductor MN | | T1_ipsi | | | |
| DNa11 | 84.5 | IN03A010 | ACH | T1_ipsi | 76 | Pleural remotor/abductor MN | | T1_ipsi | | | |
| DNa11 | 54.5 | IN03A010 | ACH | T2_ipsi | 390 | Sternal posterior rotator MN | | T2_ipsi | | | |
| DNa11 | 54.5 | IN03A010 | ACH | T2_ipsi | 303 | Pleural remotor/abductor MN | | T2_ipsi | | | |
| DNa11 | 31.5 | IN19A003 | GABA | T2_ipsi | 1284 | Sternal posterior rotator MN | | T2_ipsi | | | |
| DNa11 | 31.5 | IN19A003 | GABA | T2_ipsi | 219 | Pleural remotor/abductor MN | | T2_ipsi | | | |
| DNa11 | 45.5 | IN03A010 | ACH | T3_ipsi | 518 | Sternal posterior rotator MN | | T3_ipsi | | | |
| DNa11 | 45.5 | IN03A010 | ACH | T3_ipsi | 261 | Pleural remotor/abductor MN | | T3_ipsi | | | |
| DNa11 | 22 | IN19A003 | GABA | T3_ipsi | 1140 | Sternal posterior rotator MN | | T3_ipsi | | | |
| DNa11 | 22 | IN19A003 | GABA | T3_ipsi | 383 | Pleural remotor/abductor MN | | T3_ipsi | | | |
| DNa11 | 38.5 | IN04B074 | ACH | T3_ipsi | 445 | Sternal anterior rotator MN | | T3_ipsi | | | |
| DNa11 | 21 | IN01A018 | ACH | T1_ipsi | 195 | Pleural remotor/abductor MN | | T1_cont | | | |
| DNa11 | 21 | IN01A018 | ACH | T1_ipsi | 52 | Sternal posterior rotator MN | | T1_cont | | | |
| DNa11 | 34 | IN07B006 | ACH | T1_ipsi | 174 | Sternal posterior rotator MN | | T2_cont | | | |
| DNa11 | 34 | IN07B006 | ACH | T1_ipsi | 460 | Sternal posterior rotator MN | | T3_cont | | | |
| DNa11 | 41 | IN01A023 | ACH | T3_ipsi | 421 | Pleural remotor/abductor MN | | T3_cont | | | |
| DNa11 | 41 | IN01A023 | ACH | T3_ipsi | 319 | Sternal posterior rotator MN | | T3_cont | | | |
| DNa11 | 56 | LUL130 (INXXX468) | ACH | T1_ipsi | | | | | | | |
| DNa11 | 62.5 | LUL130 (INXXX468) | ACH | T1_ipsi | | | | | | | |
| DNa11 | 134 | LUL130 (INXXX468) | ACH | T2_ipsi | | | | | | | |
| IN03A010 | 51.5 | LUL130 (INXXX468) | ACH | T1_ipsi | | | | | | | |
| IN03A010 | 56.5 | LUL130 (INXXX468) | ACH | T2_ipsi | | | | | | | |
| IN03A010 | 82 | LUL130 (INXXX468) | ACH | T3_ipsi | | | | | | | |
| IN04B074 | 251 | IN19A003 | ACH | T3_ipsi | | | | | | | |

Supplementary Table 2

| | | | | | | | | | | | |
|---|---|---|---|---|---|---|---|---|---|---|---|
| LUL130 (INXXX468) | 141 | IN03A010 | ACH | T1_ipsi | | | | | | | |
| LUL130 (INXXX468) | 201 | IN03A010 | ACH | T2_ipsi | | | | | | | |
| LUL130 (INXXX468) | 51.5 | IN19A003 | ACH | T2_ipsi | | | | | | | |
| LUL130 (INXXX468) | 161 | IN03A010 | ACH | T3_ipsi | | | | | | | |
| LUL130 (INXXX468) | 23.5 | IN04B074 | ACH | T3_ipsi | | | | | | | |
| LUL130 (INXXX468) | 128 | IN19A003 | ACH | T3_ipsi | | | | | | | |
| From: DNa11_R(18107) | | | | | | | | | | | |
| type_pre | pre_id | weight | post_id | type_post | PMN_nt | PMN neuropil | PMN->MN weight | MN type | | MN neuropil | MN_id |
| DNa11_R | 18107 | 98 | 12050 | IN03A010 | ACH | T1_R | 355 | Fe reductor MN | | T1_R | 11226; 12891; 18983; 24050; 28884; 29484 |
| DNa11_R | 18107 | 98 | 12050 | IN03A010 | ACH | T1_R | 70 | Pleural remotor/abductor MN | | T1_R | 14346; 16419 |
| DNa11_R | 18107 | 45 | 12769 | IN03A010 | ACH | T2_R | 390 | Sternal posterior rotator MN | | T2_R | 11529; 16035; 102786; 104697 |
| DNa11_R | 18107 | 45 | 12769 | IN03A010 | ACH | T2_R | 303 | Pleural remotor/abductor MN | | T2_R | 19206; 166337; 41482 |
| DNa11_R | 18107 | 26 | 10144 | IN19A003 | GABA | T2_R | 1284 | Sternal posterior rotator MN | | T2_R | 11529; 102786; 16035; 104697 |
| DNa11_R | 18107 | 26 | 10144 | IN19A003 | GABA | T2_R | 219 | Pleural remotor/abductor MN | | T2_R | 19206; 166337; 41482 |
| DNa11_R | 18107 | 57 | 13908 | IN03A010 | ACH | T3_R | 518 | Sternal posterior rotator MN | | T3_R | 11400; 154133; 28219; 17947 |
| DNa11_R | 18107 | 57 | 13908 | IN03A010 | ACH | T3_R | 261 | Pleural remotor/abductor MN | | T3_R | 15262; 14596 |
| DNa11_R | 18107 | 34 | 10372 | IN19A003 | GABA | T3_R | 1140 | Sternal posterior rotator MN | | T3_R | 11400; 154133; 17947; 28219 |
| DNa11_R | 18107 | 34 | 10372 | IN19A003 | GABA | T3_R | 383 | Pleural remotor/abductor MN | | T3_R | 15262; 14596 |
| DNa11_R | 18107 | 50 (20; 12; 10; 8) | 26757; 19661; 27472; 26238 | IN04B074 | ACH | T3_R | 445 | Sternal anterior rotator MN | | T3_R | 11022; 11562 |
| DNa11_R | 18107 | 23 | 24896 | IN01A018 | ACH | T1_R | 195 | Pleural remotor/abductor MN | | T1_L | 13628; 12096 |
| DNa11_R | 18107 | 23 | 24896 | IN01A018 | ACH | T1_R | 52 | Sternal posterior rotator MN | | T1_L | 29689; 19667; 163719 |
| DNa11_R | 18107 | 23 | 10297 | IN07B006 | ACH | T2_R | 122 | Sternal posterior rotator MN | | T2_L | 15883; 49886; 154769; 172408 |
| DNa11_R | 18107 | 23 | 10297 | IN07B006 | ACH | T3_R | 59 | Sternal posterior rotator MN | | T3_L | 15278; 11162403133; 14449 |
| DNa11_R | 18107 | 50 | 11918 | IN01A023 | ACH | T3_R | 298 | Pleural remotor/abductor MN | | T3_L | 18286; 14168 |
| DNa11_R | 18107 | 50 | 11918 | IN01A023 | ACH | T3_R | 8 | Sternal posterior rotator MN | | T3_L | 15278; 27258; 154133; 11400; 11162403133 |
| DNa11_R | 18107 | 64 | 14650; 14230 | LUL130 (INXXX468) | ACH | T1_R | | | | | |
| DNa11_R | 18107 | 54 | 13511; 30260 | LUL130 (INXXX468) | ACH | T2_R | | | | | |
| DNa11_R | 18107 | 148 | 13293; 13574 | LUL130 (INXXX468) | ACH | T3_R | | | | | |
| IN03A010 | 12050 | 58 | 14650; 14230 | LUL130 (INXXX468) | ACH | T1_R | | | | | |
| IN03A010 | 12769 | 60 | 13511; 30260 | LUL130 (INXXX468) | ACH | T2_R | | | | | |
| IN03A010 | 13908 | 82 | 13293; 13574 | LUL130 (INXXX468) | ACH | T3_R | | | | | |
| IN04B074 | 26757; 19661; 27472; 26238 | 397 | 10372 | IN19A003 | ACH | T3_R | | | | | |
| LUL130 (INXXX468) | 14650; 14230 | 151 | 12050 | IN03A010 | ACH | T1_R | | | | | |
| LUL130 (INXXX468) | 13511; 30260 | 203 | 12769 | IN03A010 | ACH | T2_R | | | | | |
| LUL130 (INXXX468) | 13511; 30260 | 50 | 10144 | IN19A003 | ACH | T2_R | | | | | |
| LUL130 (INXXX468) | 13293; 13574 | 149 | 13908 | IN03A010 | ACH | T3_R | | | | | |
| LUL130 (INXXX468) | 13293; 13574 | 25 | 26757; 19661; 27472; 26238 | IN04B074 | ACH | T3_R | | | | | |
| LUL130 (INXXX468) | 13293; 13574 | 152 | 10372 | IN19A003 | ACH | T3_R | | | | | |
| From: DNa11_L(17155) | | | | | | | | | | | |

Supplementary Table 2

| type_pre | pre_id | weight | post_id | type_post | PMN_nt | PMN neuropil | PMN->MN weight | MN type | | MN neuropil | MN_id |
|---|---|---|---|---|---|---|---|---|---|---|---|
| DNa11_L | 17155 | 71 | 100492 | IN03A010 | ACH | T1_L | 401 | Fe reductor MN | | T1_L | 14949; 13920; 156857; 21944; 226276; 219894 |
| DNa11_L | 17155 | 71 | 100492 | IN03A010 | ACH | T1_L | 76 | Pleural remotor/abductor MN | | T1_L | 13628; 12096 |
| DNa11_L | 17155 | 64 | 12160 | IN03A010 | ACH | T2_L | 118 | Sternal posterior rotator MN | | T2_L | 49886; 15883; 172408; 154769 |
| DNa11_L | 17155 | 64 | 12160 | IN03A010 | ACH | T2_L | 222 | Pleural remotor/abductor MN | | T2_L | 17091; 21255; 219853 |
| DNa11_L | 17155 | 37 | 10156 | IN19A003 | GABA | T2_L | 683 | Sternal posterior rotator MN | | T2_L | 49886; 15883; 154796; 172408 |
| DNa11_L | 17155 | 37 | 10156 | IN19A003 | GABA | T2_L | 185 | Pleural remotor/abductor MN | | T2_L | 17091; 21255; 219853 |
| DNa11_L | 17155 | 34 | 12605 | IN03A010 | ACH | T3_L | 20 | Sternal posterior rotator MN | | T3_L | 15278; 27258; 14449; 11162403133 |
| DNa11_L | 17155 | 34 | 12605 | IN03A010 | ACH | T3_L | 212 | Pleural remotor/abductor MN | | T3_L | 14168; 18286 |
| DNa11_L | 17155 | 10 | 11303 | IN19A003 | GABA | T3_L | 84 | Sternal posterior rotator MN | | T3_L | 15278; 14449; 11162403133 |
| DNa11_L | 17155 | 10 | 11303 | IN19A003 | GABA | T3_L | 275 | Pleural remotor/abductor MN | | T3_L | 18286; 14168 |
| DNa11_L | 17155 | 27(14; 13) | 23500; 26556 | IN04B074 | ACH | T3_L | 210 | Sternal anterior rotator MN | | T3_L | 164741; 18471 |
| DNa11_L | 17155 | 19 | 11688 | IN01A018 | ACH | T1_L | 163 | Pleural remotor/abductor MN | | T1_R | 14346; 16419 |
| DNa11_L | 17155 | 19 | 11688 | IN01A018 | ACH | T1_L | 47 | Sternal posterior rotator MN | | T1_R | 21241; 25767; 13942 |
| DNa11_L | 17155 | 45 | 10340 | IN07B006 | ACH | T2_L | 174 | Sternal posterior rotator MN | | T2_R | 102786; 104697; 16035; 11529 |
| DNa11_L | 17155 | 45 | 10340 | IN07B006 | ACH | T3_L | 460 | Sternal posterior rotator MN | | T3_R | 154133; 11400; 17947; 28219 |
| DNa11_L | 17155 | 32 | 11420 | IN01A023 | ACH | T3_L | 421 | Pleural remotor/abductor MN | | T3_R | 15262; 14596 |
| DNa11_L | 17155 | 32 | 11420 | IN01A023 | ACH | T3_L | 319 | Sternal posterior rotator MN | | T3_R | 11400; 154133; 17947 |
| DNa11_L | 17155 | 48 | 13246; 15115 | LUL130 (INXXX468) | ACH | T1_L | | | | | |
| DNa11_L | 17155 | 71 | 14432; 14588 | LUL130 (INXXX468) | ACH | T2_L | | | | | |
| DNa11_L | 17155 | 120 | 14084; 13323 | LUL130 (INXXX468) | ACH | T3_L | | | | | |
| IN03A010 | 100492 | 45 | 13246; 15115 | LUL130 (INXXX468) | ACH | T1_L | | | | | |
| IN03A010 | 12160 | 53 | 14432; 14588 | LUL130 (INXXX468) | ACH | T2_L | | | | | |
| IN03A010 | 12605 | 82 | 14084; 13323 | LUL130 (INXXX468) | ACH | T3_L | | | | | |
| IN04B074 | 23500; 26556 | 105 | 11303 | IN19A003 | ACH | T3_L | | | | | |
| LUL130 (INXXX468) | 13246; 15115 | 131 | 100492 | IN03A010 | ACH | T1_L | | | | | |
| LUL130 (INXXX468) | 14432; 14588 | 199 | 12160 | IN03A010 | ACH | T2_L | | | | | |
| LUL130 (INXXX468) | 14432; 14588 | 53 | 10156 | IN19A003 | ACH | T2_L | | | | | |
| LUL130 (INXXX468) | 14084; 13323 | 173 | 12605 | IN03A010 | ACH | T3_L | | | | | |
| LUL130 (INXXX468) | 14084; 13323 | 22 | 23500; 26556 | IN04B074 | ACH | T3_L | | | | | |
| LUL130 (INXXX468) | 14084; 13323 | 104 | 11303 | IN19A003 | ACH | T3_L | | | | | |
| Top 100 of bi-synaptic DNa11->MN connections ranked by indirect weights (DN->PMN relative weight * PMN->MN relative weight) | | | | | | | | | | | |
| MNs are grouped in types; the relative weights (neuron A->B) are calculated as: sqrt ((synapse number A->B / total postsynapse number B) * (synapse number A->B / total presynapse number A)) | | | | | | | | | | | |
| Rank | DNa11 | DN->PMN weight | PMN | PMN nt type | PMN type | PMN->MN weight | MN type | | MN neuropil | Indirect weight | |
| 1 | 18107 | 0.011964461 | 12050 | ACH | IN03A010_T1_R | 0.015624239 | Fe reductor MN | | T1_R | 0.000802737 | |
| 2 | 18107 | 0.008429233 | 13908 | ACH | IN03A010_T3_R | 0.023706336 | Sternal posterior rotator MN | | T3_R | 0.00057842 | |
| 3 | 18107 | 0.003254726 | 10372 | GABA | IN19A003_T3_R | 0.056603097 | Sternal posterior rotator MN | | T3_R | 0.000514187 | |
| 4 | 18107 | 0.002254821 | 10144 | GABA | IN19A003_T2_R | 0.052540467 | Sternal posterior rotator MN | | T2_R | 0.000384639 | |
| 5 | 18107 | 0.008497227 | 11918 | ACH | IN01A023_T3_R | 0.018155201 | Pleural remotor/abductor MN | | T3_L | 0.000306491 | |
| 6 | 18107 | 0.005972518 | 12769 | ACH | IN03A010_T2_R | 0.023543783 | Sternal posterior rotator MN | | T2_R | 0.000260288 | |
| 7 | 18107 | 0.007762135 | 26757 | ACH | IN04B074_T3_R | 0.018497913 | Sternal anterior rotator MN | | T3_R | 0.000256639 | |
| 8 | 18107 | 0.005972518 | 12769 | ACH | IN03A010_T2_R | 0.030829691 | Pleural remotor/abductor MN | | T2_R | 0.000256376 | |
| 9 | 18107 | 0.008429233 | 13908 | ACH | IN03A010_T3_R | 0.01563494 | Pleural remotor/abductor MN | | T3_R | 0.000254164 | |
| 10 | 18107 | 0.004926341 | 24896 | ACH | IN01A018_T1_R | 0.023967828 | Pleural remotor/abductor MN | | T1_L | 0.000187935 | |
| 11 | 18107 | 0.003430279 | 19661 | ACH | IN04B074_T3_R | 0.027278872 | Sternal anterior rotator MN | | T3_R | 0.00017769 | |
| 12 | 18107 | 0.011964461 | 12050 | ACH | IN03A010_T1_R | 0.007938315 | Pleural remotor/abductor MN | | T1_R | 0.000176756 | |
| 13 | 18107 | 0.003488682 | 27472 | ACH | IN04B074_T3_R | 0.022194812 | Sternal anterior rotator MN | | T3_R | 0.000154269 | |

Supplementary Table 2

| | | | | | | | | | |
|---|---|---|---|---|---|---|---|---|---|
| 14 | 18107 | 0.011964461 | 12050 | ACH | IN03A010_T1_R | 0.003981367 | Sternal posterior rotator MN | T1_R | 0.000152911 |
| 15 | 18107 | 0.019975037 | 10148 | ACH | IN07B006_T2_R | 0.002690708 | Ti flexor MN | T3_L | 0.000151841 |
| 16 | 18107 | 0.008429233 | 13908 | ACH | IN03A010_T3_R | 0.009002453 | Fe reductor MN | T3_R | 0.000133754 |
| 17 | 18107 | 0.011964461 | 12050 | ACH | IN03A010_T1_R | 0.005016159 | Ta levator MN | T1_R | 0.000131929 |
| 18 | 18107 | 0.003254726 | 10372 | GABA | IN19A003_T3_R | 0.029014583 | Pleural remotor/abductor MN | T3_R | 0.000126647 |
| 19 | 18107 | 0.040194378 | 12152 | GABA | AN06B015_T3_R | 0.003104955 | Sternal anterior rotator MN | T3_L | 0.000124802 |
| 20 | 18107 | 0.040194378 | 12152 | GABA | AN06B015_T3_R | 0.002926946 | Itm MN | T2_L | 0.000117647 |
| 21 | 18107 | 0.003395363 | 154210 | GLUT | IN08A037_T3_R | 0.020381707 | Pleural remotor/abductor MN | T3_R | 0.000103907 |
| 22 | 18107 | 0.003119139 | 26238 | ACH | IN04B074_T3_R | 0.016238103 | Sternal anterior rotator MN | T3_R | 9.388887045234709e-05 |
| 23 | 18107 | 0.008429233 | 13908 | ACH | IN03A010_T3_R | 0.010813781 | MNhl65 | T3_R | 9.11518767374764e-05 |
| 24 | 18107 | 0.008497227 | 11918 | ACH | IN01A023_T3_R | 0.010371281 | Fe reductor MN | T3_L | 8.81271268761195e-05 |
| 25 | 18107 | 0.002739829 | 26165 | GLUT | IN08A037_T3_R | 0.015936689 | Pleural remotor/abductor MN | T3_R | 8.453689803140002e-05 |
| 26 | 18107 | 0.040194378 | 12152 | GABA | AN06B015_T3_R | 0.002092132 | Sternal anterior rotator MN | T2_L | 8.409196199631815e-05 |
| 27 | 18107 | 0.008429233 | 13908 | ACH | IN03A010_T3_R | 0.009917196 | Sternal adductor MN | T3_R | 8.359435277766387e-05 |
| 28 | 18107 | 0.002761963 | 26784 | GLUT | IN08A048_T3_R | 0.012246926 | Sternotrochanter MN | T3_R | 8.13307657684883e-05 |
| 29 | 18107 | 0.005972518 | 12769 | ACH | IN03A010_T2_R | 0.013528748 | MNml29 | T2_R | 8.080069207696594e-05 |
| 30 | 18107 | 0.003254726 | 10372 | GABA | IN19A003_T3_R | 0.024588003 | MNhl29 | T3_R | 8.002720788628295e-05 |
| 31 | 18107 | 0.004926341 | 24896 | ACH | IN01A018_T1_R | 0.01119191 | Sternal posterior rotator MN | T1_L | 7.569110948000916e-05 |
| 32 | 18107 | 0.019975037 | 10148 | ACH | IN07B006_T2_R | 0.002685911 | Acc. ti flexor MN | T2_L | 7.381968562344753e-05 |
| 33 | 18107 | 0.002254821 | 10144 | GABA | IN19A003_T2_R | 0.019399094 | Pleural remotor/abductor MN | T2_R | 7.360129600996968e-05 |
| 34 | 18107 | 0.002739829 | 26165 | GLUT | IN08A037_T3_R | 0.009283226 | Sternal posterior rotator MN | T3_R | 7.324648179066309e-05 |
| 35 | 18107 | 0.001792405 | 28526 | ACH | IN01A025_T1_L | 0.021529968 | Acc. ti flexor MN | T1_R | 7.15998235226849e-05 |
| 36 | 18107 | 0.002403964 | 14144 | ACH | IN01A025_T1_R | 0.015082513 | Tergopleural/Pleural promotor MN | T1_L | 6.855611045938305e-05 |
| 37 | 18107 | 0.002246749 | 21519 | ACH | IN04B081_T1_R | 0.018287105 | Sternal anterior rotator MN | T1_R | 6.620967523444615e-05 |
| 38 | 18107 | 0.003254726 | 10372 | GABA | IN19A003_T3_R | 0.019220387 | Fe reductor MN | T3_R | 6.58700588893033e-05 |
| 39 | 18107 | 0.002403964 | 14144 | ACH | IN01A025_T1_R | 0.010032837 | Acc. ti flexor MN | T1_L | 6.495052095445025e-05 |
| 40 | 18107 | 0.003099912 | 17584 | ACH | IN04B081_T1_R | 0.012417626 | Sternal anterior rotator MN | T1_R | 6.460385877340977e-05 |
| 41 | 18107 | 0.002403964 | 14144 | ACH | IN01A025_T1_R | 0.017046669 | Ta depressor MN | T1_L | 6.423063256847277e-05 |
| 42 | 18107 | 0.008429233 | 13908 | ACH | IN03A010_T3_R | 0.007510787 | MNhl29 | T3_R | 6.331017354500026e-05 |
| 43 | 18107 | 0.003625545 | 22934 | GABA | IN09A077_T1_R | 0.005997109 | Acc. ti flexor MN | T1_R | 6.204082889842694e-05 |
| 44 | 18107 | 0.002083866 | 19069 | ACH | IN03A075_T1_R | 0.022311137 | Sternal posterior rotator MN | T1_R | 6.203090809390296e-05 |
| 45 | 18107 | 0.040194378 | 12152 | GABA | AN06B015_T3_R | 0.001542275 | MNhl59 | T3_R | 6.199078109573212e-05 |
| 46 | 18107 | 0.00274913 | 10297 | ACH | IN07B006_T1_R | 0.011053527 | Sternal posterior rotator MN | T3_L | 6.125421688379511e-05 |
| 47 | 18107 | 0.002678691 | 153583 | GABA | IN09A064_T3_R | 0.011251836 | Acc. ti flexor MN | T3_R | 5.995388398962555e-05 |
| 48 | 18107 | 0.009093735 | 10844 | ACH | IN19A017_T1_R | 0.003990121 | Sternotrochanter MN | T3_R | 5.889305865846302e-05 |
| 49 | 18107 | 0.00274913 | 10297 | ACH | IN07B006_T1_R | 0.007883252 | Sternal posterior rotator MN | T2_L | 5.6271614117866e-05 |
| 50 | 18107 | 0.003099912 | 17584 | ACH | IN04B081_T1_R | 0.010196503 | Tergopleural/Pleural promotor MN | T1_R | 5.392396923526933e-05 |
| 51 | 18107 | 0.011825028 | 11098 | ACH | AN12A003_A1_R | 0.004285567 | Itm1-tibia MN | T2_R | 5.0676945501486974e-05 |
| 52 | 18107 | 0.001792405 | 28526 | ACH | IN01A025_T1_L | 0.014358315 | Tergopleural/Pleural promotor MN | T1_R | 4.996196514494235e-05 |
| 53 | 18107 | 0.002254821 | 10144 | GABA | IN19A003_T2_R | 0.021855191 | MNml29 | T2_R | 4.927954813387772e-05 |
| 54 | 18107 | 0.003254726 | 10372 | GABA | IN19A003_T3_R | 0.014515781 | MNhl65 | T3_R | 4.724488730722653e-05 |
| 55 | 18107 | 0.019975037 | 10148 | ACH | IN07B006_T2_R | 0.002359491 | Ti flexor MN | T2_L | 4.7130927459751656e-05 |
| 56 | 18107 | 0.003898452 | 18832 | GABA | IN09A064_T2_R | 0.006893313 | Acc. ti flexor MN | T2_R | 4.668355236679448e-05 |
| 57 | 18107 | 0.001792405 | 28526 | ACH | IN01A025_T1_L | 0.014164417 | Ta depressor MN | T1_R | 4.448001034065423e-05 |
| 58 | 18107 | 0.005972518 | 12769 | ACH | IN03A010_T2_R | 0.004541686 | Fe reductor MN | T2_R | 4.408418759972733e-05 |
| 59 | 18107 | 0.004867446 | 19930 | ACH | IN08B082_T3_R | 0.009034021 | MNnm08 | T1_L | 4.397260637849802e-05 |
| 60 | 18107 | 0.005972518 | 12769 | ACH | IN03A010_T2_R | 0.007332387 | MNml81 | T2_R | 4.3792814207945725e-05 |
| 61 | 18107 | 0.003395363 | 154210 | GLUT | IN08A037_T3_R | 0.007312364 | Sternal posterior rotator MN | T3_R | 4.3187393450803234e-05 |
| 62 | 18107 | 0.002761963 | 26784 | GLUT | IN08A048_T3_R | 0.012730529 | Tr extensor MN | T3_R | 4.267983357095332e-05 |
| 63 | 18107 | 0.003395363 | 154210 | GLUT | IN08A037_T3_R | 0.012506716 | Fe reductor MN | T3_R | 4.246484047524097e-05 |
| 64 | 18107 | 0.001238776 | 21782 | ACH | IN04B081_T1_R | 0.017499332 | Sternal anterior rotator MN | T1_R | 3.94578090606052e-05 |
| 65 | 18107 | 0.003119139 | 26238 | ACH | IN04B074_T3_R | 0.008735446 | Sternotrochanter MN | T3_R | 3.939788134845962e-05 |
| 66 | 18107 | 0.001809594 | 13013 | GLUT | AN06B023_T1_R | 0.020978537 | MNnm03 | T1_L | 3.796263292352113e-05 |
| 67 | 18107 | 0.008429233 | 13908 | ACH | IN03A010_T3_R | 0.004436986 | Ti extensor MN | T3_R | 3.740038646416034e-05 |
| 68 | 18107 | 0.002083866 | 19069 | ACH | IN03A075_T1_R | 0.011055551 | Pleural remotor/abductor MN | T1_R | 3.640741108154884e-05 |
| 69 | 18107 | 0.003430279 | 19661 | ACH | IN04B074_T3_R | 0.010582653 | Acc. tr flexor MN | T3_R | 3.630145812999455e-05 |
| 70 | 18107 | 0.007762135 | 26757 | ACH | IN04B074_T3_R | 0.00262562 | Sternotrochanter MN | T3_R | 3.464681709852631e-05 |
| 71 | 18107 | 0.002739829 | 26165 | GLUT | IN08A037_T3_R | 0.012500297 | Fe reductor MN | T3_R | 3.42486737330443e-05 |
| 72 | 18107 | 0.001208209 | 18960 | ACH | IN03A075_T2_R | 0.023808972 | Sternal posterior rotator MN | T2_R | 3.379986153731507e-05 |
| 73 | 18107 | 0.005506258 | 10602 | ACH | ANXXX037_A1_R | 0.002864709 | Sternotrochanter MN | T3_R | 3.3670859170451426e-05 |
| 74 | 18107 | 0.003411178 | 10512 | GLUT | AN02A002_T2_R | 0.009823216 | MNnm14 | T1_R | 3.350873423696119e-05 |

Supplementary Table 2

| | | | | | | | | | |
|---|---|---|---|---|---|---|---|---|---|
| 75 | 18107 | 0.011964461 | 12050 | ACH | IN03A010_T1_R | 0.002797474 | Sternal adductor MN | T1_R | 3.347026601463947e-05 |
| 76 | 18107 | 0.005972518 | 12769 | ACH | IN03A010_T2_R | 0.00527426 | Ti extensor MN | T2_R | 3.1500615031779594e-05 |
| 77 | 18107 | 0.001876835 | 19218 | ACH | IN03A075_T1_R | 0.016675221 | Sternal posterior rotator MN | T1_R | 3.1296639978770173e-05 |
| 78 | 18107 | 0.004502541 | 24440 | GABA | IN09A045_T3_R | 0.003564609 | Acc. ti flexor MN | T3_R | 3.0491274244002734e-05 |
| 79 | 18107 | 0.003411178 | 10512 | GLUT | AN02A002_T2_R | 0.008927345 | FNM2 | T1_R | 3.0452760034033308e-05 |
| 80 | 18107 | 0.008497227 | 11918 | ACH | IN01A023_T3_R | 0.003572116 | MNhl65 | T3_L | 3.0353078369304124e-05 |
| 81 | 18107 | 0.008497227 | 11918 | ACH | IN01A023_T3_R | 0.003559321 | MNhl29 | T3_L | 3.024435722225374e-05 |
| 82 | 18107 | 0.007762135 | 26757 | ACH | IN04B074_T3_R | 0.003887436 | MNhl59 | T3_R | 3.01748063900627e-05 |
| 83 | 18107 | 0.009093735 | 10844 | ACH | IN19A017_T1_R | 0.003284015 | Tr extensor MN | T3_R | 2.98639609045432e-05 |
| 84 | 18107 | 0.009093735 | 10844 | ACH | IN19A017_T1_R | 0.003263015 | Ti flexor MN | T3_R | 2.967299156286431e-05 |
| 85 | 18107 | 0.008497227 | 11918 | ACH | IN01A023_T3_R | 0.003370466 | Sternal adductor MN | T3_L | 2.8639610364034906e-05 |
| 86 | 18107 | 0.007762135 | 26757 | ACH | IN04B074_T3_R | 0.003668694 | Acc. tr flexor MN | T3_R | 2.8476896114453373e-05 |
| 87 | 18107 | 0.001903712 | 22780 | ACH | IN04B081_T2_R | 0.012222663 | Sternal anterior rotator MN | T2_R | 2.8436168150434857e-05 |
| 88 | 18107 | 0.001792405 | 28526 | ACH | IN01A025_T1_L | 0.012353329 | Tr extensor MN | T1_R | 2.8278578821498062e-05 |
| 89 | 18107 | 0.005298399 | 25626 | GABA | IN12B048_T3_L | 0.005293382 | MNxm02 | T3_R | 2.804645360229386e-05 |
| 90 | 18107 | 0.001876835 | 19218 | ACH | IN03A075_T1_R | 0.007589156 | Fe reductor MN | T1_R | 2.799682252168218e-05 |
| 91 | 18107 | 0.009093735 | 10844 | ACH | IN19A017_T1_R | 0.002997342 | MNhl88 | T3_R | 2.7257033407601257e-05 |
| 92 | 18107 | 0.008497227 | 11918 | ACH | IN01A023_T3_R | 0.001891346 | Sternal posterior rotator MN | T3_L | 2.7220591629648098e-05 |
| 93 | 18107 | 0.003099912 | 17584 | ACH | IN04B081_T1_R | 0.008187802 | Sternal adductor MN | T1_R | 2.5381463905381486e-05 |
| 94 | 18107 | 0.019975037 | 10148 | ACH | IN07B006_T2_R | 0.001203881 | MNml10 | T2_L | 2.4047576648173248e-05 |
| 95 | 18107 | 0.009337508 | 16639 | ACH | IN08B058_T3_R | 0.002540625 | Tr flexor MN | T3_L | 2.3723110126328798e-05 |
| 96 | 18107 | 0.006612338 | 23664 | GABA | IN09A065_T2_R | 0.003581703 | Acc. ti flexor MN | T2_R | 2.3683432700913637e-05 |
| 97 | 18107 | 0.009337508 | 16639 | ACH | IN08B058_T3_R | 0.002500001 | Sternal posterior rotator MN | T1_L | 2.3343781069169103e-05 |
| 98 | 18107 | 0.009093735 | 10844 | ACH | IN19A017_T1_R | 0.002530598 | Itm MN | T3_R | 2.3012587519110144e-05 |
| 99 | 18107 | 0.009337508 | 16639 | ACH | IN08B058_T3_R | 0.002451789 | FNM2 | T1_L | 2.2893598478107274e-05 |
| 100 | 18107 | 0.00644259 | 10725 | GABA | IN05B008_A1_R | 0.003519721 | iii1 MN | T2_L | 2.267611933388745e-05 |
| Rank | DNa11 | DN->PMN weight | PMN | PMN nt type | PMN type | PMN->MN weight | MN type | MN neuropil | Indirect weight |
| 1 | 17155 | 0.008855625 | 100492 | ACH | IN03A010_T1_L | 0.018998579 | Fe reductor MN | T1_L | 0.000658462 |
| 2 | 17155 | 0.003140606 | 10156 | GABA | IN19A003_T2_L | 0.047116091 | Sternal posterior rotator MN | T2_L | 0.000393637 |
| 3 | 17155 | 0.008842737 | 12160 | ACH | IN03A010_T2_L | 0.025911077 | Pleural remotor/abductor MN | T2_L | 0.000340232 |
| 4 | 17155 | 0.005620716 | 11420 | ACH | IN01A023_T3_L | 0.022780866 | Pleural remotor/abductor MN | T3_R | 0.000246773 |
| 5 | 17155 | 0.00541806 | 10340 | ACH | IN07B006_T1_L | 0.017058811 | Sternal posterior rotator MN | T3_R | 0.000224366 |
| 6 | 17155 | 0.008842737 | 12160 | ACH | IN03A010_T2_L | 0.008305318 | Sternal posterior rotator MN | T2_L | 0.000196186 |
| 7 | 17155 | 0.004150203 | 23500 | ACH | IN04B074_T3_L | 0.024222062 | Sternal anterior rotator MN | T3_L | 0.000182141 |
| 8 | 17155 | 0.005620716 | 11420 | ACH | IN01A023_T3_L | 0.017882199 | Sternal posterior rotator MN | T3_R | 0.000175922 |
| 9 | 17155 | 0.004680959 | 12605 | ACH | IN03A010_T3_L | 0.020269335 | Pleural remotor/abductor MN | T3_L | 0.000144032 |
| 10 | 17155 | 0.004426243 | 26556 | ACH | IN04B074_T3_L | 0.018774541 | Sternal anterior rotator MN | T3_L | 0.000143314 |
| 11 | 17155 | 0.004063319 | 11688 | ACH | IN01A018_T1_L | 0.02532031 | Pleural remotor/abductor MN | T1_R | 0.00014282 |
| 12 | 17155 | 0.00541806 | 10340 | ACH | IN07B006_T1_L | 0.010592576 | Sternal posterior rotator MN | T2_R | 0.000130446 |
| 13 | 17155 | 0.008855625 | 100492 | ACH | IN03A010_T1_L | 0.004556033 | Sternal posterior rotator MN | T1_L | 0.000130045 |
| 14 | 17155 | 0.008855625 | 100492 | ACH | IN03A010_T1_L | 0.010331217 | Pleural remotor/abductor MN | T1_L | 0.000120656 |
| 15 | 17155 | 0.03117516 | 13064 | GABA | AN06B015_T3_L | 0.002462642 | Sternal anterior rotator MN | T3_R | 0.000120413 |
| 16 | 17155 | 0.00250329 | 16079 | ACH | IN03A066_T1_L | 0.019654497 | Sternal posterior rotator MN | T1_L | 0.000120388 |
| 17 | 17155 | 0.008842737 | 12160 | ACH | IN03A010_T2_L | 0.006061674 | Fe reductor MN | T2_L | 0.000112533 |
| 18 | 17155 | 0.00250329 | 16079 | ACH | IN03A066_T1_L | 0.023769787 | Pleural remotor/abductor MN | T1_L | 0.000101636 |
| 19 | 17155 | 0.003140606 | 10156 | GABA | IN19A003_T2_L | 0.01668268 | Pleural remotor/abductor MN | T2_L | 0.00010095 |
| 20 | 17155 | 0.008842737 | 12160 | ACH | IN03A010_T2_L | 0.009646832 | MNml81 | T2_L | 8.530439463219817e-05 |
| 21 | 17155 | 0.004680959 | 12605 | ACH | IN03A010_T3_L | 0.009654144 | Fe reductor MN | T3_L | 8.52234746629642e-05 |
| 22 | 17155 | 0.005932525 | 19808 | ACH | IN08B082_T3_L | 0.01397465 | MNnm08 | T1_R | 8.290495143987929e-05 |
| 23 | 17155 | 0.001961027 | 28526 | ACH | IN01A025_T1_L | 0.021529968 | Acc. ti flexor MN | T1_R | 7.833563569362995e-05 |
| 24 | 17155 | 0.002077411 | 19372 | ACH | IN04B081_T2_L | 0.025455628 | Sternal anterior rotator MN | T2_L | 7.656820187192242e-05 |
| 25 | 17155 | 0.005620716 | 11420 | ACH | IN01A023_T3_L | 0.012977204 | MNhl65 | T3_R | 7.294117010546535e-05 |
| 26 | 17155 | 0.011621455 | 10206 | ACH | IN07B006_T2_L | 0.003345274 | Acc. ti flexor MN | T3_R | 7.237703505727192e-05 |
| 27 | 17155 | 0.005620716 | 11420 | ACH | IN01A023_T3_L | 0.010373133 | Fe reductor MN | T3_R | 7.136848917334155e-05 |
| 28 | 17155 | 0.001163509 | 10890 | GABA | IN08A006_T3_L | 0.030598094 | Sternal anterior rotator MN | T3_L | 7.013565692882269e-05 |
| 29 | 17155 | 0.003140606 | 10156 | GABA | IN19A003_T2_L | 0.020772841 | MNml29 | T2_L | 6.523931212593097e-05 |
| 30 | 17155 | 0.013680034 | 13509 | ACH | ANXXX131_T1_R | 0.004666911 | Fe reductor MN | T1_L | 6.384350346445066e-05 |
| 31 | 17155 | 0.003019152 | 21569 | ACH | IN04B081_T2_L | 0.014133225 | Sternal anterior rotator MN | T2_L | 5.945213603197117e-05 |
| 32 | 17155 | 0.011840432 | 192710 | ACH | IN19A017_T1_L | 0.004674728 | MNhl59 | T3_L | 5.535079429939021e-05 |

Supplementary Table 2

| | | | | | | | | | | |
|---|---|---|---|---|---|---|---|---|---|---|
| 33 | 17155 | 0.009786432 | 11121 | ACH | AN12A003_A1_L | 0.005615082 | Itm MN | | T2_L | 5.4951620621400425e-05 |
| 34 | 17155 | 0.001961027 | 28526 | ACH | IN01A025_T1_L | 0.014358315 | Tergopleural/Pleural promotor MN | | T1_R | 5.46621780274645e-05 |
| 35 | 17155 | 0.001727189 | 26488 | GLUT | IN08A037_T3_L | 0.018592177 | Pleural remotor/abductor MN | | T3_L | 5.3924285726012736e-05 |
| 36 | 17155 | 0.001282818 | 11303 | GABA | IN19A003_T3_L | 0.022162902 | Sternal posterior rotator MN | | T3_L | 4.989869326042264e-05 |
| 37 | 17155 | 0.001282818 | 11303 | GABA | IN19A003_T3_L | 0.024331371 | Pleural remotor/abductor MN | | T3_L | 4.9873862958285785e-05 |
| 38 | 17155 | 0.004680959 | 12605 | ACH | IN03A010_T3_L | 0.005082223 | Sternal posterior rotator MN | | T3_L | 4.935898653331658e-05 |
| 39 | 17155 | 0.004029269 | 100945 | GABA | IN09A065_T2_L | 0.004407768 | Acc. ti flexor MN | | T2_L | 4.928823602030253e-05 |
| 40 | 17155 | 0.001554985 | 16059 | GABA | IN06B047_T3_R | 0.031687411 | hg4 MN | | T2_L | 4.927346205029806e-05 |
| 41 | 17155 | 0.001649311 | 27882 | ACH | IN04B074_T3_L | 0.015360437 | Sternal anterior rotator MN | | T3_L | 4.885501707102872e-05 |
| 42 | 17155 | 0.001961027 | 28526 | ACH | IN01A025_T1_L | 0.014164417 | Ta depressor MN | | T1_R | 4.866450382507485e-05 |
| 43 | 17155 | 0.004063319 | 11688 | ACH | IN01A018_T1_L | 0.010003476 | Sternal posterior rotator MN | | T1_R | 4.800700642534626e-05 |
| 44 | 17155 | 0.004426243 | 26556 | ACH | IN04B074_T3_L | 0.010396673 | MNhl59 | | T3_L | 4.6018199215166104e-05 |
| 45 | 17155 | 0.002444571 | 21236 | ACH | IN08B082_T3_L | 0.018781403 | MNnm08 | | T1_R | 4.591246764707814e-05 |
| 46 | 17155 | 0.011621455 | 10206 | ACH | IN07B006_T2_L | 0.002257032 | MNhl68 | | T3_R | 4.581795361120889e-05 |
| 47 | 17155 | 0.001578071 | 14144 | ACH | IN01A025_T1_R | 0.015082513 | Tergopleural/Pleural promotor MN | | T1_L | 4.500334969526986e-05 |
| 48 | 17155 | 0.004227602 | 22195 | GABA | IN09A045_T2_L | 0.007214872 | Acc. ti flexor MN | | T2_L | 4.3330939921654296e-05 |
| 49 | 17155 | 0.001578071 | 14144 | ACH | IN01A025_T1_R | 0.010032837 | Acc. ti flexor MN | | T1_L | 4.2636476716905354e-05 |
| 50 | 17155 | 0.011840432 | 192710 | ACH | IN19A017_T1_L | 0.002081579 | Sternotrochanter MN | | T3_L | 4.2463250528134055e-05 |
| 51 | 17155 | 0.001578071 | 14144 | ACH | IN01A025_T1_R | 0.017046669 | Ta depressor MN | | T1_L | 4.216390923081814e-05 |
| 52 | 17155 | 0.001239813 | 164298 | ACH | IN04B074_T3_L | 0.017015294 | Sternal anterior rotator MN | | T3_L | 4.1888918544641474e-05 |
| 53 | 17155 | 0.023269834 | 15569 | ACH | AN19B042_T1_L | 0.001791628 | Itm MN | | T1_R | 4.169089516510725e-05 |
| 54 | 17155 | 0.001363378 | 12007 | GABA | IN14B006_T3_L | 0.016476556 | Pleural remotor/abductor MN | | T3_R | 4.087914293822595e-05 |
| 55 | 17155 | 0.011840432 | 192710 | ACH | IN19A017_T1_L | 0.002061994 | Acc. ti flexor MN | | T3_L | 4.051480711725365e-05 |
| 56 | 17155 | 0.001848204 | 19833 | ACH | IN04B081_T1_L | 0.017755902 | Sternal anterior rotator MN | | T1_L | 3.974526261179265e-05 |
| 57 | 17155 | 0.00288459 | 25119 | GABA | IN09A042_T3_L | 0.007148393 | Acc. ti flexor MN | | T3_L | 3.918931613341322e-05 |
| 58 | 17155 | 0.003200476 | 13318 | ACH | IN20A.22A003_T2_ | 0.007331781 | Sternotrochanter MN | | T2_L | 3.8707043705034635e-05 |
| 59 | 17155 | 0.005932525 | 19808 | ACH | IN08B082_T3_L | 0.003630199 | Sternal posterior rotator MN | | T3_R | 3.738487397543345e-05 |
| 60 | 17155 | 0.001378388 | 23863 | GLUT | IN08A037_T3_L | 0.017223246 | Pleural remotor/abductor MN | | T3_L | 3.71043850153881e-05 |
| 61 | 17155 | 0.001554985 | 16059 | GABA | IN06B047_T3_R | 0.023831369 | MNwm35 | | T2_L | 3.705743108824026e-05 |
| 62 | 17155 | 0.011621455 | 10206 | ACH | IN07B006_T2_L | 0.00188477 | Ti flexor MN | | T3_R | 3.660067927346217e-05 |
| 63 | 17155 | 0.00357042 | 22953 | GABA | IN09A064_T1_L | 0.010247782 | Acc. ti flexor MN | | T1_L | 3.6588887505251876e-05 |
| 64 | 17155 | 0.03117516 | 13064 | GABA | AN06B015_T3_L | 0.001162941 | MNnm12 | | T1_R | 3.625486154361563e-05 |
| 65 | 17155 | 0.001282818 | 11303 | GABA | IN19A003_T3_L | 0.027786594 | MNhl29 | | T3_L | 3.564513586980291e-05 |
| 66 | 17155 | 0.011621455 | 10206 | ACH | IN07B006_T2_L | 0.001853132 | Ti flexor MN | | T2_R | 3.447181436129193e-05 |
| 67 | 17155 | 0.006661675 | 13246 | ACH | INXXX468_T1_L | 0.005086792 | Tr flexor MN | | T1_L | 3.38865569286848e-05 |
| 68 | 17155 | 0.004680959 | 12605 | ACH | IN03A010_T3_L | 0.00723345 | MNhl29 | | T3_L | 3.385948324009425e-05 |
| 69 | 17155 | 0.001619108 | 11848 | ACH | INXXX066_T3_L | 0.009099034 | Sternotrochanter MN | | T3_R | 3.3629129538370935e-05 |
| 70 | 17155 | 0.03117516 | 13064 | GABA | AN06B015_T3_L | 0.001025502 | Itm MN | | T1_R | 3.197017555775072e-05 |
| 71 | 17155 | 0.001727189 | 26488 | GLUT | IN08A037_T3_L | 0.018228676 | Fe reductor MN | | T3_L | 3.1484370796872426e-05 |
| 72 | 17155 | 0.001961027 | 28526 | ACH | IN01A025_T1_L | 0.012353329 | Tr extensor MN | | T1_R | 3.093890933673357e-05 |
| 73 | 17155 | 0.008842737 | 12160 | ACH | IN03A010_T2_L | 0.003479294 | MNml29 | | T2_L | 3.076648220821263e-05 |
| 74 | 17155 | 0.001411876 | 21322 | ACH | IN04B081_T1_L | 0.012339934 | Sternal anterior rotator MN | | T1_L | 2.8792377483396498e-05 |
| 75 | 17155 | 0.011840432 | 192710 | ACH | IN19A017_T1_L | 0.001229318 | MNml78 | | T2_L | 2.875884815822224e-05 |
| 76 | 17155 | 0.001363378 | 12007 | GABA | IN14B006_T3_L | 0.010024057 | Acc. ti flexor MN | | T3_R | 2.8650697863695093e-05 |
| 77 | 17155 | 0.001073569 | 20441 | ACH | IN04B081_T2_L | 0.016886081 | Sternal anterior rotator MN | | T2_L | 2.8269982200535887e-05 |
| 78 | 17155 | 0.001363378 | 12007 | GABA | IN14B006_T3_L | 0.008974455 | Sternal posterior rotator MN | | T3_R | 2.759050214889822e-05 |
| 79 | 17155 | 0.00184997 | 23103 | GABA | IN09A064_T2_L | 0.005600758 | Acc. ti flexor MN | | T2_L | 2.713773320663187e-05 |
| 80 | 17155 | 0.005431345 | 16549 | ACH | IN08B058_T3_L | 0.004926694 | MNml29 | | T2_R | 2.6758573616329668e-05 |
| 81 | 17155 | 0.005431345 | 16549 | ACH | IN08B058_T3_L | 0.004861989 | MNxm02 | | T3_R | 2.6407141758758516e-05 |
| 82 | 17155 | 0.007705312 | 12220 | GABA | IN09A011_A1_L | 0.003381426 | MNhl64 | | T3_L | 2.605494296357311e-05 |
| 83 | 17155 | 0.00541806 | 10340 | ACH | IN07B006_T1_L | 0.004417826 | MNhl29 | | T3_R | 2.393604646731731e-05 |
| 84 | 17155 | 0.014027067 | 14084 | ACH | INXXX468_T3_L | 0.001679712 | Itm MN | | T3_L | 2.3561440257415678e-05 |
| 85 | 17155 | 0.003140606 | 10156 | GABA | IN19A003_T2_L | 0.007426943 | Fe reductor MN | | T2_L | 2.3325102451630053e-05 |
| 86 | 17155 | 0.006661675 | 13246 | ACH | INXXX468_T1_L | 0.003490387 | Itm MN | | T1_L | 2.325182343561669e-05 |
| 87 | 17155 | 0.005431345 | 16549 | ACH | IN08B058_T3_L | 0.002533432 | Sternal posterior rotator MN | | T3_R | 2.3228422437671183e-05 |
| 88 | 17155 | 0.001337875 | 23763 | GLUT | IN08A048_T3_L | 0.008709106 | Sternotrochanter MN | | T3_L | 2.3041949890542605e-05 |
| 89 | 17155 | 0.011840432 | 192710 | ACH | IN19A017_T1_L | 0.001928904 | Pleural remotor/abductor MN | | T2_L | 2.2839059151862424e-05 |
| 90 | 17155 | 0.004680959 | 12605 | ACH | IN03A010_T3_L | 0.004851834 | Sternal adductor MN | | T3_L | 2.271123502471641e-05 |
| 91 | 17155 | 0.005431345 | 16549 | ACH | IN08B058_T3_L | 0.002161982 | Sternal posterior rotator MN | | T2_R | 2.240745892304894e-05 |
| 92 | 17155 | 0.001275614 | 21189 | ACH | IN04B081_T1_L | 0.011001371 | Sternal anterior rotator MN | | T1_L | 2.0439923325506762e-05 |
| 93 | 17155 | 0.011840432 | 192710 | ACH | IN19A017_T1_L | 0.001713666 | MNhl87 | | T3_L | 2.029054547167901e-05 |

Supplementary Table 2

| | | | | | | | | | |
|---|---|---|---|---|---|---|---|---|---|
| 94 | 17155 | 0.001282818 | 11303 | GABA | IN19A003_T3_L | 0.014521104 | Fe reductor MN | T3_L | 2.0210209767451345e-05 |
| 95 | 17155 | 0.001411876 | 21322 | ACH | IN04B081_T1_L | 0.005657476 | Tergopleural/Pleural promotor MN | T1_L | 2.000103387890802e-05 |
| 96 | 17155 | 0.003200476 | 13318 | ACH | IN20A.22A003_T2_l | 0.003783322 | Tr extensor MN | T2_L | 1.966428573901824e-05 |
| 97 | 17155 | 0.001363378 | 12007 | GABA | IN14B006_T3_L | 0.014047369 | Fe reductor MN | T3_R | 1.915187020358172e-05 |
| 98 | 17155 | 0.001452839 | 25287 | GABA | IN09A064_T3_L | 0.007236717 | Acc. ti flexor MN | T3_L | 1.9045036129868972e-05 |
| 99 | 17155 | 0.003200476 | 13318 | ACH | IN20A.22A003_T2_l | 0.005872362 | MNml79 | T2_L | 1.879435543876093e-05 |
| 100 | 17155 | 0.001619108 | 11848 | ACH | INXXX066_T3_L | 0.009478271 | Tr extensor MN | T3_R | 1.876246862248029e-05 |

Supplementary Table 3

| Statistics performed in Prism GraphPad | | | | | | | | | | | | |
|---|---|---|---|---|---|---|---|---|---|---|---|---|
| Figure panel | Method | Data sets | Mean 1 | Mean 2 | Mean Diff. | SE of diff. | n1 | n2 | t | DF | Summary | *p* value |
| Fig. 1d | Unpaired t test | LAL013 vs Empty | 208.4 | 36.93 | -171.5 | 9.467 | 44 | 39 | 18.12 | 81 | **** | <0.0001 |
| | One sample test | LAL013 vs 0 | 208.4 | 0 | 208.4 | 8.567 | 44 | | 24.33 | 43 | **** | <0.0001 |
| | One sample test | Empty vs 0 | 36.93 | 0 | 36.93 | 2.753 | 39 | | 13.41 | 38 | **** | <0.0001 |
| Figure panel | Method | Data sets | Mean 1 | Mean 2 | Mean Diff. | SE of diff. | n1 | n2 | t | DF | Summary | *p* value |
| Fig. 1e | Unpaired t test | LAL013 vs Empty | -2.236 | 1.399 | 3.636 | 0.6077 | 44 | 39 | 5.982 | 81 | **** | <0.0001 |
| | One sample test | LAL013 vs 0 | -2.236 | 0 | -2.236 | 0.3842 | 44 | | 5.82 | 43 | **** | <0.0001 |
| | One sample test | Empty vs 0 | 1.399 | 0 | 1.399 | 0.4784 | 39 | | 2.925 | 38 | ** | 0.0058 |
| Figure panel | Method | Data sets | Number of treatments | Number of values (total) | | | | F | R squared | | Summary | *p* value |
| Fig. 1p | Ordinary one-way ANOVA | Unilateral; No label; Bilateral | 3 | 87 | | | | 48 | 0.5333 | | **** | <0.0001 |
| LAL013 | Method | Data sets | Mean 1 | Mean 2 | Mean Diff. | SE of diff. | n1 | n2 | t | DF | Summary | Adjusted *p* value |
| | Holm-Sidak's multiple comparisons test | Unilateral vs. No label | 186.6 | -10.26 | 196.9 | 27.2 | 46 | 25 | 7.237 | 84 | **** | <0.0001 |
| | | Unilateral vs. Bilateral | 186.6 | -82.74 | 269.3 | 31.77 | 46 | 16 | 8.477 | 84 | **** | <0.0001 |
| | | No label vs. Bilateral | -10.26 | -82.74 | 72.48 | 35.05 | 25 | 16 | 2.068 | 84 | * | 0.0417 |
| Figure panel | Method | Data sets | Number of treatments | Number of values (total) | | | | F | R squared | | Summary | *p* value |
| Fig. 1s | Ordinary one-way ANOVA | Unilateral; No label; Bilateral | 3 | 39 | | | | 14 | 0.4394 | | **** | <0.0001 |
| LAL013 flight | Method | Data sets | Mean 1 | Mean 2 | Mean Diff. | SE of diff. | n1 | n2 | t | DF | Summary | Adjusted *p* value |
| | Holm-Sidak's multiple comparisons test | Unilateral vs. No label | 16.3 | 0.8556 | 15.45 | 3.622 | 18 | 14 | 4.265 | 36 | *** | 0.0003 |
| | | Unilateral vs. Bilateral | 16.3 | -3.887 | 20.19 | 4.528 | 18 | 7 | 4.459 | 36 | *** | 0.0002 |
| | | No label vs. Bilateral | 0.8556 | -3.887 | 4.743 | 4.705 | 14 | 7 | 1.008 | 36 | ns | 0.3202 |
| Figure panel | Method | Data sets | Number of treatments | Number of values (total) | | | | F | R squared | | Summary | *p* value |
| Fig.2c | Ordinary one-way ANOVA | Unilateral; No label; Bilateral | 3 | 54 | | | | 256 | 0.9094 | | **** | <0.0001 |
| DNa11 | Method | Data sets | Mean 1 | Mean 2 | Mean Diff. | SE of diff. | n1 | n2 | t | DF | Summary | Adjusted *p* value |
| | Holm-Sidak's multiple comparisons test | Unilateral vs. No label | 247 | -2.372 | 249.4 | 11.35 | 26 | 25 | 21.98 | 51 | **** | <0.0001 |
| | | Unilateral vs. Bilateral | 247 | -4.778 | 251.8 | 24.7 | 26 | 3 | 10.2 | 51 | **** | <0.0001 |
| | | No label vs. Bilateral | -2.372 | -4.778 | 2.406 | 24.75 | 25 | 3 | 0.0972 | 51 | ns | 0.9229 |
| | Method | Data sets | Number of treatments | Number of values (total) | | | | F | R squared | | Summary | *p* value |

Supplementary Table 3

| | | | | | | | | | | | | |
|---|---|---|---|---|---|---|---|---|---|---|---|---|
| DNae003 | Ordinary one-way ANOVA | Unilateral; No label; Bilateral | 3 | 50 | | | | 22 | 0.4889 | | **** | <0.0001 |
| | Method | Data sets | Mean 1 | Mean 2 | Mean Diff. | SE of diff. | n1 | n2 | t | DF | Summary | Adjusted *p* value |
| | Holm-Sidak's multiple comparisons test | Unilateral vs. No label | 46.79 | -6.149 | 52.94 | 8.265 | 22 | 23 | 6.405 | 47 | **** | <0.0001 |
| | | Unilateral vs. Bilateral | 46.79 | -6.162 | 52.95 | 13.73 | 22 | 5 | 3.856 | 47 | *** | 0.0007 |
| | | No label vs. Bilateral | -6.149 | -6.162 | 0.0126 | 13.68 | 23 | 5 | 0.0009 | 47 | ns | 0.9993 |
| | Method | Data sets | Number of treatments | Number of values (total) | | | | F | R squared | | Summary | *p* value |
| DNa03 | Ordinary one-way ANOVA | Unilateral; No label; Bilateral | 3 | 83 | | | | 37 | 0.4825 | | **** | <0.0001 |
| | Method | Data sets | Mean 1 | Mean 2 | Mean Diff. | SE of diff. | n1 | n2 | t | DF | Summary | Adjusted *p* value |
| | Holm-Sidak's multiple comparisons test | Unilateral vs. No label | 79.05 | -5.488 | 84.54 | 10.19 | 36 | 36 | 8.296 | 80 | **** | <0.0001 |
| | | Unilateral vs. Bilateral | 79.05 | 3.196 | 75.86 | 14.89 | 36 | 11 | 5.093 | 80 | **** | <0.0001 |
| | | No label vs. Bilateral | -5.488 | 3.196 | -8.684 | 14.89 | 36 | 11 | 0.5831 | 80 | ns | 0.5615 |
| | Method | Data sets | Number of treatments | Number of values (total) | | | | F | R squared | | Summary | *p* value |
| DNa02 | Ordinary one-way ANOVA | Unilateral; No label; Bilateral | 3 | 51 | | | | 37 | 0.4825 | | **** | <0.0001 |
| | Method | Data sets | Mean 1 | Mean 2 | Mean Diff. | SE of diff. | n1 | n2 | t | DF | Summary | Adjusted *p* value |
| | Holm-Sidak's multiple comparisons test | Unilateral vs. No label | 18.31 | -8.939 | 27.25 | 7.159 | 23 | 18 | 3.807 | 48 | ** | 0.0012 |
| | | Unilateral vs. Bilateral | 18.31 | -7.139 | 25.45 | 8.616 | 23 | 10 | 2.954 | 48 | ** | 0.0097 |
| | | No label vs. Bilateral | -8.939 | -7.139 | -1.799 | 8.972 | 18 | 10 | 0.2005 | 48 | ns | 0.8419 |
| Figure panel | Method | Data sets | Number of treatments | Number of values (total) | | | | F | R squared | | Summary | *p* value |
| Fig. 2d | Ordinary one-way ANOVA | Unilateral; No label; Bilateral | 3 | 41 | | | | 19 | 0.5029 | | **** | <0.0001 |
| DNa03 flight | Method | Data sets | Mean 1 | Mean 2 | Mean Diff. | SE of diff. | n1 | n2 | t | DF | Summary | Adjusted *p* value |
| | Holm-Sidak's multiple comparisons test | Unilateral vs. No label | 15.82 | -0.01927 | 15.84 | 2.57 | 18 | 19 | 6.162 | 38 | **** | <0.0001 |
| | | Unilateral vs. Bilateral | 15.82 | 4.875 | 10.95 | 4.32 | 18 | 4 | 2.534 | 38 | * | 0.0308 |
| | | No label vs. Bilateral | -0.01927 | 4.875 | -4.894 | 4.299 | 19 | 4 | 1.138 | 38 | ns | 0.2621 |
| Figure panel | Method | Data sets | Number of treatments | Number of values (total) | | | | F | R squared | | Summary | *p* value |
| Fig. 2g | Ordinary one-way ANOVA | Empty; LAL013; DNa11; DNa03; DNae003; DNa02 | 6 | 42 | | | | 18 | 0.7124 | | **** | <0.0001 |
| | Method | Data sets | Mean 1 | Mean 2 | Mean Diff. | SE of diff. | n1 | n2 | q | DF | Summary | Adjusted *p* value |
| | Dunnett's multiple comparisons test | Empty vs. LAL013 | 14.04 | -65.46 | 79.5 | 11.42 | 12 | 6 | 6.96 | 36 | **** | <0.0001 |

Supplementary Table 3

| | | | | | | | | | | | | |
|---|---|---|---|---|---|---|---|---|---|---|---|---|
| | | Empty vs.DNa11 | 14.04 | -66.84 | 80.88 | 11.42 | 12 | 6 | 7.081 | 36 | **** | <0.0001 |
| | | Empty vs. DNae003 | 14.04 | -26.44 | 40.48 | 11.42 | 12 | 6 | 3.544 | 36 | ** | 0.0053 |
| | | Empty vs. DNa03 | 14.04 | -1.055 | 15.1 | 11.42 | 12 | 6 | 1.322 | 36 | ns | 0.6032 |
| | | Empty vs. DNa02 | 14.04 | 8.628 | 5.412 | 11.42 | 12 | 6 | 0.4738 | 36 | ns | 0.9903 |
| Figure panel | Method | Data sets | Number of treatments | Number of values (total) | | | | | Exact or approximate | Kruskal-Wallis statistic | Summary | *p* value |
| Fig. 3p | Kruskal-Wallis test | Empty; DNa03; DNa11; DNa02 | 4 | 60 | | | | | Approximate | 16.16 | ** | 0.0011 |
| 90° menotaxis per fly | Method | Data sets | Mean rank 1 | Mean rank 2 | diff. | Z | n1 | n2 | | | Summary | Adjusted *p* value |
| | Dunn's multiple comparisons test | Empty SS vs. DNa03 | 15.33 | 38.2 | -22.87 | 3.587 | 15 | 15 | | | ** | 0.001 |
| | | Empty SS vs. VES008 | 15.33 | 31.87 | -16.53 | 2.593 | 15 | 15 | | | * | 0.0285 |
| | | Empty SS vs. DNa02 | 15.33 | 36.6 | -21.27 | 3.336 | 15 | 15 | | | ** | 0.0026 |
| Figure panel | Method | Data sets | Number of treatments | Number of values (total) | | | | | Exact or approximate | Kruskal-Wallis statistic | Summary | *p* value |
| Fig. 3q | Kruskal-Wallis test | Empty; DNa03; DNa11; DNa02 | 4 | 864 | | | | | Approximate | 97.6 | **** | <0.0001 |
| 90° menotaxis pooled | Method | Data sets | Mean rank 1 | Mean rank 2 | diff. | Z | n1 | n2 | | | Summary | Adjusted *p* value |
| | Dunn's multiple comparisons test | Empty SS vs. DNa03 | 310.3 | 515.4 | -205.1 | 8.882 | 268 | 206 | | | **** | <0.0001 |
| | | Empty SS vs. VES008 | 310.3 | 479.8 | -169.5 | 6.898 | 268 | 167 | | | **** | <0.0001 |
| | | Empty SS vs. DNa02 | 310.3 | 467.4 | -157.1 | 6.952 | 268 | 223 | | | **** | <0.0001 |
| **Statistics performed in MATLAB** | | | | | | | | | | | | |
| Figure panel | Method | Data sets | | tstat | df | sd | | n | | | Summary | *p* value |
| Fig. 1l | paired t-test | LAL013-SS1 Off vs On | | -3.6215 | 28 | 19.6004 | | 29 | | | ** | 0.0011 |
| | | LAL013-SS2 Off vs On | | -3.0285 | 29 | 23.3403 | | 30 | | | ** | 0.0051 |
| | | Empty Off vs On | | -0.4443 | 28 | 28.8379 | | 29 | | | ns | 0.6602 |
| Figure panel | Method | Data sets | | tstat | df | sd | | n | | | Summary | *p* value |
| Fig. 1m | paired t-test | LAL013-SS1 Off vs On | | -5.2823 | 25 | 24.5468 | | 26 | | | **** | 1.80E-05 |
| | | LAL013-SS2 Off vs On | | -3.3153 | 29 | 16.3891 | | 30 | | | ** | 0.0025 |
| | | Empty Off vs On | | -0.8092 | 28 | 24.6875 | | 29 | | | ns | 0.4252 |
| Figure panel | Method | Data sets | | tstat | df | sd | n1 | n2 | | | Summary | *p* value |
| Fig. 1u | Unpaired t test | ExR7 vs Empty | | 5.9384 | 52 | 12.6052 | 27 | 27 | | | **** | 2.41E-07 |
| | One sample test | ExR7 vs 0 | | 11.1672 | 26 | 14.9462 | 27 | | | | **** | 2.03E-11 |
| | One sample test | Empty vs 0 | | 6.2832 | 26 | 9.7157 | 27 | | | | **** | 1.19E-06 |
| Figure panel | Method | Data sets | | tstat | df | sd | n1 | n2 | | | Summary | *p* value |
| Fig. 1v | Unpaired t test | ExR7 vs Empty | | -4.2239 | 52 | 0.9202 | 27 | 27 | | | **** | 9.69E-05 |
| | One sample test | ExR7 vs 0 | | -4.1655 | 26 | 0.8682 | 27 | | | | *** | 0.0003 |
| | One sample test | Empty vs 0 | | 1.9395 | 26 | 0.9694 | 27 | | | | ns | 0.0634 |
| Figure panel | Method | Data sets | | tstat | df | sd | | n | | | Summary | *p* value |
| Fig. 2h | paired t-test | DNa03 Off vs On | | 3.2634 | 26 | 5.1583 | | 27 | | | ** | 0.0031 |
| | | DNa11 Off vs On | | 1.9532 | 37 | 4.4800 | | 38 | | | ns | 0.0584 |

Supplementary Table 3

| | | | | | | | | | | | | |
|---|---|---|---|---|---|---|---|---|---|---|---|---|
| | | LAL013 Off vs On | | 0.1935 | 21 | 5.5193 | | 22 | | | ns | 0.8484 |
| Figure panel | Method | Data sets | | r | mean (PVA amplitude) | mean (ExR7) | | n | | | Summary | *p* value (corr coef ≥ 0) |
| Fig. 4e | Pearson correlation test | EPG Mean 0.2-0.25 | | -0.2061 | 0.0803 | 0.2887 | | 39950 | | | **** | 0 |
| | | EPG Mean 0.25-0.3 | | -0.2876 | 0.1019 | 0.3369 | | 19775 | | | **** | 0 |
| | | EPG Mean 0.3-0.35 | | -0.3447 | 0.1271 | 0.3740 | | 8267 | | | **** | 1.4309E-229 |
| | | EPG Mean 0.35-0.4 | | -0.3120 | 0.1544 | 0.4054 | | 3283 | | | **** | 2.27824E-75 |
| | | EPG Mean 0.4-0.45 | | -0.3704 | 0.1865 | 0.4504 | | 1240 | | | **** | 6.52258E-42 |
| | | EPG Mean 0.45-0.5 | | -0.5170 | 0.2117 | 0.4782 | | 379 | | | **** | 1.3505E-27 |
| | | EPG Mean 0.5-0.55 | | -0.4271 | 0.2354 | 0.5033 | | 115 | | | **** | 9.70001E-07 |
| Figure panel | Method | Data sets | | tstat | df | sd | | n | | | Summary | *p* value |
| Fig. 5i | paired t-test | DNa03 vs DNa11 | | 3.0479 | 5 | 0.0884 | | 6 | | | * | 0.0285 |
| Figure panel | Method | Data sets | Source | SS | df | MS | F | | | | Summary | *p* value |
| Fig. S7d | 1-way ANOVA | Peak Correlation Coefficient | Groups | 0.0857 | 4 | 0.0214 | 2.2 | | | | ns | 0.0862 |
| | | | Error | 0.3160 | 33 | 0.0096 | | | | | | |
| | | | Total | 0.4017 | 37 | | | | | | | |
| Figure panel | Method | Data sets | Source | SS | df | MS | F | | | | Summary | *p* value |
| Fig. S7e | 1-way ANOVA | Lag at Peak Correlation | Groups | 0.9019 | 4 | 0.2255 | 24 | | | | **** | 1.95078E-09 |
| | | | Error | 0.3063 | 33 | 0.0093 | | | | | | |
| | | | Total | 1.2081 | 37 | | | | | | | |
| | Method | Data sets | 95% interval lower limit | mean | 95% interval higher limit | | n1 | n2 | | | Summary | *p* value |
| | Tukey-Kramer test | LAL013 vs DNa11 | 0.1242 | 0.2613 | 0.3984 | | 6 | 13 | | | **** | 4.01E-05 |
| | | LAL013 vs DNae003 | -0.0329 | 0.1275 | 0.2879 | | 6 | 6 | | | ns | 0.1729 |
| | | LAL013 vs DNa03 | 0.3123 | 0.4588 | 0.6052 | | 6 | 9 | | | **** | 1.18E-08 |
| | | LAL013 vs DNa02 | 0.1757 | 0.3550 | 0.5344 | | 6 | 4 | | | **** | 2.15E-05 |
| | | DNa11 vs DNae003 | -0.2709 | -0.1338 | 0.0033 | | 13 | 6 | | | ns | 0.0587 |
| | | DNa11 vs DNa03 | 0.0770 | 0.1975 | 0.3180 | | 13 | 9 | | | *** | 3.74E-04 |
| | | DNa11 vs DNa02 | -0.0652 | 0.0937 | 0.2526 | | 13 | 4 | | | ns | 0.4470 |
| | | DNae003 vs DNa03 | 0.1848 | 0.3313 | 0.4777 | | 6 | 9 | | | **** | 2.01E-06 |
| | | DNae003 vs DNs02 | 0.0482 | 0.2275 | 0.4069 | | 6 | 4 | | | ** | 0.0073 |
| | | DNa03 vs DNa02 | -0.2707 | -0.1037 | 0.0632 | | 9 | 4 | | | ns | 0.3951 |
| Figure panel | Method | Data sets | 95% interval lower limit | 95% interval higher limit | | | | n | | | Summary | *p* value (different from 0) |
| Fig. S9c | Bootstrap with replacement: 10000 iterations | LAL013 vs 0 | 0.043 | 0.15 | | | | 4 | | | * | <0.05 |
| | | DNa03 vs 0 | -0.018 | 0 | | | | 5 | | | ns | >=0.05 |
| **Statistics performed in Python: statsmodels and scipy** | | | | | | | | | | | | |
| Figure panel | Method | Data sets | U | | | | n1 | n2 | | | Summary | p value |
| Fig. 1f | Mann-Whitney U tests | LAL013 activtion vs. Control | 863 | | | | 30 | 30 | | | **** | 1.07E-09 |

Supplementary Table 3

| Figure panel | Method | Data sets | U | | | | n1 | n2 | p | | Summary | Adjusted *p* value |
|---|---|---|---|---|---|---|---|---|---|---|---|---|
| Fig. 1g | Mann-Whitney U tests with Bonferroni correction | LAL013-SS2 silencing vs. LAL013-SS2 Control | 134 | | | | 29 | 30 | 5.21E-06 | | **** | 1.56E-05 |
| | | LAL013-SS2 silencing vs. Empty Silencing | 134 | | | | 29 | 30 | 5.21E-06 | | **** | 1.56E-05 |
| | | LAL013-SS2 Control vs. Empty Silencing | 415 | | | | 30 | 30 | 0.6100 | | ns | 1 |
| Figure panel | Method | Data sets | U | | | | n1 | n2 | p | | Summary | Adjusted *p* value |
| Fig. 1h | Mann-Whitney U tests with Bonferroni correction | LAL013-SS1 silencing vs. LAL013-SS1 Control | 299 | | | | 27 | 36 | 0.0096 | | * | 0.0288 |
| | | LAL013-SS1 silencing vs. Empty Silencing | 252 | | | | 27 | 30 | 0.0148 | | * | 0.0444 |
| | | LAL013-SS1 Control vs. Empty Silencing | 554 | | | | 36 | 30 | 0.8620 | | ns | 1 |
| Figure panel | Method | Data sets | Variables | Trial No | | | n1 | n2 | | | Summary | *p* value |
| Fig. 4j-k; Fig. S6d-e | Wilcoxon rank sums tests | ExR7-SS1>Kir vs ExR7-SS1 Control | Tortuosity | Trial 1 | | | 19 | 19 | | | **** | 5.00E-05 |
| | | ExR7-SS1>Kir vs ExR7-SS1 Control | Path length | Trial 1 | | | 19 | 19 | | | ** | 0.0034 |
| | | ExR7-SS1>Kir vs ExR7-SS1 Control | Translational V | Trial 1 | | | 19 | 19 | | | ** | 0.0048 |
| | | ExR7-SS1>Kir vs ExR7-SS1 Control | Abs. Rotational V | Trial 1 | | | 19 | 19 | | | ** | 0.0028 |
| | | ExR7-SS1>Kir vs ExR7-SS1 Control | Tortuosity | Trial 2 | | | 20 | 19 | | | ** | 0.0010 |
| | | ExR7-SS1>Kir vs ExR7-SS1 Control | Path length | Trial 2 | | | 20 | 19 | | | ** | 0.0064 |
| | | ExR7-SS1>Kir vs ExR7-SS1 Control | Translational V | Trial 2 | | | 20 | 19 | | | ** | 0.0090 |
| | | ExR7-SS1>Kir vs ExR7-SS1 Control | Abs. Rotational V | Trial 2 | | | 20 | 19 | | | ** | 0.0032 |
| | | ExR7-SS2>Kir vs ExR7-SS2 Control | Tortuosity | Trial 1 | | | 20 | 20 | | | ** | 0.0087 |
| | | ExR7-SS2>Kir vs ExR7-SS2 Control | Path length | Trial 1 | | | 20 | 20 | | | ns | 0.8711 |
| | | ExR7-SS2>Kir vs ExR7-SS2 Control | Translational V | Trial 1 | | | 20 | 20 | | | ns | 0.7661 |
| | | ExR7-SS2>Kir vs ExR7-SS2 Control | Abs. Rotational V | Trial 1 | | | 20 | 20 | | | ns | 0.3867 |
| | | ExR7-SS2>Kir vs ExR7-SS2 Control | Tortuosity | Trial 2 | | | 19 | 20 | | | ** | 0.0059 |
| | | ExR7-SS2>Kir vs ExR7-SS2 Control | Path length | Trial 2 | | | 19 | 20 | | | ns | 0.8441 |

Supplementary Table 3

| | | | | | | | | | | | | |
|---|---|---|---|---|---|---|---|---|---|---|---|---|
| | | ExR7-SS2>Kir vs ExR7-SS2 Control | Translational V | Trial 2 | | | 19 | 20 | | | ns | 0.8004 |
| | | ExR7-SS2>Kir vs ExR7-SS2 Control | Abs. Rotational V | Trial 2 | | | 19 | 20 | | | ns | 0.7787 |
| Figure panel | Method | Data sets | Variables | Trial No | | | n1 | n2 | | | Summary | *p* value |
| Fig. 4l-m; Fig. S6f-g | Wilcoxon rank sums tests | LAL013-SS1>Kir vs LAL013-SS1 Control | Tortuosity | Trial 1 | | | 20 | 19 | | | * | 0.0145 |
| | | LAL013-SS1>Kir vs LAL013-SS1 Control | Path length | Trial 1 | | | 20 | 19 | | | ns | 0.0918 |
| | | LAL013-SS1>Kir vs LAL013-SS1 Control | Translational V | Trial 1 | | | 20 | 19 | | | ns | 0.1440 |
| | | LAL013-SS1>Kir vs LAL013-SS1 Control | Abs. Rotational V | Trial 1 | | | 20 | 19 | | | * | 0.0169 |
| | | LAL013-SS1>Kir vs LAL013-SS1 Control | Tortuosity | Trial 2 | | | 20 | 20 | | | * | 0.0102 |
| | | LAL013-SS1>Kir vs LAL013-SS1 Control | Path length | Trial 2 | | | 20 | 20 | | | ns | 0.8924 |
| | | LAL013-SS1>Kir vs LAL013-SS1 Control | Translational V | Trial 2 | | | 20 | 20 | | | ns | 0.7251 |
| | | LAL013-SS1>Kir vs LAL013-SS1 Control | Abs. Rotational V | Trial 2 | | | 20 | 20 | | | ns | 0.2036 |
| | | LAL013-SS2>Kir vs LAL013-SS2 Control | Tortuosity | Trial 1 | | | 20 | 20 | | | ** | 0.0038 |
| | | LAL013-SS2>Kir vs LAL013-SS2 Control | Path length | Trial 1 | | | 20 | 20 | | | ** | 0.0013 |
| | | LAL013-SS2>Kir vs LAL013-SS2 Control | Translational V | Trial 1 | | | 20 | 20 | | | ns | 0.0620 |
| | | LAL013-SS2>Kir vs LAL013-SS2 Control | Abs. Rotational V | Trial 1 | | | 20 | 20 | | | ** | 0.0068 |
| | | LAL013-SS2>Kir vs LAL013-SS2 Control | Tortuosity | Trial 2 | | | 18 | 20 | | | ** | 0.0060 |
| | | LAL013-SS2>Kir vs LAL013-SS2 Control | Path length | Trial 2 | | | 18 | 20 | | | ns | 0.1016 |
| | | LAL013-SS2>Kir vs LAL013-SS2 Control | Translational V | Trial 2 | | | 18 | 20 | | | ns | 0.8151 |
| | | LAL013-SS2>Kir vs LAL013-SS2 Control | Abs. Rotational V | Trial 2 | | | 18 | 20 | | | * | 0.0468 |
